# Control Cell Migration by Engineering Integrin Ligand Assembly

**DOI:** 10.1101/2021.09.14.460227

**Authors:** Xunwu Hu, Sona Rani Roy, Chengzhi Jin, Guanying Li, Qizheng Zhang, Asano Natsuko, Shunsuke Asahina, Tomoko Kajiwara, Bolu Feng, Kazuhiro Aoki, Atsushi Takahara, Ye Zhang

## Abstract

Advances in mechanistic understanding of integrin-mediated adhesion highlight the importance of precise control of ligand presentation in directing cell migration. Top-down nanopatterning limited the spatial presentation to sub-micron. To enhance it to molecular level, we propose a bottom-up nanofabrication strategy. Via self-assembly and co-assembly, precise control of ligand presentation is succeeded by varying the proportions of assembling ligand and nonfunctional peptide. Assembled nanofilaments fulfill multi-functions exerting enhancement to suppression effect on cell migration with tunable amplitudes. Self-assembled nanofilaments possessing super high ligand density selectively suppress cancer cell migration by preventing integrin/actin disassembly at cell rear, which provides new insights to ligand-density-dependent-modulation, revealing valuable inputs to therapeutic innovations in tumor metastasis.

**One-Sentence Summary:** Engineering integrin ligand assembly from bottom-up offers a generalized tool to selectively control cell migration with tunable amplitudes.

---

Cell migration plays a central role in a wide variety of biological phenomena, from embryogenesis to tumor metastasis, etc.(*1*) There is considerable interest in understanding cell migration on a molecular level because this could lead to novel therapeutic approaches in biotechnology.(*2*) Integrins, as the major family of cell receptors responsible for cell adhesion, have long served as the primary targets of biomaterials.(*3-5*) The development of top-down nanofabrication techniques(*6,7*) enhanced control over spatial presentation of integrin ligand, which promoted the mechanistic study of integrin-mediated adhesions to inspire biomaterial innovations. For example, fibronectin (50 μg/ml)-coated polystyrene microbeads (mean diameter 11.9 μm) facilitated the elucidation of synergic effects of integrin occupancy and aggregation on cellular response,(*8*) and RGD-functionalized Ti lithography nanopattern (10 nm lines with 40-490 nm distance) assisted the demonstration of ligand geometrical effects on adhesion cluster formation.(*9*) Enhancing spatial resolution beyond sub-micron for insights of subsequent cellular response will reveal design principles for future generations of biomaterials. To address the challenges, we develop a bottom-up fabrication strategy(*10*) by combining molecular self-assembly and co-assembly(*11*) for extracellular constructs.

As shown in Figure 1a, our strategy is to covalently connect non-functional assembling motif to ECM-derived integrin ligand synthesizing an assembling ligand. Self-assembly of the assembling ligand forms nanofilaments exhibiting super high ligand density. Via introducing the non-functional assembling motifs, co-assembled nanofilaments with precisely controlled ligand densities are produced by varying the proportion of the two components. To put our design into practice, repeats of _L_-phenylalanine (F, FF, and FFF) facilitating intermolecular aromatic interactions,(*12, 13*) were coupled to the N-terminus of IKLLI derived from laminin α1 chain(*14*) generating candidate assembling ligands. Preliminary evaluations indicated that FFFIKLLI preserved the conformation of IKLLI (Figure S1a, b), enhanced the integrin-binding affinity (Figure S1c-e), and selectively targeted integrin α_3_β_1_ (Figure S1f),(*15*) a molecular marker of malignant carcinomas, exerting suppression effects on cancer cell migration without inducing cytotoxicity (Figure S1g-i). Therefore, FFF and FFFIKLLI were selected as assembling motif and assembling ligand, respectively, for the proposed nanofabrication.

**Fig. 1.**
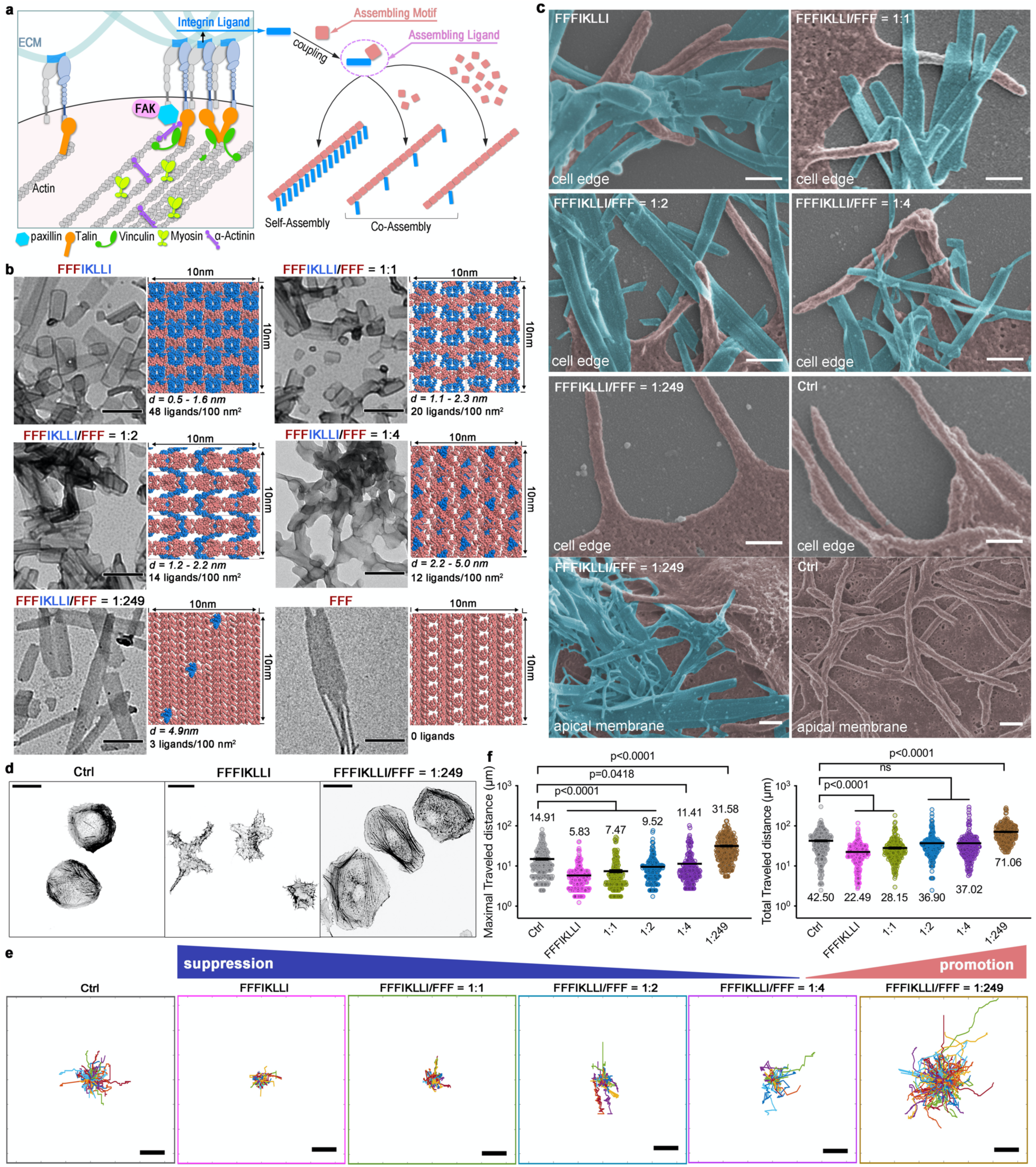
Engineering integrin ligand assembly to control cell migration. (**a**) Schematic illustration of precise control of integrin ligand presentation on nanofilaments via peptide assembly. (**b**) TEM images of nanofilaments obtained via molecular self-assembly and co-assembly of FFFIKLLI (100 μM) and FFF at various ratios, and the estimated molecular packing structures. IKLLI motif is presented in blue and FFF motif is presented in pink. The scale bars represent 200 nm. (**c**) Zoom-in SEM images (false color) of HuH-7 cell edge and apical membrane after 3-day incubations. FFFIKLLI was maintained at a concentration of 100 μM. Cell body is highlighted in pink, while the nanofilaments are highlighted in blue. The scale bars represent 300 nm. (**d**) The phalloidin staining of Huh-7 cells. Scale bars represent 20 μm. The trajectory plots (**e**), and the correlated quantitative analysis of maximal and total travel distance (**f**) of randomly selected migrating cells for each incubation condition. Kruskal-Wallis with Dunn’s multiple comparisons test was applied in data analysis. Error bars represent standard error of mean. n = 261, 280, 230, 260, 214, and 278 cells from left to right panel, respectively. Scale bars in panel **e** represent 50 μm.

Self-assembly of FFFIKLLI and co-assembly of FFFIKLLI with FFF at various ratios (Figure S2) all formed stable rectangular nanofilaments (∼100 nm width, ∼100-500 nm length) in water (Figure 1b, S3). Self-assembly of FFFIKLLI selectively targeted cancer cells expressing integrin α_3_β_1_, including HuH-7, HeLa, HepG2, A549, MKN1, and U-87 MG cells, to suppress their migration without inducing cytotoxicity (Figure S4). By comparison, it exerted negligible influence on cell lines lacking integrin α_3_β_1_ expression, for example MCF-7, Ect1/E6E7 cells (Figure S5a-c), and integrin α_3_β_1_-silenced HuH-7 cells (Figure S5d-g). Co-assembled nanofilaments attenuated the suppression effect on cell migration by raising the proportion of FFF and led to negligible influence when the two components reached 1 to 44 (FFFIKLLI/FFF) ratio (Figure S6). Intriguingly, at 1 to 89 ratio, co-assembled nanofilaments turned to an opposite function exerting promotion effect on cell migration. At the ratio of 1 to 249, more than 1.5 times faster cell migration was detected.

Quantitative estimation of surface ligand density of nanofilaments was conducted via molecular dynamics simulation based on the crystal structure of FFF unit cell,(*12*) followed by polymorph prediction. After the initial search, Fourier-transform infrared (FTIR) spectra of nanofilaments, which indicated the hydrogen bonding transition from N-H…N to N-H…O (*16*) due to the increasing proportion of FFF (Figure S7) were applied to select the adaptive packing modes (Figure S8). The polymorph predictions suggested that the molecular packing of self-assembled FFFIKLLI could expose 48 ligands/100 nm^2^, which is the highest record of ligand density, with the shortest distance between ligands ranging from 0.5 to 1.6 nm. Co-assembly of FFFIKLLI with FFF leads to decreased ligand density with increased ligand distance. For example, from 1 to 1, to 1 to 4, and to 1 to 249 ratio, the estimated ligand density decreased from 20 to 12 to 3 ligands per 100 nm^2^ with minimum ligand distance increased from 1.1 to 2.2 to 4.9 nm, respectively (Figure 1b, S9). Considering their influence on cell migration, we here categorize the ligand presentation on nanofilaments into four levels. Self-assembled FFFIKLLI possesses super high ligand density; co-assembled FFFIKLLI and FFF at 1 to 1 and 1 to 2 ratios possesses high ligand density; co-assembly at 1 to 4 ratio possesses intermediate ligand density; and co-assembly at 1 to 249 ratio possesses low ligand density.

Possessing different ligand density, nanofilaments demonstrated a variety of intimacy to the cell edge, especially the finger-like projections. SEM images of HuH-7 cells exhibited that nanofilaments with super high ligand density almost entangled with all peripheral projections. By reducing the ligand density, part of the cell edge had an intimate association with nanofilaments leaving more cell projections untouched. With low ligand density, the nanofilaments mainly attached to the apical membrane while the whole cell edge was untouched (Figure 1c, S10). Because cell migration is tightly associated with cell morphology, we next investigated the influence of nanofilaments on cell spreading (Figure 1d, S11) and characterized the correlated cell motilities (Figure 1e). HuH-7 cells were round and less spread on glass (Ctrl). Upon the treatment of nanofilaments with super high ligand densities, restricted HuH-7 cells exhibited reduced spreading area with tentacle-like actin extensions in all directions. By reducing the ligand density from high to intermediate level, HuH-7 cells gradually resumed the smooth cell edge correlated to their partially restored motility. Upon the treatment of nanofilaments with low ligand density, cells exhibited broad, flat lamellipodia correlated to almost 2 times enhancement on migration (Figure 1f).

Cell migration was defined as a four-step cycle: protrusion, adhesion, traction, and retraction, requiring spatiotemporal ordinations of the cytoskeleton and extracellular adhesion.(*17*) Therefore, to access the impact of nanofilaments on migration-correlated membrane dynamics, we performed a morphodynamical analysis by mapping the protrusion and retraction in response to various nanofilaments over time (Figure 2a, S12). The kymographs clearly indicated that nanofilaments with super high ligand density restricted the formation of protrusions. By reducing the ligand density to the intermediate level, the restriction effect was gradually attenuated. The quantitative analysis of protrusion and retraction velocity (Figure 2b, S13) showed that super high ligand density caused a global loss of the cluster periphery dynamics, while low ligand density promoted the formation of protrusion leading to enhanced cell migration.

**Fig. 2.**
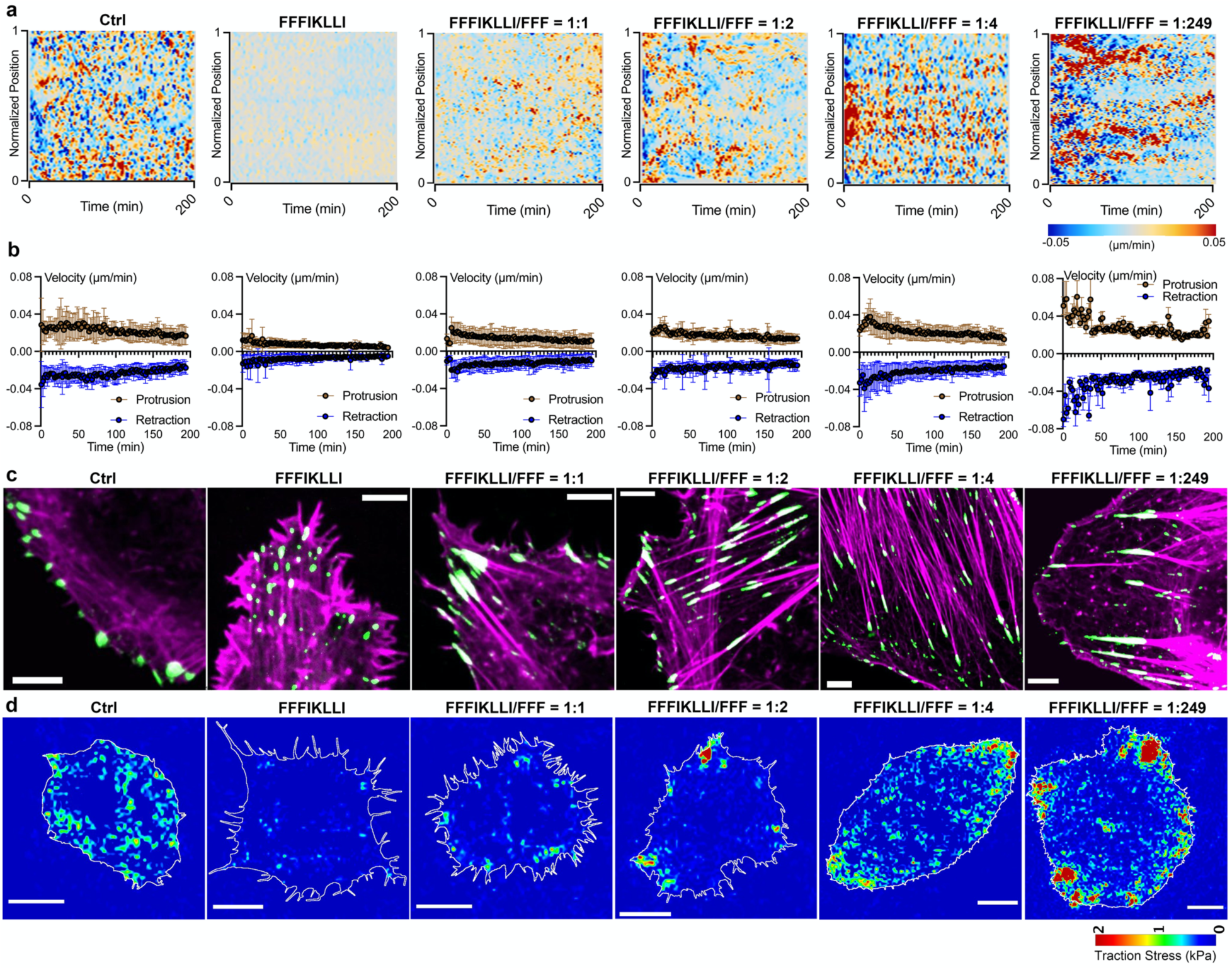
Nanofilaments regulate membrane dynamics and FA organizations in a ligand density dependent manner. Kymograph of normalized edge velocity (**a**) and mean velocity over time for protrusions and retractions of HuH-7 cells (**b**), merged F-actin phalloidin staining, and paxillin immunofluorescence in HuH-7 cells (**c**), heat-scale plots of traction stress magnitudes of HuH-7 cells (**d**), upon the treatment of various peptide assemblies for 12 hr. Scale bars in panel **c**, and **d** represent 5 μm and 20 μm, respectively.

We next studied integrin-mediated adhesion, which is a critical regulator of cell migration. Compared with control cells exhibiting peripheral FAs associated with peripheral actomyosin bundles, nanofilaments with super high ligand density caused vanishment of peripheral FAs but induced the formation of small dot-like FAs associated with tousled F-actin aggregates (Figure 2c, S14, S15). Compared with control, treated HuH-7 cells exhibited more than 60% reduction on traction stresses (Figure 2d and Figure S16). By reducing the ligand density, peripheral FAs associated with peripheral actomyosin bundles were formed again, together with the formation of larger streak-like FAs distributed on ventral cell surface connected with stress fibers. Particularly, at 1 to 2 ratio, the average traction stresses were resumed to control level, while the maximal traction force exerted at the cell periphery with few stress foci localized to the protrusion area increased dramatically. At the intermediate level, the ligand presentation was optimal to the formation of ventral stress fibers inducing double enhancement of average traction stresses. The bipolar distribution of stress foci localized on the two lateral cell extremities indicated the altered actin organization polarized the cell to guide its migration.(*18*) With low ligand density, nanofilaments triggered the formation of nascent adhesions (*19*) localized across the lamellipodia and FAs increased in size toward the convergence zone associated with F-actin bundles collected in a transverse band. Meanwhile, the traction force distribution exhibited a front-to-rear polarization with fused stress foci localized on lamellipodia corresponding to the distribution of FAs exerting 6 times higher retraction stresses compared with control, which indicated an enhanced cell migration.

In response to nanofilaments with high, intermediate, and low ligand densities, HuH-7 cells exhibited three phenotype FA organizations corresponding to the cell migration velocities that have been well demonstrated on biomaterials with different ligand concentrations.(*20*) However, super high ligand density, which triggered an interesting interplay between F-actin and FAs led to tentacle-like actin extensions on cell periphery, has never been well-presented nor illustrated. By tracking actin and FA dynamics via time-lapse imaging of mRuby-Lifeact and EGFP-paxillin co-transfected HuH-7 cells (Figure S17), we observed that while the stress-fiber-associated FAs slide inward, the actin cytoskeleton at the cell rear was not fully disassembled (Figure 3a). Different from the well-studied reduced trailing edge retraction caused by stable adhesion within the cell rear,(*21*) we only observed the co-localization of integrin α_3_β_1_ with talin and α-actinin remaining on the actin filaments at the cell rear while vinculin, paxillin, and FAK located in the inward-sliding FAs connected with stress fibers (Figure 3b, 3c, S18, S19). Such segmentation of FA complex which led to failed FA disassembly on cell edge is highly possible due to the excessive binding interaction between integrin α_3_β_1_ and ligands clustered in a super high density via self-assembly. Together with a great inhibition of Rac1 activity (Figure 3d) on cell periphery (Figure 3e) which indicated the prevention of both protrusion formation and forward motion, it was demonstrated that self-assembly of FFFIKLLI restricted both trailing edge retraction and leading-edge protrusion of HuH-7 cells resulting into depolarization suppressing cell motility.

**Fig. 3.**
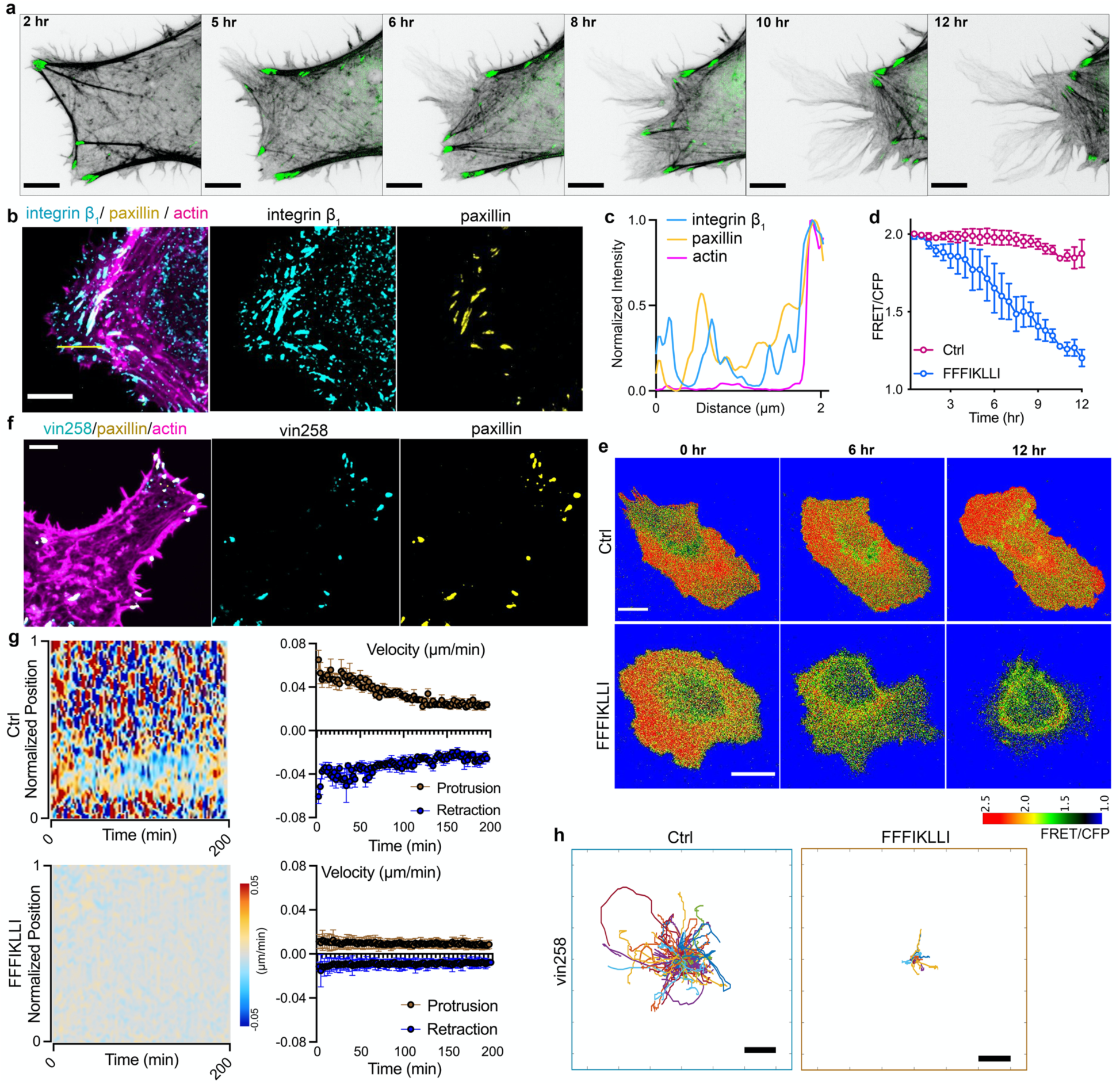
Nanofilaments with super high ligand density induced uncoordinated FA dynamics preventing trailing edge retraction. (**a**) Time-lapse series showing actin cytoskeleton (grey) and paxillin (green) in HuH-7 cells expressing mRuby-Lifeact-7 and pEGFPC1-mEGFP-paxillin upon the treatment of FFFIKLLI (100 μM) for 12 hr. Scale bars represent 2 μm. (**b**) F-actin phalloidin staining (magenta), integrin β1 (cyan) and paxillin (yellow) immunofluorescence in HuH-7 cell after 12 hr treatment of FFFIKLLI (100 μM). Scale bar represents 2 μm. (**c**) Fluorescence intensity distribution profile of integrin β1, paxillin, and actin cytoskeleton along the yellow line on merged image of panel **b**. (**d**) 12 hr time course Rac1 activity of HuH-7 cells with or without the treatment of FFFIKLLI (100 μM). Rac1 activity was measured by FRET. Symbols represent the mean FRET/CFP emission ratio ± s.d. (**e**) Representative FRET/CFP ratio images of HuH-7 cells expressing RaichuEV-Rac1 with or without the treatment of FFFIKLLI (100 μM) at the indicated time points and coded according to a pseudo color scale, which ranges from yellow to purple with an increase in Rac1 activity. Scale bars represent 20 μm. (**f**) F-actin phalloidin (magenta), vinculin N-terminal domain lacking the tail domain (vin258) (cyan), and paxillin (yellow) immunofluorescence in HuH-7 cell expressing pEFGPC1/GgVcL 1-258 upon the treatment of FFFIKLLI (100 μM) for 12 hr. Scale bar represents 5 μm. (**g**) Kymograph of normalized edge velocity and mean velocity over time for protrusions and retractions of HuH-7 cells expressing pEGFPC1/GgVcL 1-258 with or without the treatment of FFFIKLLI for 12 hr. (**h**) The trajectory plots of ∼200 randomly selected migrating HuH-7 cells expressing pEGFPC1/GgVcL 1-258 with or without the treatment of FFIKLLI. Scale bars represent 50 μm.

To understand the influence of super high ligand density on a molecular level, we did further exploration on the inside-out signaling to overcome the outside-in restriction. By expressing vin258, a mutant that possesses vinculin D1 domain exhibiting high affinity to talin and paxillin but lack of actin-binding domain,(*22*) in HuH-7 cells, FA complex maintained united on the periphery without connecting to stress fibers after 12 hr treatment of FFFIKLLI (Figure 3f, S20a). However, expressing the FA stabilizing forms of vinculin failed to preserve the actomyosin network, could not resume protrusion nor trailing edge retraction (Figure 3g, S20b-e). Eventually, the suppression effect on cell motility was remained (Figure 3h, S20f-g). To preserve FAs on cell edge and associated with stress fibers, we applied Rho Activator II on HuH-7 cells to drive elevated level of actomyosin contractility.(*23, 24*) Followed by 12 hr treatment of nanofilaments, the FAs remained on cell periphery associated with peripheral actin bundles (Figure 4a, S21a-d). Enhanced contractile forces eased the full disassembly of FAs facilitating trailing edge retraction, which was confirmed by the velocity profile of edge dynamics (Figure 4b). However, the protrusion activity was still restricted (Figure 4b) resulting into partial restoration of cell motility (Figure 4c, S21e-g). Because Rac1 can trigger new leading-edge formation by controlling local actin assembly and FA formation when activated at the cell edge,(*25*) we then activated Rac1 constantly by translocating Tiam1 to the plasma membrane of HuH-7 cells to compel the formation of lamellipodia (Figure S22). Upon the treatment of nanofilaments for 12 hr, cells exhibited both NAs localized across the lamellipodia, and FAs associated with F-actin bundles collected in a transverse band (Figure 4d, S23a). Both leading-edge protrusion and trailing edge retraction were not restricted (Figure 4e, S23b-f), same as the cell motility (Figure 4f). Together, the excessive binding interactions between integrins and the super high-density ligands deactivated the endogenous Rac1 effectively via an outside-in path. Besides reducing ligand density on nanofilaments extracellularly, constant activation of Rac1/Tiam signaling was the only effective rescue from in-side out. In regard to Rac1’s particular roles for tumor metastasis, the therapeutic potentials of super high ligand density are promising.

**Fig. 4.**
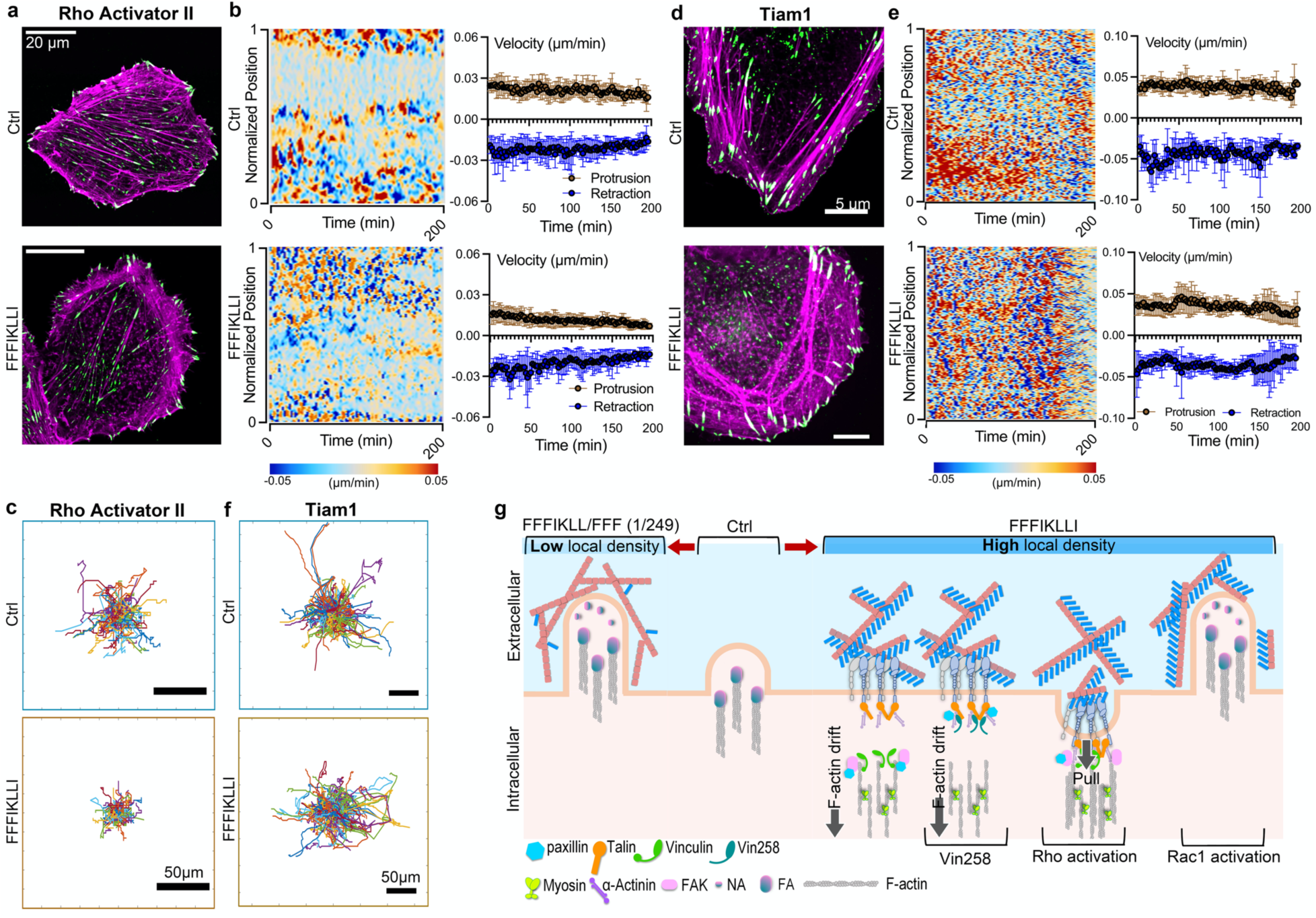
Overcome the influence of super high density of ligands via Rac1 signaling. F-actin phalloidin staining (magenta) and paxillin (green) immunofluorescence (**a**), kymograph of normalized edge velocity and mean velocity over time for protrusions and retractions (**b**), and the trajectory plots (n = ∼200) (**c**) of HuH-7 cells pretreated with Rho Activator II (1 μg/mL) with or without 12 hr treatment of FFFIKLLI (100 μM). F-actin phalloidin staining (magenta) and paxillin (green) immunofluorescence (**d**), kymograph of normalized edge velocity and mean velocity over time for protrusions and retractions (**e**), and the trajectory plots (n = ∼200) (**f**) of HuH-7 cells with elevated Rac1 activation with or without 12 hr treatment of FFFIKLLI (100 μM). (**g**) Schematic summary of outside-in regulation of HuH-7 cell motility via precise control of ligand density on nanofilaments, and the restoration of cell motility via inside-out signaling.

In summary, we developed a bottom-up nanofabrication technique for precise control of ligand presentation achieving by far the highest spatial resolution, which is beyond the submicron limitation of classic top-down techniques, and the highest surface ligand density in biomaterials without involving complex processing. We succeeded in producing uniformed nanofilaments with high to low ligand density via simple steps and achieved biphasic control of cancer cell motility from outside-in path (Figure 4g). The produced super high-density ligands established an excessive binding affinity with integrin α_1_β_3_ inducing segmentation of FA complex preventing trailing edge retraction, which selectively suppressed cancer cell migration. To promote a wide application of the technique, we randomly selected peptide sequences derived from ECM components possessing different binding interests to integrin isoforms for extracellular constructs.(*26, 27*) Fibronectin-derived GRGDSP, LRGDN, and synergy peptide PHSRN, laminin β1 chain-derived YIGSR,(*28*) and α1 chain-derived IKVAV, targeting integrin α_5_β_1_, α_v_β_3_, α_3_β_1_, or α_6_β_1_, were coupled with FFF obtaining a series of assembling ligands. Upon the treatment of these assembling ligands, cells phenocopied the morphology and motility of FFFIKLLI-treated cells indicating that the design strategy could be applied as generalized tool to fabricate biomaterials with therapeutic potentials in targeting subtype malignant tumor associated with different integrin expression pattern.

## Acknowledgments

The pEGFPC1-mEGFP-paxillin expression vector was generously provided by A. Kusumi Lab at Okinawa Institute of Science and Technology, Japan. The Rac1 FRET Sensor, pCAGGS-RaichuEV-Rac1 was generously provided by K. Aoki Lab at NIBB, Japan. The expression vectors for Rac1/Tiam1 activation system (Lyn11-linker-FB, YFP-FKBP, YFP-FKBP-linker-Tiam1, and pTriEx-PA-Rac1) were generously provided by T. Inoue at Johns Hopkins University, USA.

## Funding

This work was supported by

Takeda Science Foundation (2017 medical science) (YZ)

JSPS Grant-in-Aid for Scientific Research (B) 21H02063 (YZ)

OIST Proof-of-Concept (POC) Program 2019 (YZ)

## Author contributions

Conceptualization: XH, YZ

Methodology: XH, SRR, CJ, GL, QZ, AN, SA, TK, BF, KA, AT

Investigation: XH, YZ

Visualization: XH

Funding acquisition: YZ

Supervision: YZ

Writing – original draft: XH, YZ

Writing – review & editing: XH, KA, YZ

## Data and materials availability

All data are available in the main text or the supplementary materials.

## Supplementary Materials

Materials and Methods

Supplementary Text

Figs. S1 to S24

References (*1*–*9*)

