## Supporting Information for "Control Cell Migration by Engineering Integrin Ligand Assembly"

**This PDF file includes:**

Materials and Methods
Supplementary Text
Figs. S1 to S24
Appendix S1 to S13
References 1-14

### Materials and Methods

Synthesis

All the solvents and chemicals are commercially available. Chemicals were used without further purification: solvents were further purified by Ultimate Solvent System (Nikko Hansen, Japan) before use. $^{1}$H NMR spectra were recorded in deuterated solvent on a Bruker Advance 500 MHz spectrometer, and a JEOL JNM-ECZR 600 MHz spectrometer.

Five peptides (FFFIKLLI, FFFKLIIL, FFFGRGDSP, FFFLRGDN, FFFIKVAV), plus all Fmoc-amino and resin that used in the peptide synthesis, were purchased from GL Biochem (Shanghai) Ltd. China. Peptide FFF, IKLLI, FIKLLI, FFIKLLI, FFFPHSRN and FFFYIGSR were synthesized by following the Fmoc-based synthesis principle as described previously(*1,2*).

**FFF**

Purity: 99.51%

Molecular weight: 459.57. ESI-MS (m/z) $[M+H]^+$ = 460.15

$^{1}$H NMR (500 MHz, DMSO-$d_6$) $\delta$ 12.83 (s, 1H), 8.74 (d, $J$ = 7.5 Hz, 1H), 8.56 (d, $J$ = 7.8 Hz, 1H), 7.99 (s, 2H), 7.30 – 7.17 (m, 15H), 4.64 (td, $J$ = 8.7, 4.6 Hz, 1H), 4.50 (td, $J$ = 8.3, 5.3 Hz, 1H), 3.97 (s, 1H), 3.13 – 3.00 (m, 3H), 2.94 (dd, $J$ = 14.0, 8.8 Hz, 1H), 2.86 (dd, $J$ = 14.3, 8.5 Hz, 1H), 2.79 (dd, $J$ = 14.0, 9.3 Hz, 1H).

**FFIKLLI**

Purity: 98.94%

Molecular weight: 893.17. ESI-MS (m/z) $[M-H]^-$ = 891.56

**FFFIKLLI**

Purity: 98.32%

Molecular weight: 1040.44. ESI-MS (m/z) $[M+H]^+$ = 1040.60

$^{1}$H NMR (500 MHz, DMSO-$d_6$) $\delta$ 12.61 (1H, s), 8.72-8.65 (1H, m), 8.40 (1H, d, $J$ = 8.0 Hz), 8.07 (1H, d, $J$ = 8.8 Hz), 8.05 (1H, d, $J$ = 8.2 Hz), 8.01-7.94 (4H, m), 7.74 (1H, d, $J$ = 8.4 Hz), 7.66 (2H, s), 7.31-7.13 (15H, m), 4.72-4.66 (1H, m), 4.64-4.58 (1H, m), 4.40-4.22 (4H, m), 4.16 (1H, dd, $J$ = 8.3, 5.9 Hz), 3.97 (1H, s), 3.06-2.98 (3H, m), 2.87-2.77 (3H, m), 2.77-2.70 (2H, m), 1.81-1.69 (2H, m), 1.66-1.34 (12H, m), 1.34-1.24 (2H, m), 1.21-1.04 (2H, m), 0.91-0.75 (24H, m) ppm.

**FFFKLIIL**

Purity: 98.72%

Molecular weight: 1040.34. ESI-MS (m/z) $[M+H]^+$ = 1040.75

$^{1}$H NMR (500 MHz, DMSO-$d_6$) $\delta$ 12.48 (s, 1H), 8.68 (d, $J$ = 8.2 Hz, 1H), 8.40 (d, $J$ = 7.9 Hz, 1H), 8.21 (d, $J$ = 8.0 Hz, 1H), 8.12 (d, $J$ = 7.9 Hz, 1H), 8.05 – 7.97 (m, 3H), 7.93 (d, $J$ = 8.8 Hz, 1H), 7.81 – 7.73 (m, 3H), 7.29 – 7.13 (m, 15H), 4.66 – 4.58 (m, 2H), 4.37 (dd, $J$ = 15.1, 7.7 Hz, 1H), 4.31 (dd, $J$ = 13.8, 7.9 Hz, 1H), 4.25 – 4.16 (m, 3H), 3.97 (s, 1H), 3.08 – 2.99 (m, 3H), 2.88 – 2.70 (m, 5H), 1.75 – 1.36 (m, 12H), 1.35 – 1.25 (m, 2H), 1.05 (dt, $J$ = 13.6, 7.2 Hz, 2H), 0.91 – 0.74 (m, 24H).

**FFFGRGDSP**

Purity: 98.54%.

Molecular weight: 1029.20. ESI-MS (m/z) $[M+H]^+$ = 1029.40

$^1$H NMR (500 MHz, DMSO-$d_6$) $\delta$ 9.43 (s, 1H), 8.65 (s, 1H), 8.60 (s, 1H), 8.57 – 8.43 (m, 3H), 8.24 (t, $J$ = 5.4 Hz, 1H), 8.21 – 8.09 (m, 1H), 7.42 (s, 1H), 7.32 – 7.12 (m, 17H), 4.62 – 4.52 (m, 3H), 4.44 – 4.27 (m, 3H), 4.22 (dd, J = 8.7, 4.1 Hz, 1H), 3.86 – 3.72 (m, 4H), 3.66 – 3.51 (m, 3H), 3.50 – 3.32 (m, 2H), 3.20 – 3.10 (m, 1H), 3.09 – 2.94 (m, 5H), 2.86 (dd, $J$ = 13.9, 9.6 Hz, 1H), 2.82 – 2.70 (m, 2H), 2.57 (dd, $J$ = 16.5, 5.0 Hz, 1H), 2.15 – 2.03 (m, 1H), 1.89 – 1.77 (m, 2H), 1.75 – 1.40 (m, 4H).

**FFFLRGDN**
Purity: 98.73%.
Molecular weight: 1015.21. ESI-MS (m/z) $[M-H]^-$ = 1013.60
$^1$H NMR (500 MHz, DMSO-$d_6$) $\delta$ 12.46 (s, 2H), 8.69 (d, $J$ = 8.3 Hz, 1H), 8.41 (d, $J$ = 8.1 Hz, 1H), 8.24 (d, $J$ = 8.2 Hz, 1H), 8.21 (d, $J$ = 7.8 Hz, 1Hz), 8.17 – 8.14 (m, 1H), 8.11 (d, $J$ = 8.0 Hz, 1H), 8.02 (d, $J$ = 7.6 Hz, 1H), 7.98 (s, 2H), 7.59 (s, 1H), 7.40 (s, 1H), 7.31 – 7.17 (m, 15H), 6.93 (s, 1H), 4.65 – 4.59 (m, 1H), 4.51 – 4.46 (m, 1H), 4.44 – 4.28 (m, 2H), 4.13 (s, 2H), 3.97 (s, 1H), 3.82 – 3.67 (m, 2H), 3.62 (d, $J$ = 3.3 Hz, 1H), 3.59 (d, $J$ = 3.3 Hz, 1H), 3.55 (s, 1H), 3.53 (s, 1H), 3.48 – 3.42 (m, 2H), 3.05 – 2.99 (m, 2H), 2.88 – 2.75 (m, 3H), 2.68 (dd, $J$ = 16.7, 5.0 Hz, 1H), 1.75 – 1.60 (m, 2H), 1.58 – 1.43 (m, 5H), 0.89 (d, $J$ = 6.6 Hz, 3H), 0.86 (d, $J$ = 6.5 Hz, 3H).

**FFFIKVAV**
Purity: 98.70%.
Molecular weight: 970.21. ESI-MS (m/z) $[M+H]^+$ = 970.55
$^1$H NMR (500 MHz, DMSO-$d_6$) $\delta$ 12.64 (s, 1H), 8.70 (d, $J$ = 8.3 Hz, 1H), 8.41 (d, $J$ = 8.1 Hz, 1H), 8.13 – 8.07 (m, 2H), 8.03 (d, $J$ = 7.4 Hz, 1H), 7.99 (s, 2H), 7.88 (d, $J$ = 8.6 Hz, 1H), 7.77 (d, $J$ = 8.9 Hz, 1H), 7.72 (s, 2H), 7.31 – 7.15 (m, 15H), 4.69 (td, $J$ = 8.8, 4.3 Hz, 1H), 4.61 (td, $J$ = 8.6, 4.4 Hz, 1H), 4.41 – 4.36 (m, 1H), 4.36 – 4.30 (m, 1H), 4.28 – 4.23 (m, 1H), 4.19 (dd, $J$ = 8.8, 6.5 Hz, 1H), 4.14 (dd, $J$ = 8.6, 5.6 Hz, 1H), 3.97 (s, 1H), 3.06 – 2.99 (m, 3H), 2.87 – 2.71 (m, 5H), 2.05 (dq, $J$ = 13.5, 6.8 Hz, 1H), 1.96 (dq, $J$ = 13.5, 6.7 Hz, 1H), 1.78 – 1.70 (m, 1H), 1.70 – 1.62 (m, 1H), 1.58 – 1.40 (m, 4H), 1.37 – 1.26 (m, 2H), 1.19 (d, $J$ = 7.0 Hz, 3H), 1.13 – 1.06 (m, 1H), 0.89 – 0.78 (m, 18H).

Preparation of peptides assemblies

To prepare the stock solutions of peptide assemblies, required amount of peptide was dissolved in Milli-Q water at 100 mM (for FFF) or 10 mM (for the other peptides) and adjust the solution to reach a final pH of 7.0 using 1N NaOH solution. To prepare the working solution, the desired amount of stock solution was diluted using Milli-Q water or culture medium. Stand the working solutions for 30 min before applying them to the experiments.

Circular dichroism (CD) spectroscopy

CD spectra of the peptide assemblies in aqueous solutions were recorded under nitrogen atmosphere using a spectrophotometer JASCO J-820 (Jasco, Japan). Quartz cuvette with 0.5 mm path length was used as a sample container. Continuous scanning mode is applied. And each sample is scanned three times to obtain an averaged spectrum.

Fourier-transform infrared (FTIR) microscopy

To prepare FTIR samples, an aliquot of the peptide solution was dropped on a TS CaF2 window (5mm diameter) (Edmund Optics, Japan) and freeze-dried using lyophilizer (Freeze Dryer, Labconco). The CaF2 Window with assembled nanofilaments was mounted for FTIR microscopic analysis. FTIR microscopic analysis of a plain CaF2 Window served as the reference. An FTIR microscope (Nicolet iN10, Thermo Fisher Scientific) equipped with an MCT detector was used for collecting IR spectra. Spectra ranging from 4000 to 650 $cm^{-1}$ were obtained in transmission mode, aperture of 50 μm, with the resolution of 2 $cm^{-1}$. Each spectrum was derived from an average of 128 scans for each sample.

Transmission electron microscopy (TEM) imaging of peptide assemblies

To prepare TEM samples, 5 μL peptide solution was dropped on a glow-charged copper grid (400 mesh) coated with carbon film. After removing excess solution, the grid was gently washed three times to remove excess nanofilaments. Then the sample grid was negative stained using 1.0% (w/v) uranyl acetate and air-dried. TEM images were captured at a high vacuum using a transmission electron microscope (JEM-1230R, JEOL, Japan).

Molecular dynamics simulation and polymorph prediction

Molecular mechanics calculations were performed using Materials Studio. For all simulations, the Ewald method (*3,4*) was used for the electrostatic and van der Waals interaction terms. Gasteiger charges were used for an initial conformational search. As the crystal structure prediction method uses a rigid body approximation in the initial search for crystal packing alternatives, the analysis to determine low energy geometry was performed by following the protocols reported by Kim etc. (*5*), and the results were used as input for the packing calculations. The conformation of FFF was reported by Ellenbogen etc(*6*). Therefore, FFFIKLLI was drawn based on the structure of the FFF motif, and geometrical energy minimization scans were performed using the Forcite module of Materials Studio. After finding the lowest energy conformation of FFFIKLLI, the reported structure of the FFF unit cell was used as the starting point for crystal structure prediction using the Materials Studio Polymorph Predictor (PP). By replacing FFF by FFFIKLLI from the reported unit cell, PP calculation was performed.

The PP was set to its default fine setting (this sets the simulated annealing algorithm to a temperature range of 300-100000.0 K with a heating factor of 0.025, requiring 12 consecutive steps to be accepted before cooling and a maximum of 7000 steps) with the force field Dreiding 2.21 with Gasteiger charges. The 10 most common space groups found in organic crystals registered in the CSD were selected, including $P2_1/c$, $P1$, $P2_12_12_1$, $P2_1$, $C2/c$, $Pbca$, $Pna2_1$, $Pbcn$, $Cc$, and $C2$. Clustering of the predicted polymorphs was done using the polymorph clustering routine in Materials Studio. After the final clustering, hydrogen bonding analysis was performed on the calculated crystal structures to identify the packing modes matching the FTIR spectra regarding the hydrogen bonding signals. After extending the structure along the unit axes, the surface that exposes most integrin ligand IKLLI was presented in a defined square area aligned to the self-assembled nanostructures. The molecular packing of the mixture of FFFIKLLI and FFF at 1:249 ratio was predicted based on the crystal structure of FFF unit cell and the molecular packing structure of 1:4 ratio. The ligand distance was measured using the distance measurement functions of Materials Studio.

MicroScale Thermophoresis

The interaction between integrins and peptide ligands in PBS was quantified by a Monolith® NT.115Pico (NanoTemper Technology, Germany).(*7*) Recombinant human integrin proteins, including alpha-3/ beta-1, alpha-5/beta-1, and alpha-6/ beta-1, were purchased from R&D Systems (USA). Recombinant integrin proteins were labeled by Monolith$^{TM}$ Protein Labeling Kit RED-NHS (NanoTemper Technology) following the instruction provided by the manufacturer. The fluorescence-labeled integrin solution was mixed with peptides solution in 15 concentration gradients at a 1:1 ratio. The Monolith$^{TM}$ NT.115 MST Premium Coated Capillaries (NanoTemper Technology) were applied to draw the mixed solution with the order of declining ligand concentration. Pretest was performed to ensure the protein concentration and fluorescent intensities in each tube were equal. Following that, quantitative binding affinity was measured three times.

<u>Cell Culture and sample preparation</u>

Human hepatocellular carcinoma cell lines HuH-7 and Hep G2, human gastric adenocarcinoma cell line MKN1, human breast cancer cell line MCF-7 and human cervical cancer cell line HeLa were purchased from Riken BioResource Research Center. Human lung carcinoma cell line A549, human glioblastoma astrocytoma cell line U-87 MG, and human ectocervical cell line Ect1/E6E7 were purchased from American Type Culture Collection (ATCC).

Huh-7 and Hela were cultured in Dulbecco's Modified Eagle Medium (DMEM, Gibco) supplemented with 10% fetal bovine serum (FBS, Gibco), penicillin (100 µg/mL), and streptomycin (100µg/mL). MCF-7 was cultured in MEM (Gibco) with 10% FBS, penicillin (100 µg/mL), and streptomycin (100µg/mL). Hep G2 was cultured in Minimum Essential Media (MEM, Gibco) containing 10% FBS, Non-Essential Amino Acids (0.1 mm, Gibco), penicillin (100 µg/mL), and streptomycin (100µg/mL). U-87 MG was cultured in Eagle's Minimum Essential Medium (EMEM, ATCC) containing 10% FBS, penicillin (100 µg/mL), and streptomycin (100 µg/mL). MKN1 was cultured in RPMI 1640 medium (Gibco) containing 10% FBS, penicillin (100 µg/mL), and streptomycin (100µg/mL). A549 cell line was maintained in Ham's F-12K (Kaighn's) medium (Gibco) with 10% FBS, penicillin (100 µg/mL), and streptomycin (100µg/mL). Ect1/E6E7 was cultured in Keratinocyte-Serum Free medium (Gibco) with 0.1 ng/ml human recombinant EGF, 0.05 mg/ml bovine pituitary extract, and additional calcium chloride 44.1 mg/L (final concentration 0.4 mM).

For peptide treatment, stock solutions of peptide or the equivalent volume of DPBS were diluted in the culture medium contained low FBS (0.5%) and aged for 30 min before applying to the cells.

For Rho activation, cells were treated with 1 µg/ml Rho Activator II (#CN03-A, Cytoskeleton, Inc.) for 4 hr before applying to additional manipulations.

For transfection, cells were transfected by electroporation with 2.5-5 µg of endotoxin-free plasmid DNA per ~1 × 106 cells using the 4D-Nucleofector (Lonza) according to the manufacturer's instruction, except were indicated otherwise.

<u>Scanning electron microscopy (SEM) imaging of peptide assemblies treated HuH-7 cells</u>

For the preparation of the SEM samples, HuH-7 cells in the exponential growth phase were seeded on a 35 mm glass bottom petri dish. Once the cells were fully attached, peptide assemblies were added to the cell culture. After 12h incubation, the culture medium was removed, and the cell culture was washed three times using 1xPBS buffer. 2.5% glutaric dialdehyde was added to the cells for 30min followed by 1% osmium to fix the cells. Then the cells were dehydrated using ethanol, rinsed with t-BuOH, and freeze-dried in a lyophilizer (Freeze Dryer, Labconco) for more than 12 hr. All samples were coated with 5nm Osmium (Os) using Os coating device OPC80T (Filgen) before imaging. And the SEM images were captured using an ultra-high-resolution FE-SEM JSM-IT800SHL (JEOL, Japan) at 1.0 kV WD 6mm.

3-(4,5)-dimethylthiahiazo (-z-y1)-3, 5- diphenylte- trazolium bromide (MTT) assay

Huh-7 cells were seeded in 96-well plates at a density of $5x10^3$ cells/ well. The cells were allowed to adhere by incubating at 37 °C with 5% $CO_2$ for 12 hr. 100 µL of Cultured medium containing 10% FBS and the desired concentration of peptide replaced to each well and the cells were then incubated with the peptide solution for another 24, 48, or 72 hr. After the desired time of incubation, 10 µL of MTT (12 mM, Invitrogen) solution at a concentration of 5 mg/mL was added to each well and incubated at 37 °C for another 4 hr. The reduction reaction was terminated by adding 100 µL of 10% of SDS solution (in Milli-Q) and incubated for 12 hr to dissolve the formazan crystals formed in the cells. A Nivo multimode plate reader (PerkinElmer) was used to measure the optical density at the 570 nm wavelength of each well. All experiments were conducted in triplicate, and the results were calculated as mean ± standard deviation and presented as cell viability.

Wound healing assay

Healthy cells were seeded in 96-well Image Lock plates (Essen Bioscience, UK). Each well contained 2 $x10^4$ to $5x10^4$ cells in 100 µL of complete cell culture medium. Cell monolayer with approximately 90% confluence formed after 12 hr and cells were next starved in serum-free culture medium for 12 hrs. Homogenous scratch wounds with a width of about 700 µm were made by an Essen BioScienceWound Maker. The detached cells were washed with Dulbecco's Buffered Saline (DPBS, Gibco) 3 times. Cells were incubated with 100 µL of culture medium containing 0.5 % FBS and desired concentration of each peptide. Wound closure was monitored by an IncucyteS3 (Essen Bioscience) with a 10x objective. Images for each well were acquired every 2 hr to 8 hr for 2 days or until the wounds were closed completely. The wound healing rate was quantified by Incucyte Scratch Wound Analysis Module (Essen Bioscience). The experiments were repeated at least 3 times, and the results were presented as mean ± standard deviation of at least 3 independent experiments.

Western blotting

Protein lysates were prepared in ice-cold RIPA lysis, and extraction buffer (Thermo Scientific) supplemented with 1% Halt Protease Inhibitor (Thermo Scientific) and 1% Halt Phosphatase inhibitor (Thermo Scientific). Cells were scraped off from the culture surface using a cold plastic cell scraper, and the cell suspension was then gently transferred into a pre-cooled microcentrifuge tube and maintained on ice for 30 min with constant agitation before centrifuged at 15,000 rpm for 20 min. The supernatant was then transferred to a new tube, and the protein concentration of the supernatant was quantified according to the Pierce™ BCA Protein Assay Kit (Thermo Scientific). The rest of the supernatant was mixed with 4x laemmli sample buffer (Bio-Rad) in a 1:3 ration and boiled at 100°C for 10 min. Lysates can be aliquoted and stored at -80 °C freezer. When running the gel, 30 µg of protein was loaded into each well of the SDS-PAGE gels, along with a molecular weight marker (Bio-Rad). The gel was run for 2 hr at constantly 100 V to separate the proteins based on the molecular weight. The proteins were then transferred from the SDS-gel to a PVDF membrane (Bio-Rad) using a Transfer-blot turbo system (Bio-Rad). Once the membranes were blocked with Blocking One P solution (Nacalai) for 20 min at room temperature with gentle shaking, they were probed with antibodies against GAPDH (6C5, Abcam), integrin beta 1 (12G10, Abcam), integrin alpha 3 (ASC-1, Invitrogen), integrin alpha 6 (EPR18124, Abcam) diluted in Tris-Buffered Saline containing 0.1% Tween-20 and 5% Blocking one P solution and the membrane was incubated in the dark for overnight at 4 $^{o}$C with gentle shaking. The unbounded antibodies were washed out with tris-buffered saline (TBS) containing 0.1% Tween-20 (TBS- T) for 10 min for at least 6 times. The membranes were further incubated with secondary antibody conjugated with horseradish peroxidase (goat anti-mouse: #G-21040, goat anti-rabbit: #31460, Invitrogen) for 1 hr at room temperature. The membranes were rewashed with TBS-T for 10 min for at least 6 times. The protein bands were visualized according to the Chemiluminescence ECL detection kit (Bio-Rad) with a Fujifilm/GE LAS-3000. Protein bands were quantified by measuring peak areas using ImageJ. The peak area for each protein was normalized against the peak area of the loading control.

Integrin knockdown constructs and fluorescent protein fusion constructs

Integrin β1 shRNA plasmid (#sc-29375-SH), Integrin α3 shRNA plasmid (#sc-35684-SH), Integrin α6 shRNA plasmid (#sc-43129-SH), and control shRNA plasmid-A (#sc-108060) were purchased from Santa Cruz Biotechnology for knockdown of the target integrins. The knockdown efficiency was evaluated by western blotting.

The pEGFPC1-mEGFP-paxillin expression vector was a gift from A. Kusumi (Okinawa Institute of Science and Technology, Japan). mRuby-Lifeact-7 was a gift from Michael Davidson (RRID: Addgene_54560). pEGFPC1/Gg Vinculin 1-258 (akaVD1) was a gift from S. Craig (RRID: Addgene_46270)(*8*). The Rac1 FRET Sensor, pCAGGS-RaichuEV-Rac1was generously provided by K. Aoki (National Institute for Basic Biology, Japan)(*9*). The expression vectors for Rac1/Tiam1 activation system (Lyn11-linker-FRB, YFP-FKBP, YFP-FKBP-linker-Tiam1, and pTriEx-PA-Rac1) was generously provided by T. Inoue (Johns Hopkins University, USA)(*10*).

Confocal Microscopy and image analysis

All microscope imaging was performed with a Zeiss LSM 780 or Olympus SD-OSR. To assess the F-actin organization after certain treatment, cells were seeded on a glass-bottom culture dish (D11130H, Matsunami). After culture with the peptides, cells were fixed with 4% paraformaldehyde phosphate buffer solution (PFA, 30525-89-4, Wako) for 10 min at room temperature, washed, and permeabilized with 0.1% Triton X-100 (Sigma) in PBS for 15 minutes. Cells were washed with DPBS twice and incubated with ActinRed (Rhodamine-conjugated phalloidin, R37112, Invitrogen) or ActinGreen (Alexa-Fluor-488-conjugated phalloidin, R37110, Invitrogen) for 15 min at room temperature. Cells were washed with DPBS before being imaged. Morphological characteristics, including cell spreading area and perimeter area ratio, were quantified by Image J.

For live-cell time-lapse imaging, transfected HuH-7 cells were placed inside a stage-top incubator (Tokai Hit) and were tracked with a 100x/1.35 Silicon UPlanSApo objective for more than 12 hr. 5.5-μm stack images were acquired every 20 minutes, and imaging parameters were adjusted to minimize photobleaching and avoid cell death.

For tracking the cell migration, HuH-7 cells were labeled with fluorescent protein fusion constructs or Hoechst 33342. After being treated with or without the peptides for 12 hr, the cells were tracked with a 20x/0.8 Plan-Apochromat objective for 6 hr. The centroids of labeled cells were tracked using the TrackMate plugin (https://imagej.net/plugins/trackmate/) in ImageJ(*11*). A custom MATLAB script generated by Mr. B. Feng was applied to reconstruct the cell migration trajectory.

Immunocytochemistry

After 4% PFA fixation, cells were blocked by 5% bovine serum albumin (BSA, A7906, Sigma) in DPBS containing 0.1% Triton X-100 for 1 hr, followed by the incubation of primary antibodies against integrin beta 1 (12G10, Abcam), paxillin (Y113, Abcam), CD49c (integrin alpha 3, ASC-1, Invitrogen), talin 1 (8D4, Abcam), vinculin (EPR8185, Abcam), FAK (#3285, Cell signaling Technology), α-actinin (H-2, Santa Cruz Biotechnology), and Phospho-Myosin Light Chain 2 (Thr18/Ser19) (pMLC, #3674, Cell Signaling Technology) in 1% BSA for overnight at 4 $^{o}$C. The samples were washed twice with PBS before applying fluorescence-conjugated secondary antibodies, including Anti-Rabbit (Alexa Fluor 488, ab150077; Alexa Fluor 568, ab175471; and Alexa Fluor 647, ab150075; Abcam) and Anti-mouse (Alexa Fluor 488, ab150113; Alexa Fluor 568, ab175473; and Alexa Fluor 647, ab150115; Abcam), in 1% BSA with or without dye-conjugated phalloidin and DAPI (R37606, Invitrogen) at room temperature for 45 min.

Morphodynamics analysis

Morphodynamics maps were generated using the open-source ImageJ plugin "ADAPT"(*12*). The analyses were done on the maximum intensity projection of 5.5-µm stack images of mRuby-Lifeact-7 transfected HuH-7 cells pretreated with peptides for 12 hr. A spinning-disk confocal (Olympus SD-OSR) equipped with the camera Prime BSI sCMOS camera and a 100×/1.35 Silicon UPlanSApo objective was used to obtain the images every 2 minutes. All velocity values are directly measured using the plugin and plotted using GraphPad Prism. For a better visual representation, the map images (around 10000×100 pixels) were stretched to optimize visualization.

Traction Stress analysis

mRuby-Lifeact-7 transfected HuH-7 cells were seeded on a PDMS substrate functionalized with 0.2 µm FluoSpheres™ Carboxylate-Modified Microspheres (F8807, Invitrogen) at a density of approximately 6000 cells / cm$^2$. After being pretreated with peptides for 12 hr, the samples were imaged by a 20x/0.8 Plan-Apochromat objective, and the bead displacement obtained from confocal imaging was converted into force-displacement fields following established protocol(*13, 14*). Briefly, Images of beads with and without cell attachment were first aligned to correct experimental drift using ImageJ plugin "align slices in stack". The displacement field was subsequently calculated by another ImageJ plugin "PIV (Particle Image Velocimetry)". The cross-correlation PIV with 64 ×32-pixel size was used on all images for the PIV analysis to produce the position and vector field of the bead displacement. With the displacement field obtained from the PIV analysis, the traction force field was then reconstructed by the ImageJ plugin "FTTC".

Statistics and reproducibility

No statistical methods were used to predetermine sample size. The experiments were not randomized, and the investigators were not blinded to allocation during experiments and outcome assessment. All measurements were performed on 1-3 biological replicates from separate experiments. The exact sample size and exact statistical test performed for each experiment are indicated in the appropriate figure legends. Statistical analyses were performed using GraphPad Prism (GraphPad Software, www.graphpad.com). All bar graphs show mean values with error bars (s.e.m. or s.d., as defined in legends). The reported *P* values were corrected for multiple comparisons, where appropriate. Precise *P* values are shown in the figures and, when appropriate, are rounded to the nearest single significant digit. *P* values less than 0.0001 maybe be provided as a range. *P* values less than 0.05 are considered to be significant.

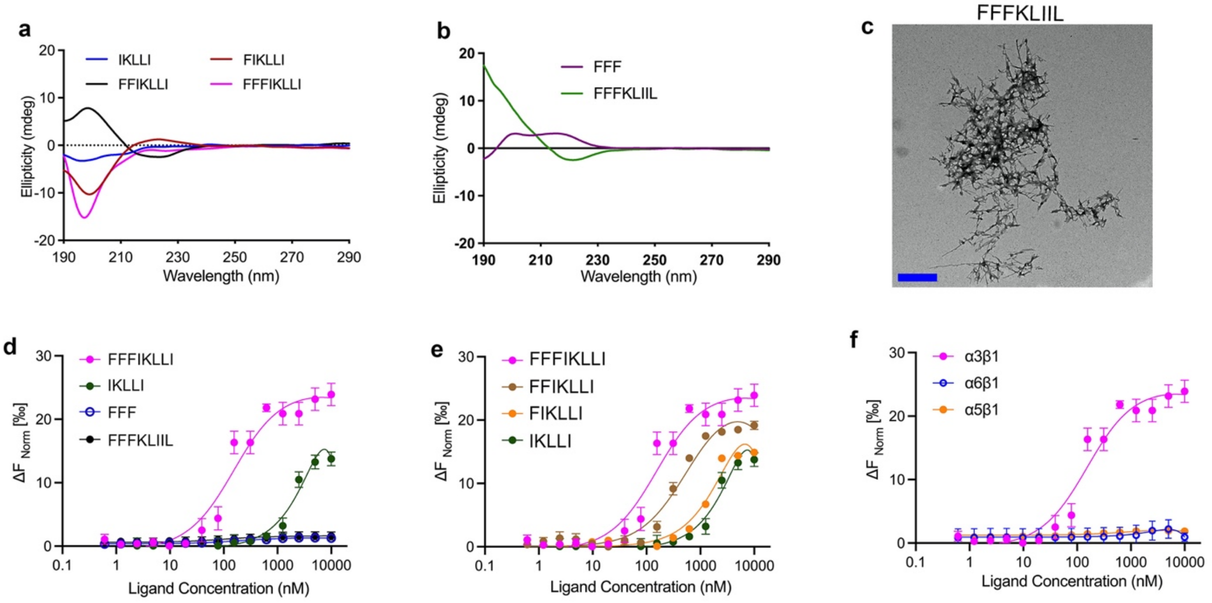


**Fig. S1.**

(**a**) Circular dichroism (CD) spectra of IKLLI, FIKLLI, FFIKLLI, FFFIKLLI at 100 μM concentration. (**b**) CD spectra of FFFKLIIL and FFF at 100 μM concentration. (**c**) TEM images of self-assembly of FFFKLIIL (scrambled peptide sequence was synthesized as negative control) in water at 100 μM concentration. Scale bar represents 500 nm. (**d**) The binding affinity of FFFIKLLI, IKLLI, FFF and FFFKLIIL to integrin α3β1 measured by MicroScale Thermophoresis. Protein concentration, 2nM. (**e**) The binding affinity of FFFIKLLI, FFIKLLI, FIKLLI and KLIIL to integrin α3β1 measured by MicroScale Thermophoresis. Protein concentration, 2nM. (**f**) The binding affinity of FFFIKLLI to integrin α3β1, α5β1and α6β1 measured by MicroScale Thermophoresis. Protein concentration, 2 nM. Error bars represent standard deviation (s.d.).

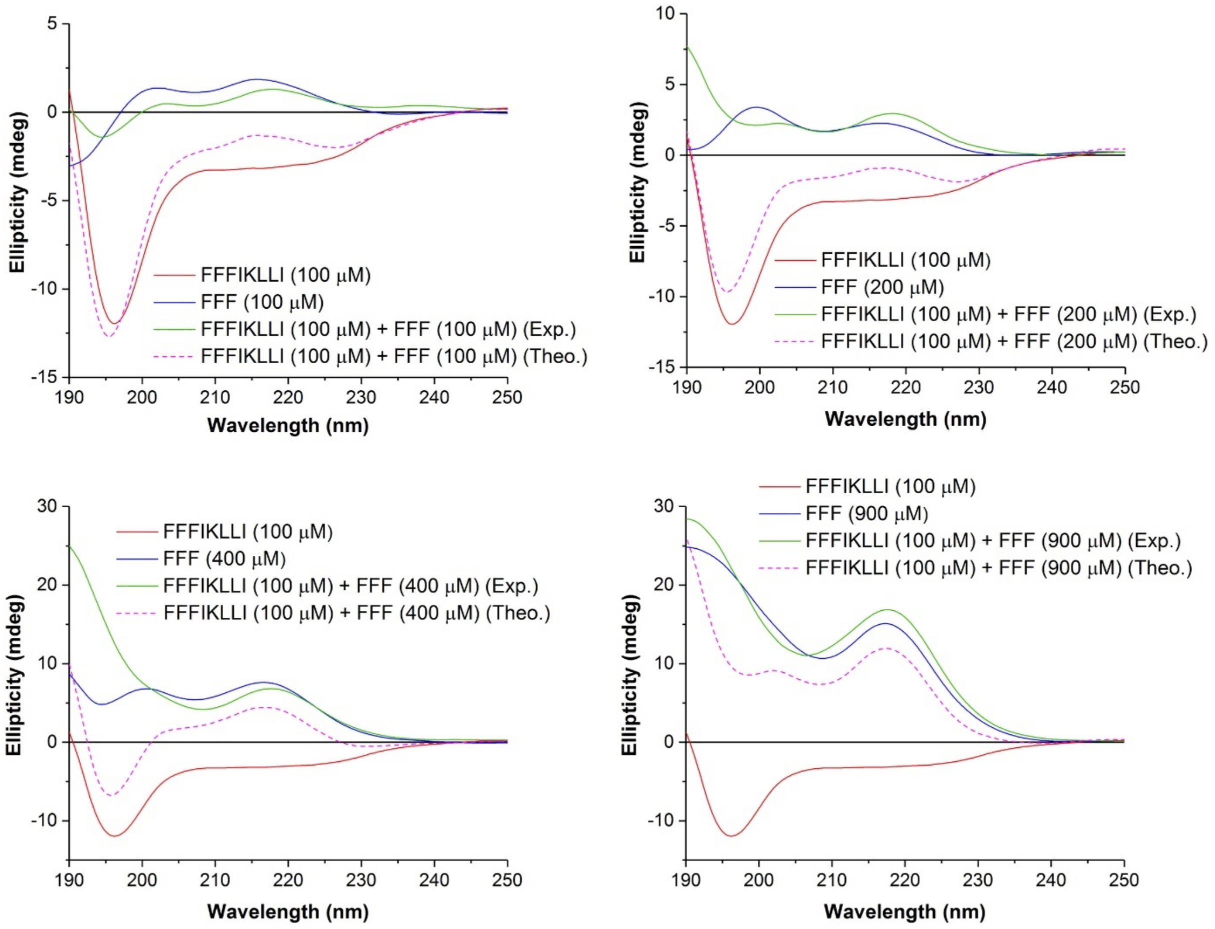


**Fig. S2.**

CD spectra of FFFIKLLI (100 μM), FFF (100, 200, 400, and 900 μM), and their mixtures. Exp. represents the experimental spectra of the mixture, while Theo. represents the simple sum of single component CD spectra of FFFIKLLI and FFF.

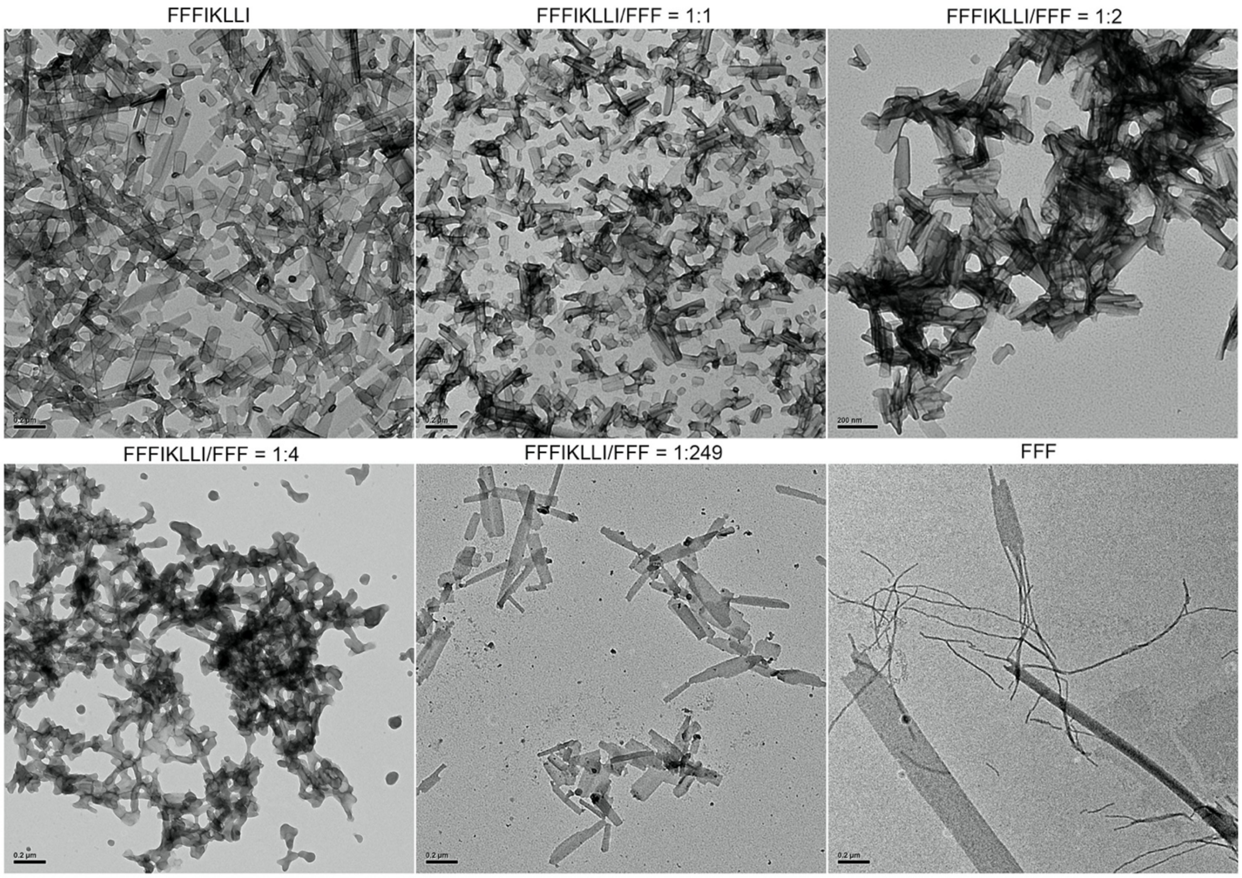


**Fig. S3.**
TEM images of nanofilaments formed via peptide self-assembly or co-assembly in water. The concentration of FFFIKLLI is kept 100 μM in every mixture. Self-assembly of FFF was imaged in solution of 100 μM FFF. Scale bar, 200 nm.

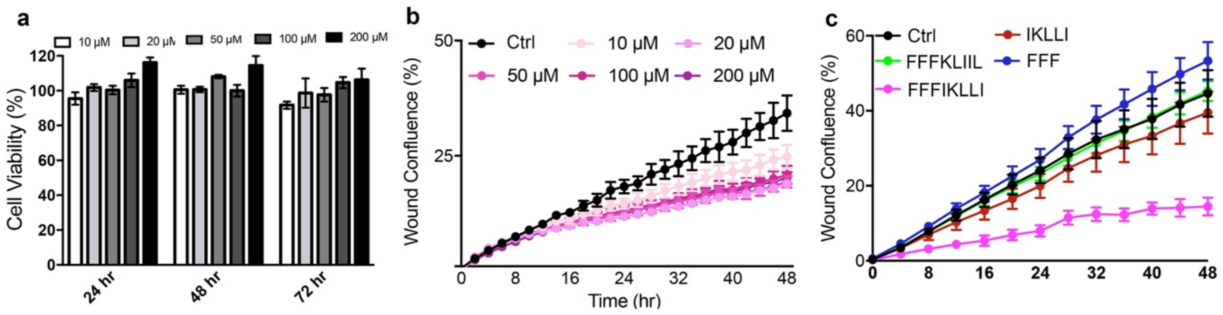


**Fig. S4.**

(**a**) HuH-7 cell viability using MTT assay. Cells were incubated with 10, 20, 50, 100, 200 μM of FFFIKLLI for 24, 48, or 72 hr. Error bars represent s.d.. (**b**) Wound healing rate of HuH-7 cells with or without the treatment of FFFIKLLI at 10, 20, 50, 100, and 200 μM. Error bars represent s.d.. (**c**) Wound healing rate of HuH-7 cells with or without the treatment of FFF, IKLLI, FFFIKLLI, and FFFKLIIL at 100 μM. Error bars represent s.d..

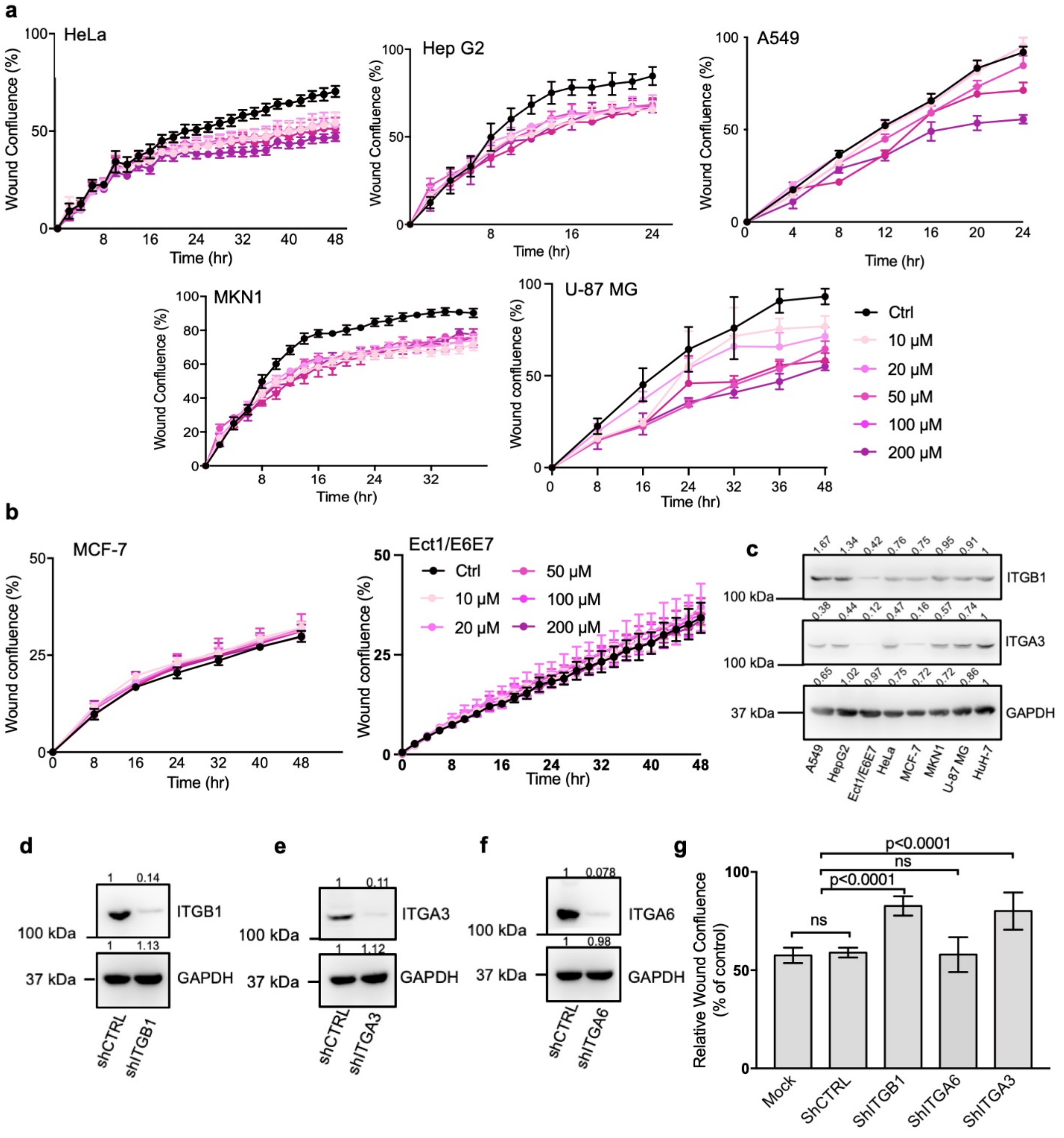


**Fig. S5.**

**(a):** Wound healing rate of peptide FFFIKLLI treatment on HeLa, Hep G2, A549, MKN1 and U-87MG cell lines. Error bars represent s.d.. **(b):** Wound healing rate of peptide FFFIKLLI treatment on MCF-7 and Ect1/E6E7 cell lines. Error bars represent s.d.. **(c):** Protein expression of integrin β1 and integrin α3 in different cells. **(d-f):** Knockdown efficiency of integrins in HuH-7. **(g):** 48 hr wound healing rate of peptide FFFIKLLI treatment on integrin knockdown HuH-7 cells. Peptide concentration, 100 μM. Kruskal-Wallis with Dunn's multiple comparisons test was used for analysis of the data.  Error bars represent s.d..

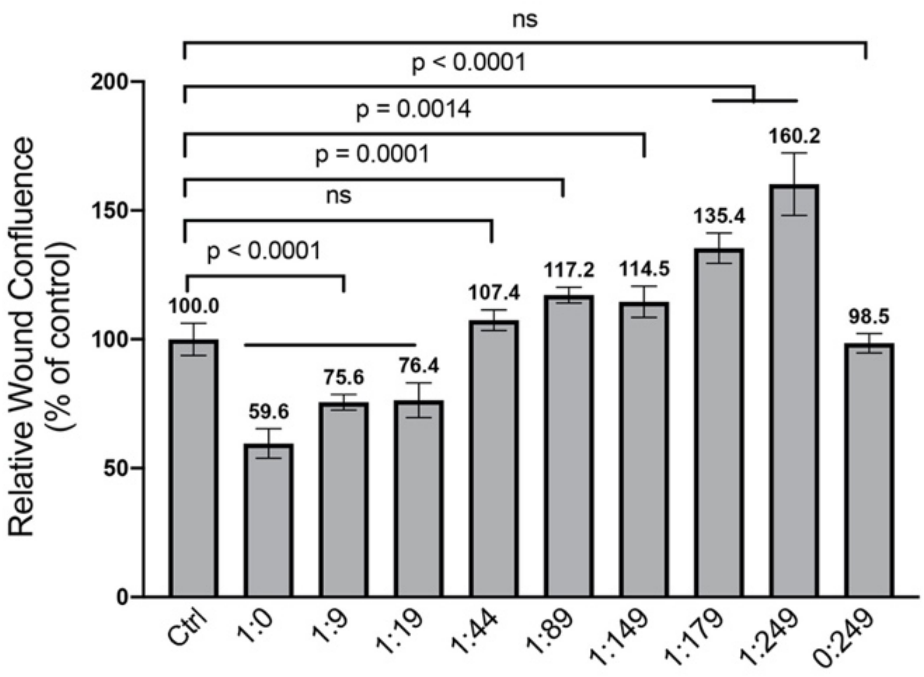


**Fig. S6.**

Wound healing rate of HuH-7 cells upon the treatment of co-assembled FFFIKLLI (fixed at 100 μM) and FFF at various ratios.

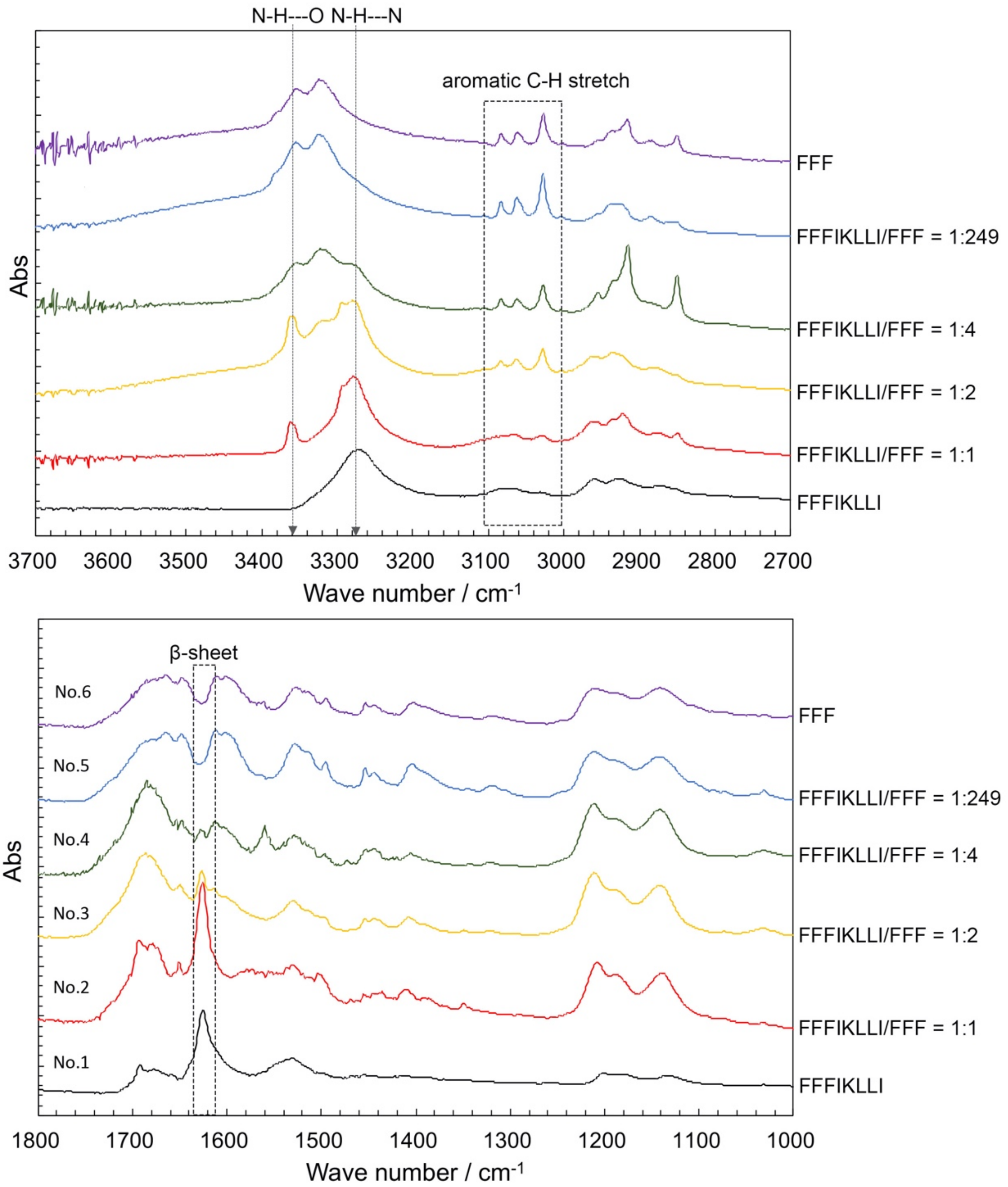


**Fig. S7.**

FTIR spectra of peptide assemblies of FFFIKLLI (100 μM), mixture of FFFIKLLI (100 μM) and FFF in various ratios, and FFF (100 μM) formed in aqueous solution at room temperature.

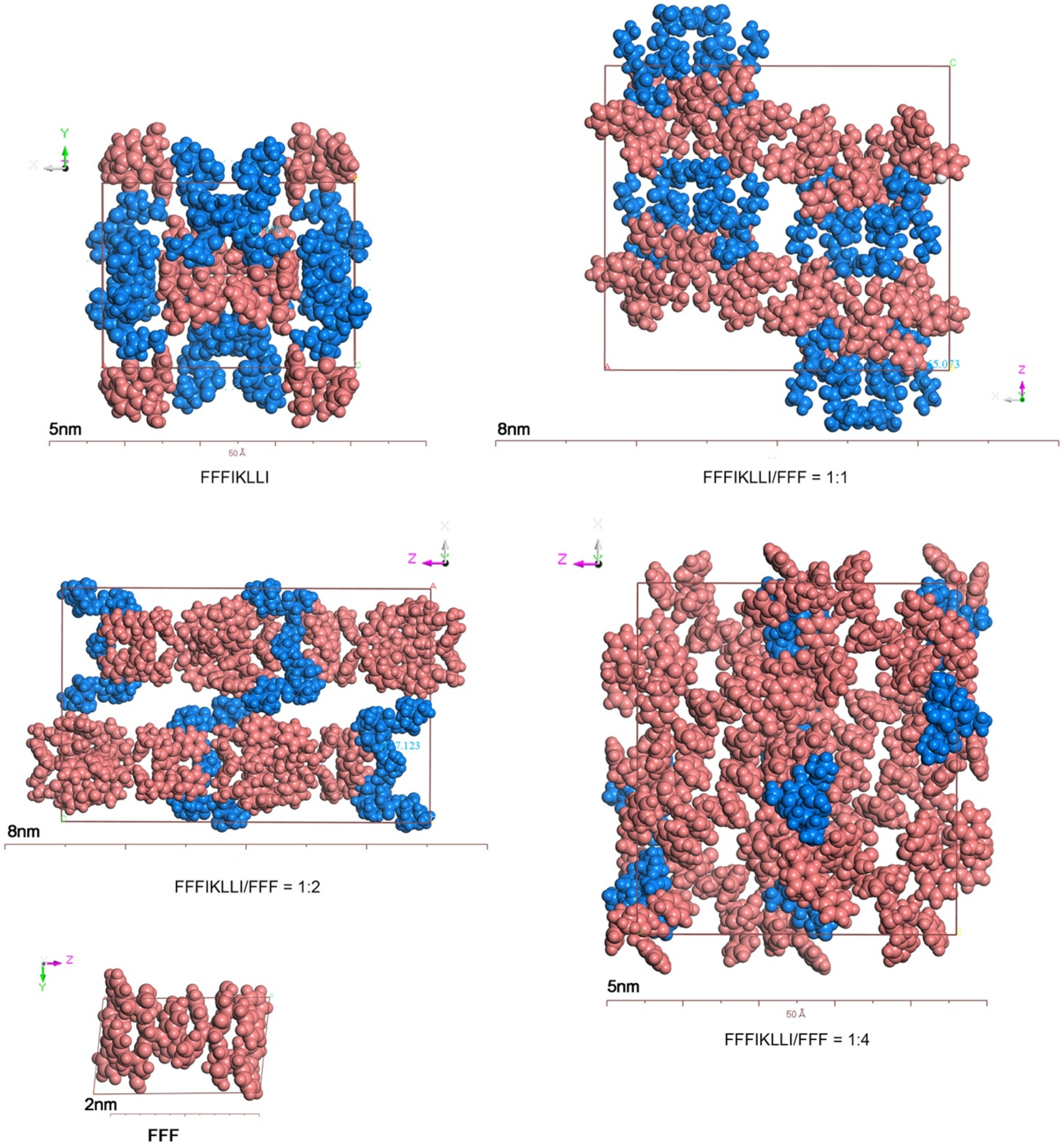


**Fig. S8.**

Predictions of crystal structure unit cell with proper hydrogen bonding types matching to the FTIR analysis presented in space-filling model. Pink represents FFF motif, and blue represents IKLLI motif.

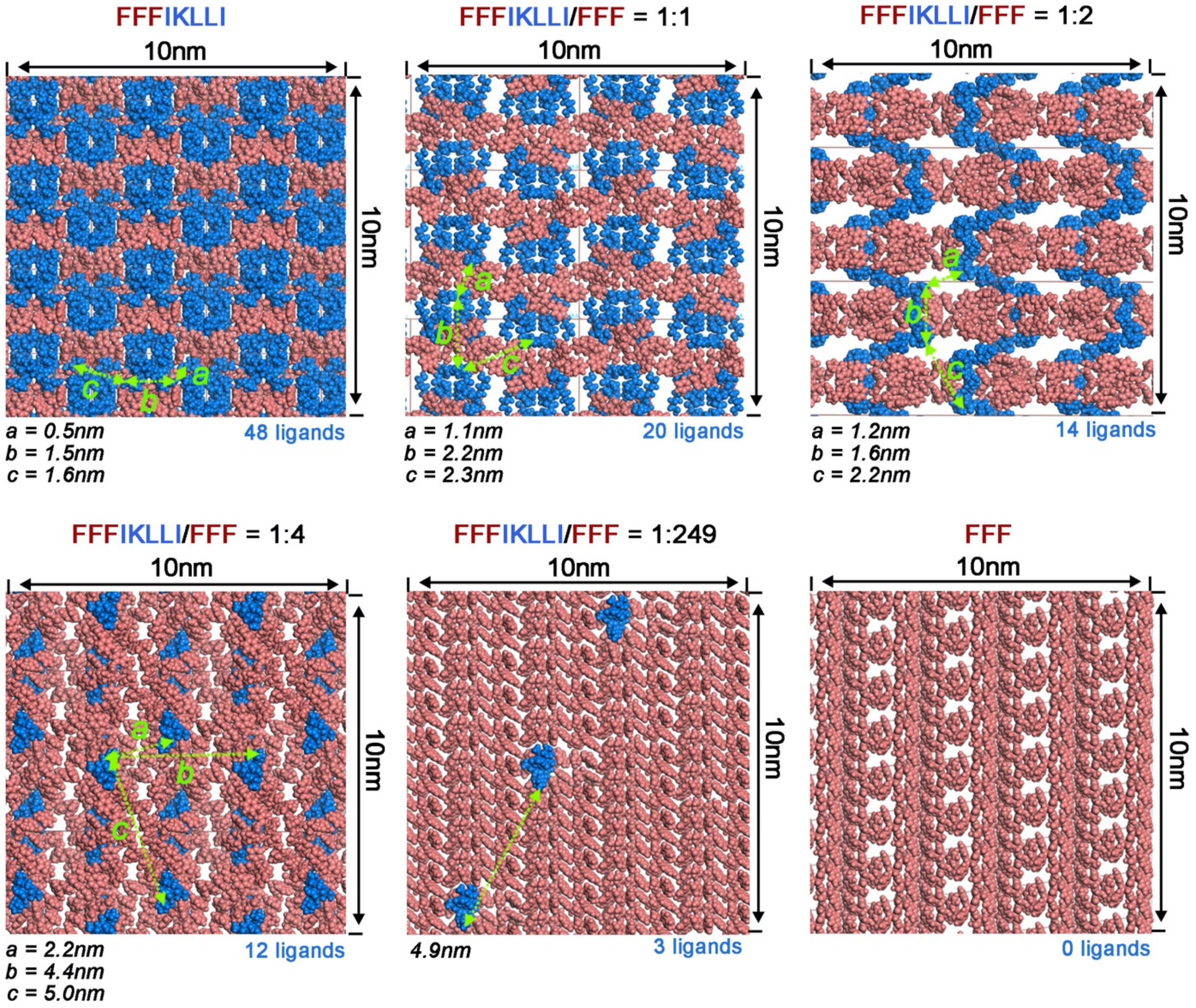


**Fig. S9.**
Space-filling model of surface structure formed by extending the unit cells of polymorph predictions. FFF motif was presented in pink, while IKLLI motif was presented in blue.

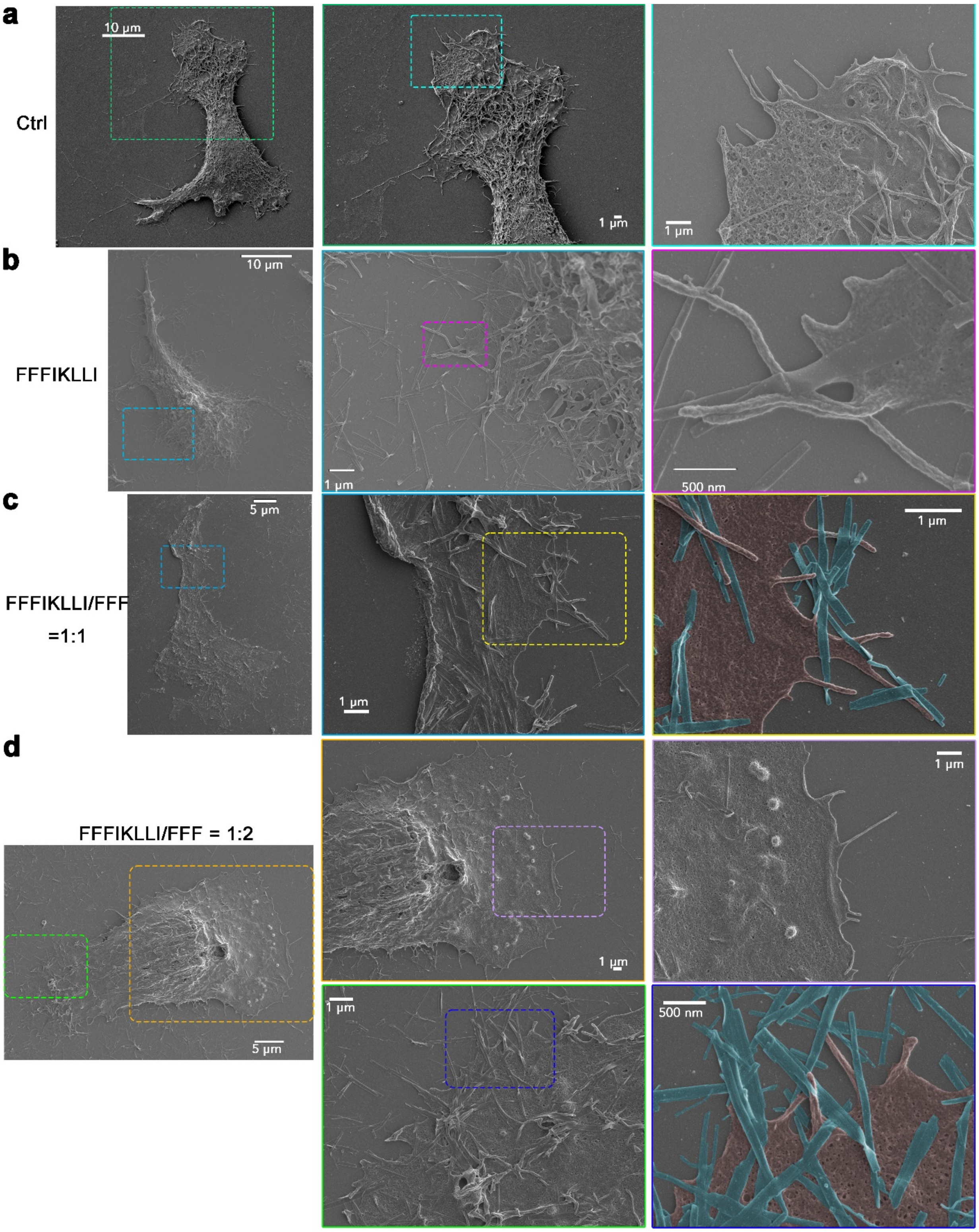

a
Ctrl
10 μm
1 μm
1 μm
b
FFFIKLLI
10 μm
1 μm
500 nm
c
FFFIKLLI/FFF
=1:1
5 μm
1 μm
1 μm
d
FFFIKLLI/FFF = 1:2
5 μm
1 μm
1 μm
1 μm
500 nm

e

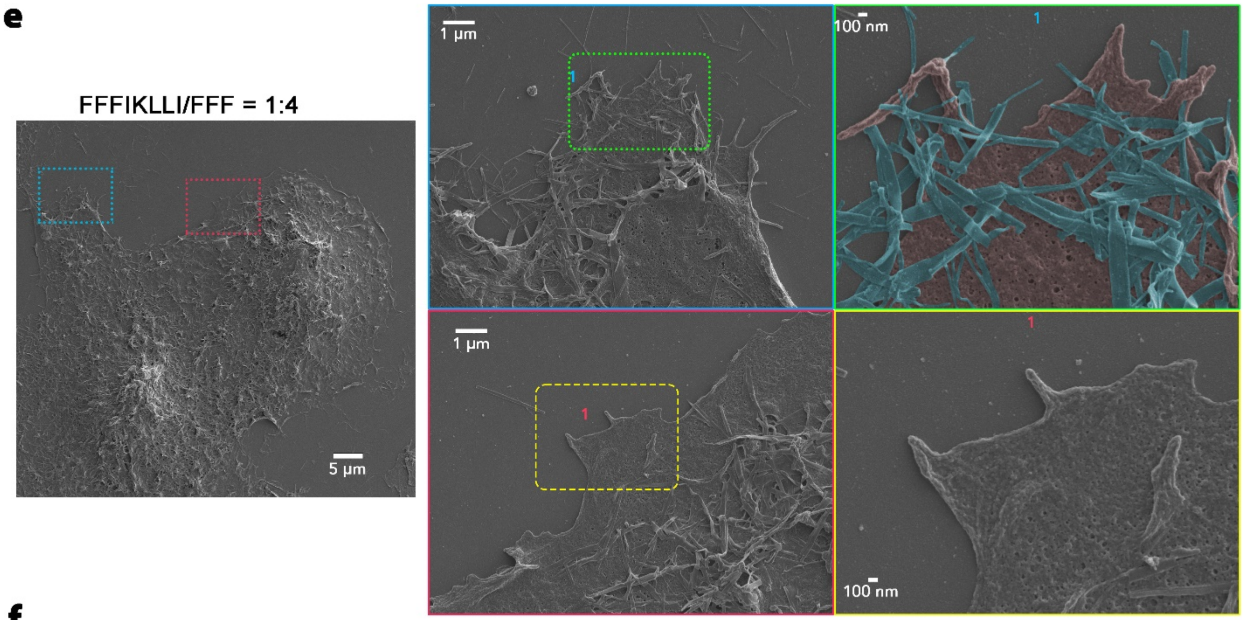


f

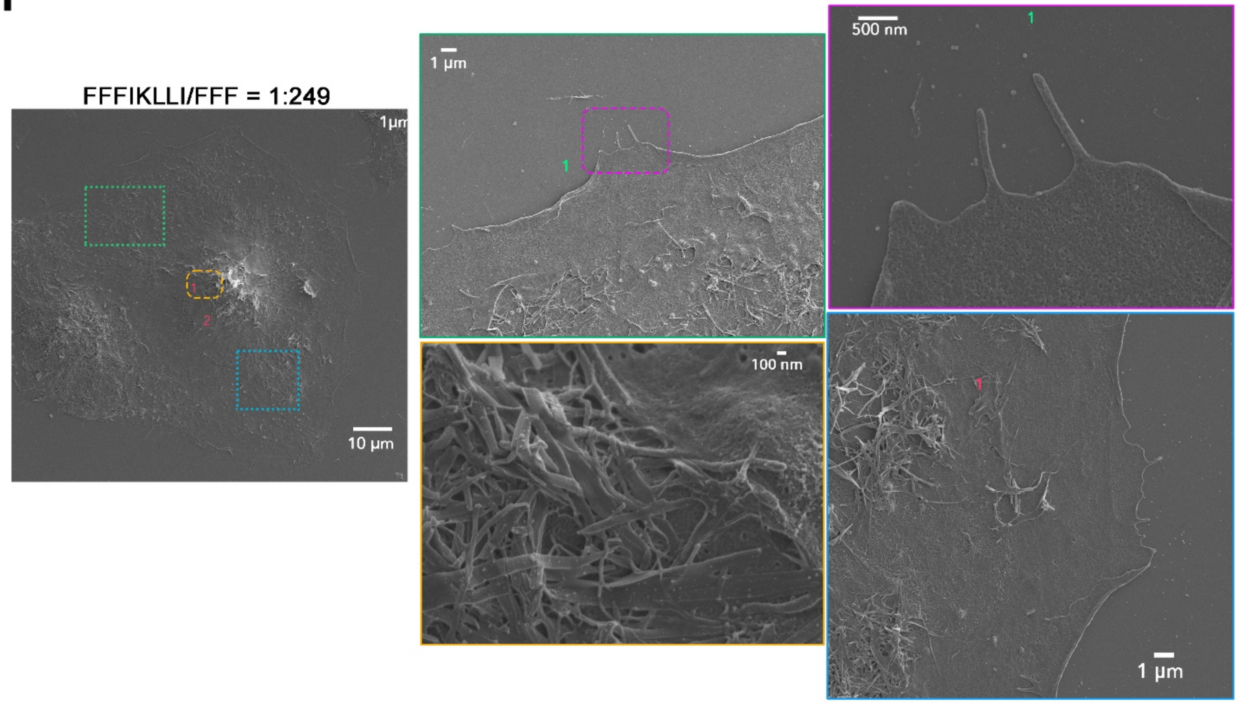


**Fig. S10.**

SEM images of HuH-7 cell with or without the treatment of nanofilaments formed by FFFIKLLI (100 µM) and FFF at various ratios for 12 hr. Cell bodies were highlighted in pink and the nanofilaments were highlighted in blue for false color.

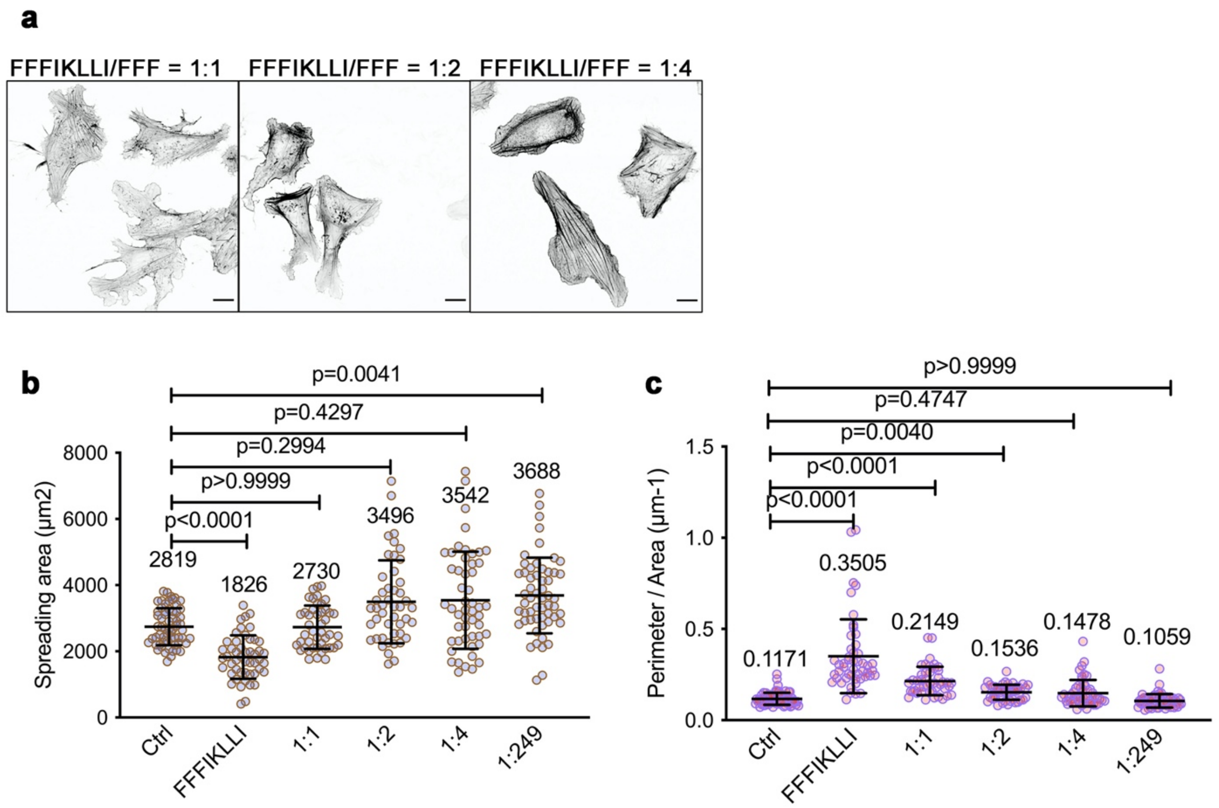


**Fig. S11.**

**(a).** The phalloidin staining of HuH-7 cells incubated with nanofilaments assembled by FFFIKLLI and FFF at various ratios for 12 hr. Scale bar, 20 μm. **(b, c).** The spreading area and the perimeter area ratio of HuH-7 cells for each condition. Kruskal-Wallis with Dunn's multiple comparisons test was used for analysis of the data. Error bars represent standard deviation. n= 61, 49, 45, 46, 51, 56 cells, respectively.

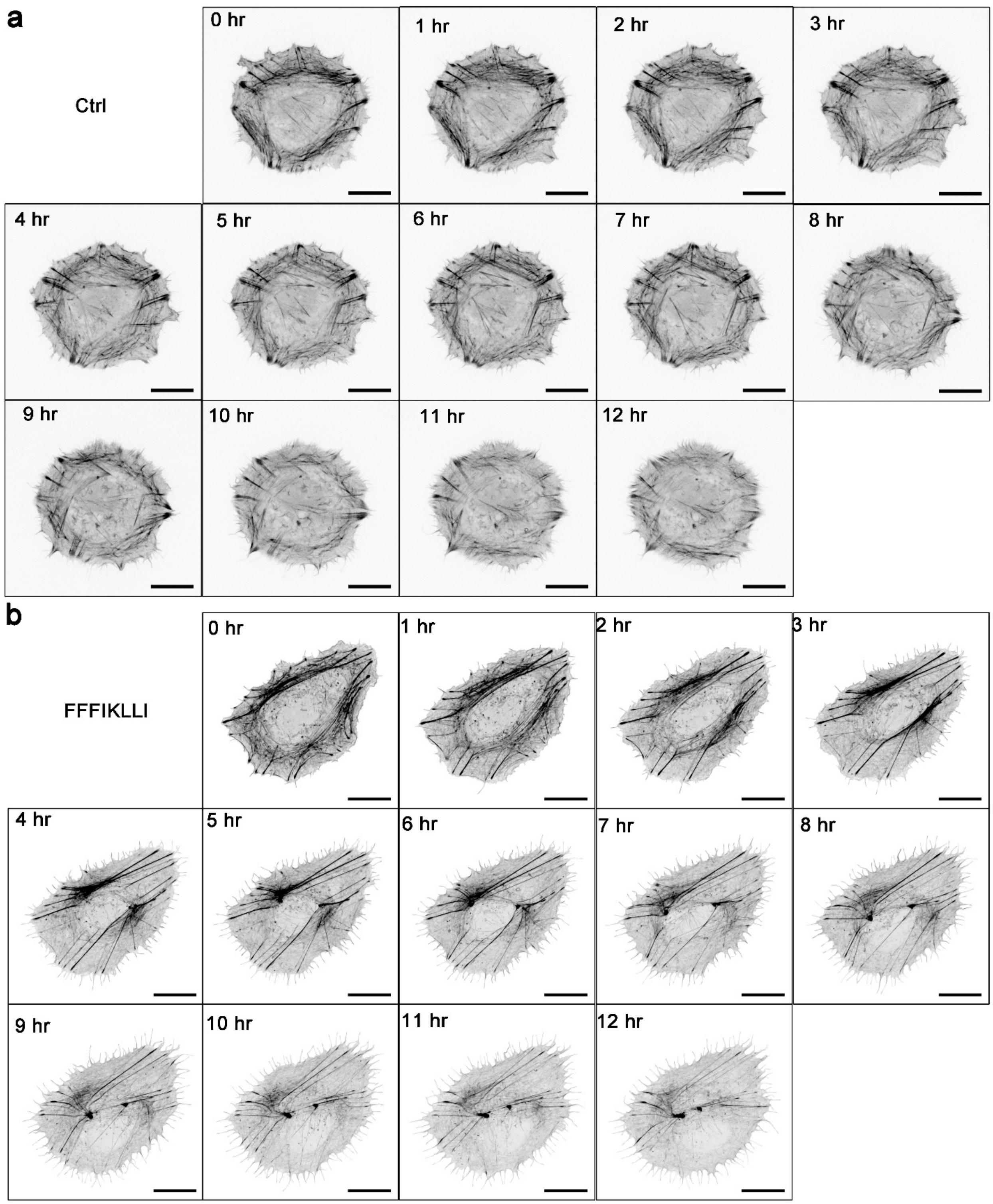
a
Ctrl
0 hr
1 hr
2 hr
3 hr
4 hr
5 hr
6 hr
7 hr
8 hr
9 hr
10 hr
11 hr
12 hr
b
FFFIKLLI
0 hr
1 hr
2 hr
3 hr
4 hr
5 hr
6 hr
7 hr
8 hr
9 hr
10 hr
11 hr
12 hr

c

FFFIKLLI/FFF = 1:1

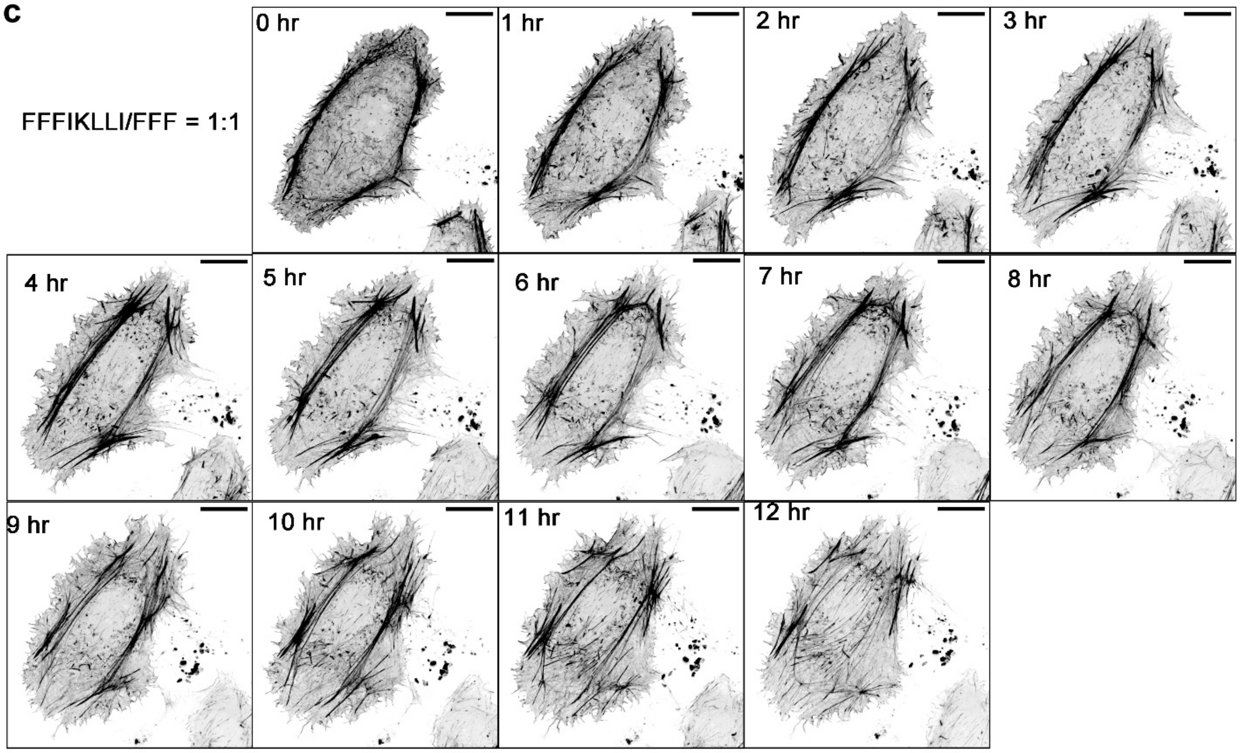


d

FFFIKLLI/FFF = 1:2

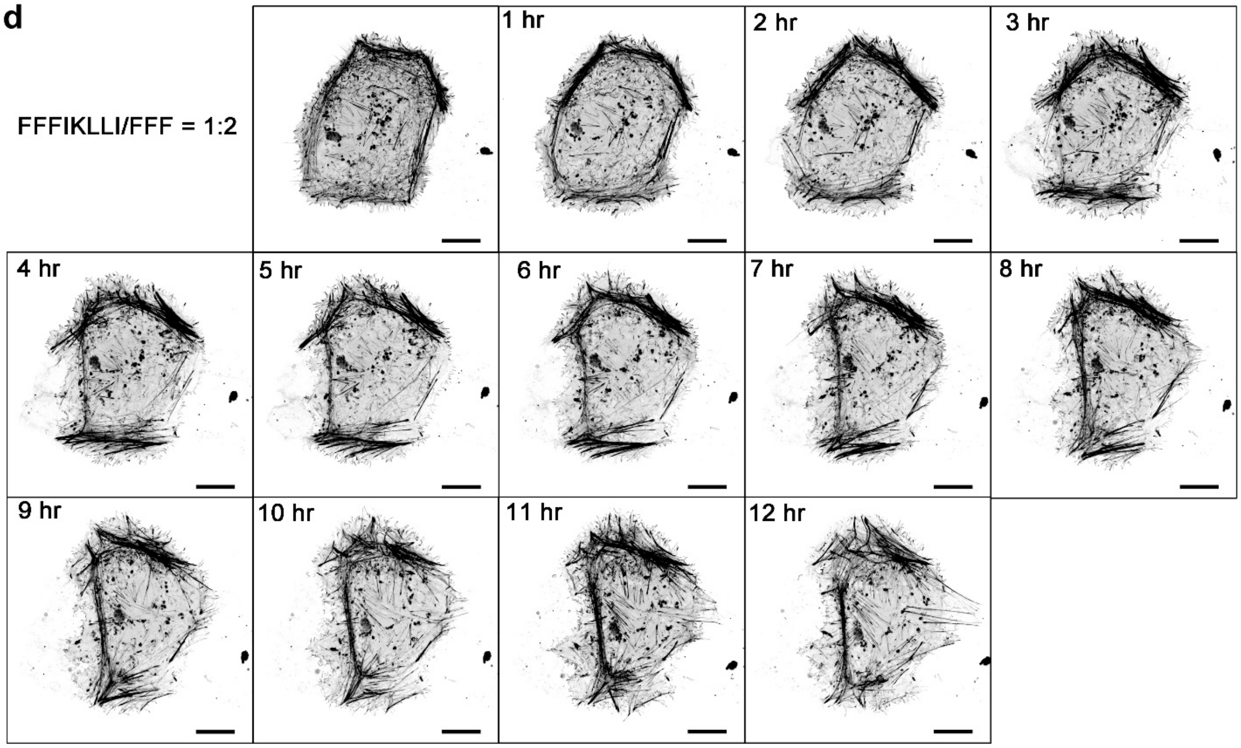

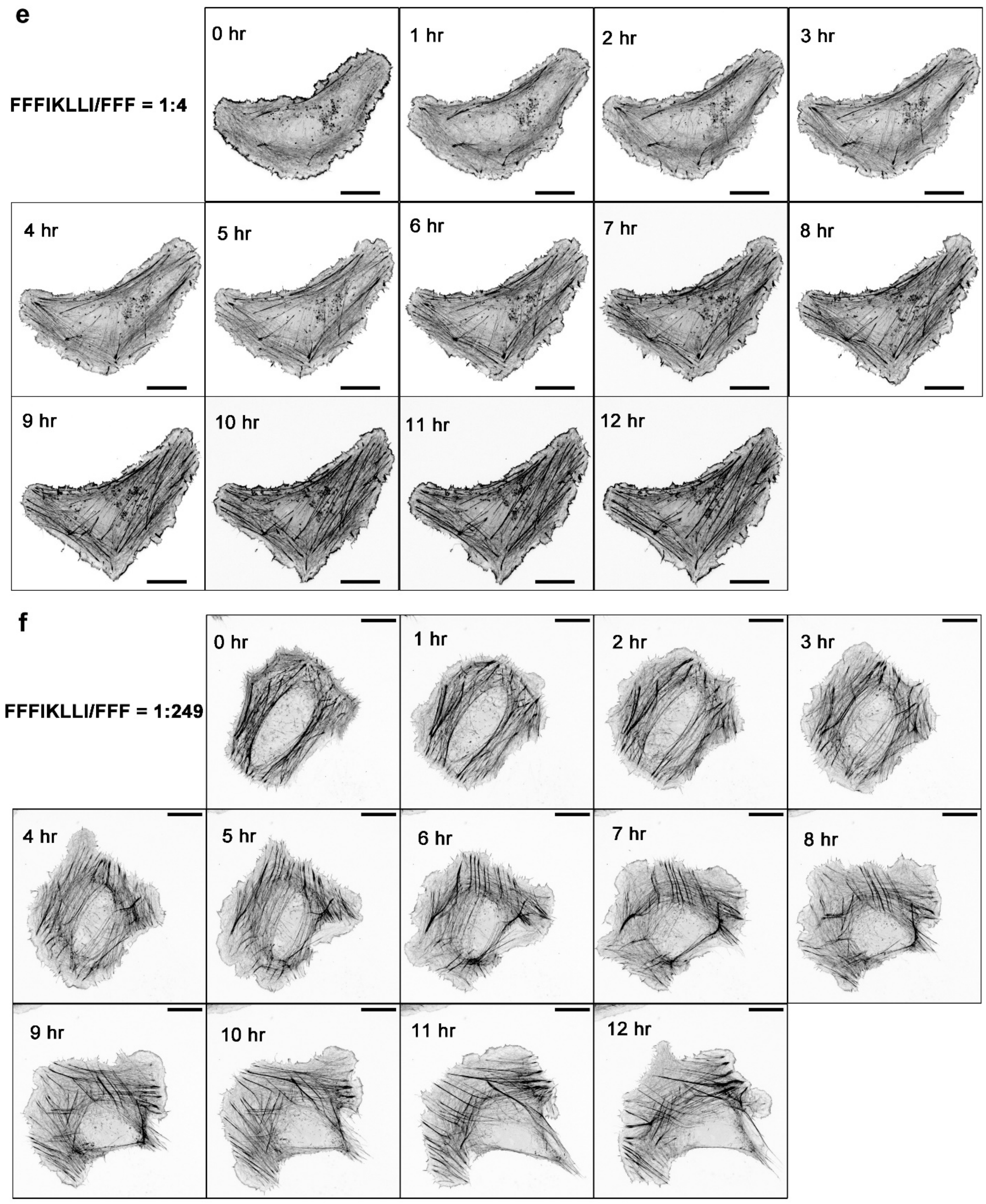


**Fig. S12**.

Time-lapse images of mRuby-Lifeact-7 transfected HuH-7 cells with the treatment of nanofilaments for 12 hr. Scale bar, 20 μm.

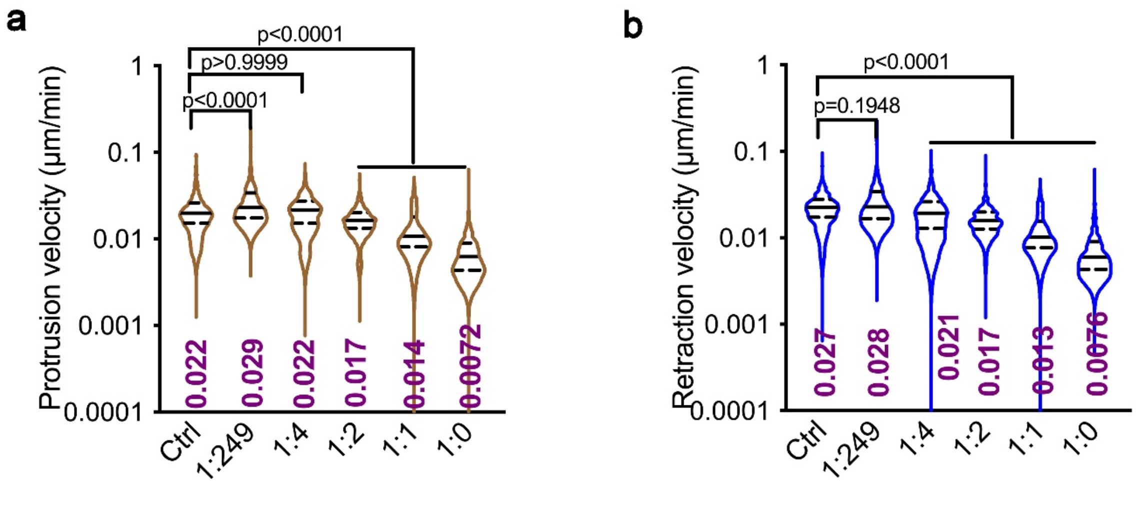


**Fig. S13.**

Violin plot of all protrusion or retraction velocity values collected from each time point for every cell. Kruskal-Wallis with Dunn's multiple comparisons test was used for analysis of the data. Median and quartiles were presented in the plot. n = 10 cells for each group.

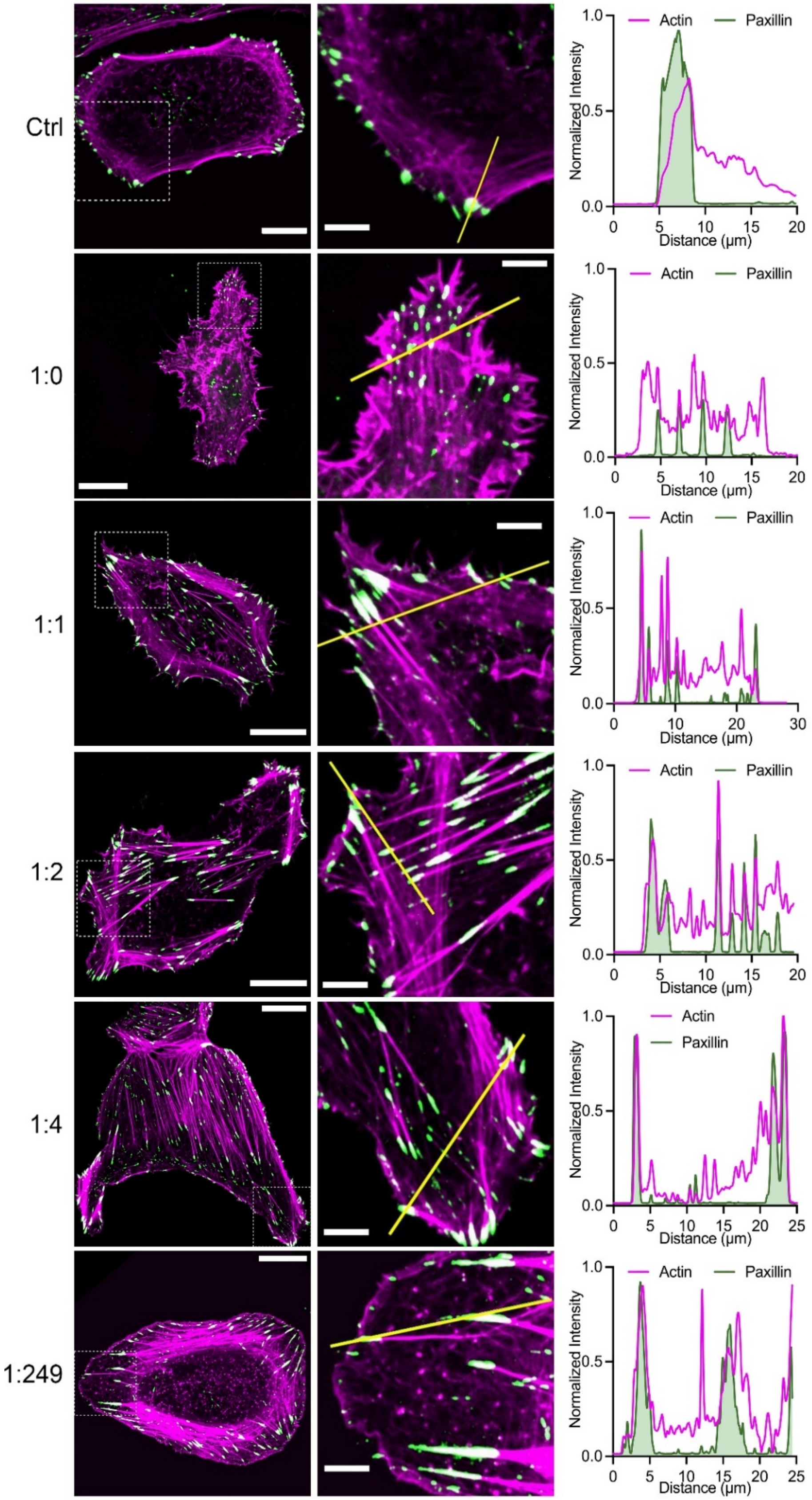


**Fig. S14.**

Immunofluorescence of paxillin co-stain with phalloidin for each treatment condition. Grenn: anti-paxillin; magenta: phalloidin. Scale bar, 20 μm (the first column) and 5 μm (the second column). The third column, intensity profiles.

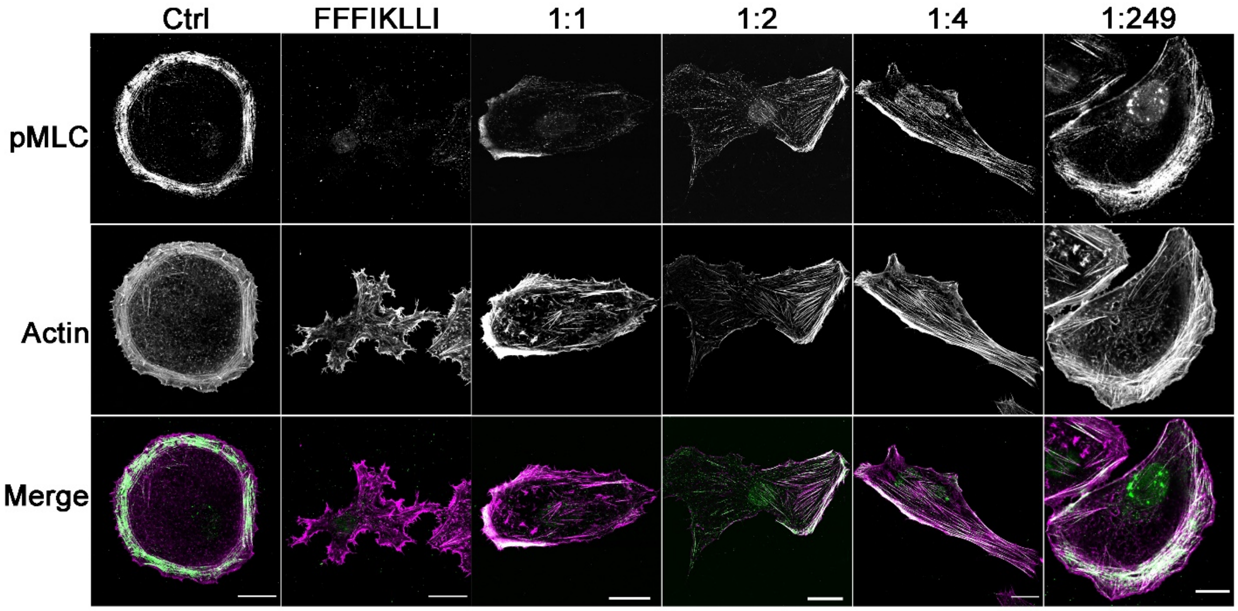


**Fig. S15.**
Immunofluorescence of phospho-Myosin Light Chain 2 (pMLC) co-stain with phalloidin for each treatment condition. The first row, immunofluorescence of pMLC; the second row, phalloidin staining; the third row, merge images, green: anti-pMLC, magenta, phalloidin. Scale bar: 20 μm.

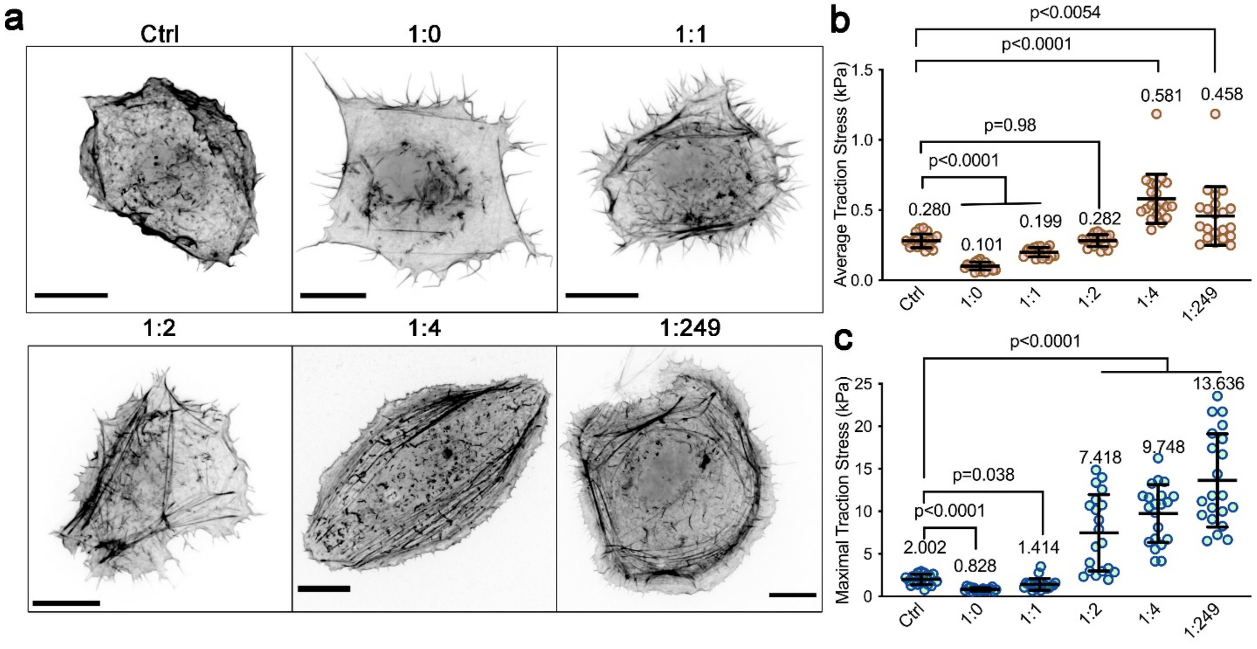


**Fig. S16. Peptide assemblies with various ligand densities control the cell traction force.**

**(a)** Morphology of HuH-7 cells cultured on PDMS subtracts. Cell was transfected with mRuby-Lifeact-7 and treated with peptide assemblies for 12 hr. Scale bar, 20 μm. **(b-c)** Average and Maximal traction stress of each condition. Kruskal-Wallis with Dunn's multiple comparisons test was used for analysis of the data. Error bars represent standard deviation. n = 20 (Ctrl), 18 (1:0), 19 (1:1), 20 (1:2), 21 (1:4), 21 (1:249) cells.

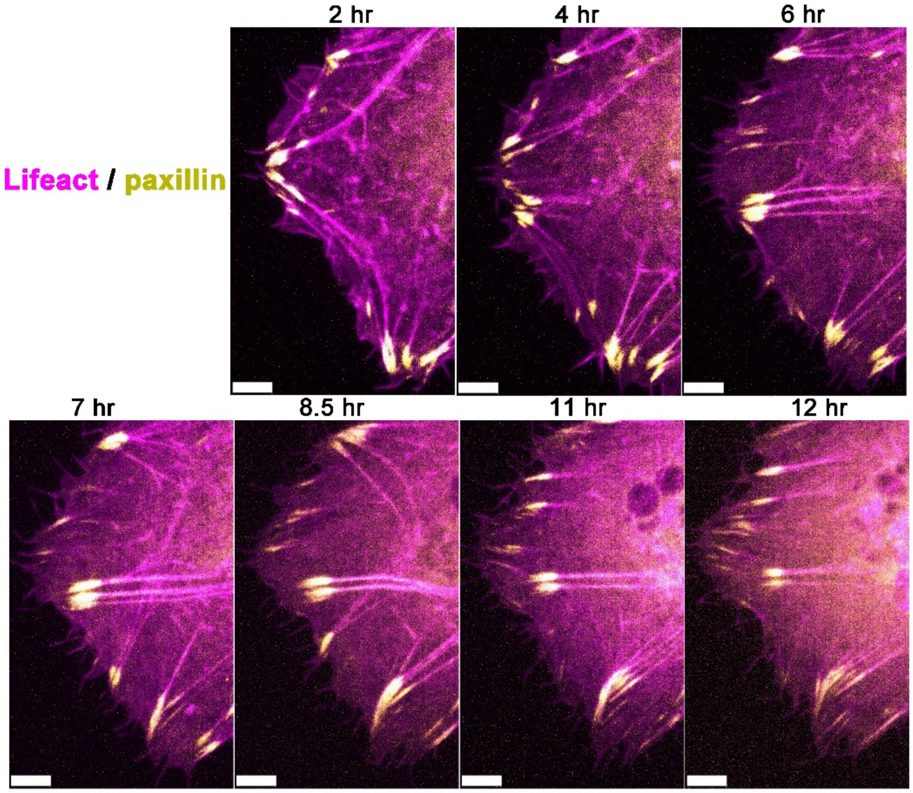


**Fig. S17.**

Time-lapse images of mRuby-Lifeact-7 and pEGFPC1-mEGFP-paxillin co-transfected HuH-7 cells without the treatment of nanofilaments for 12 hr. Scale bar, 2 μm.

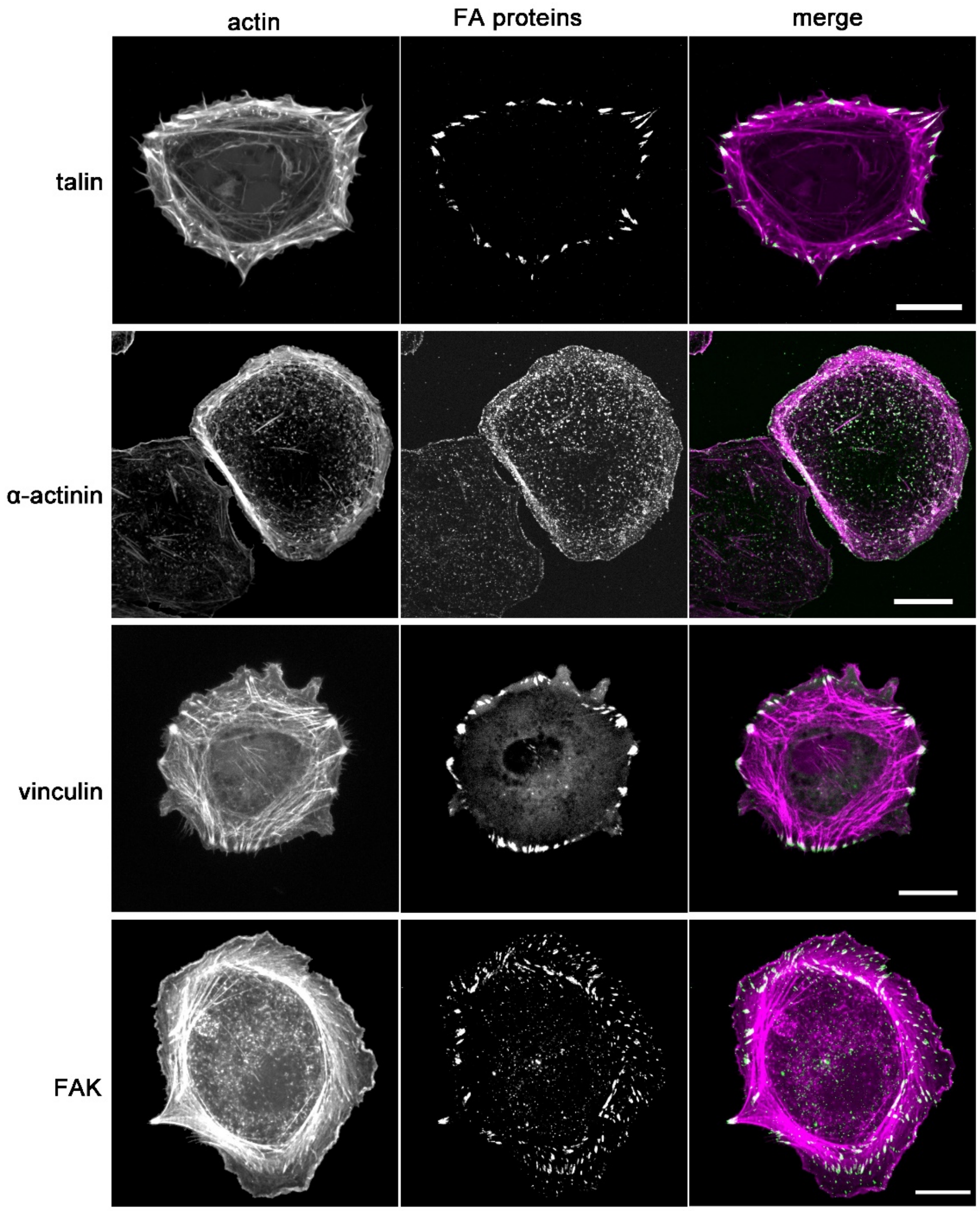


**Fig. S18.**

Immunofluorescence of focal adhesion (FA) proteins co-stain with phalloidin. HuH-7 cells were incubated without FFFIKLLI. The first column, phalloidin staining; the second column, immunofluorescence of FA proteins; the third row, merge images, green: FA proteins, magenta, phalloidin. Scale bar: 20 μm.

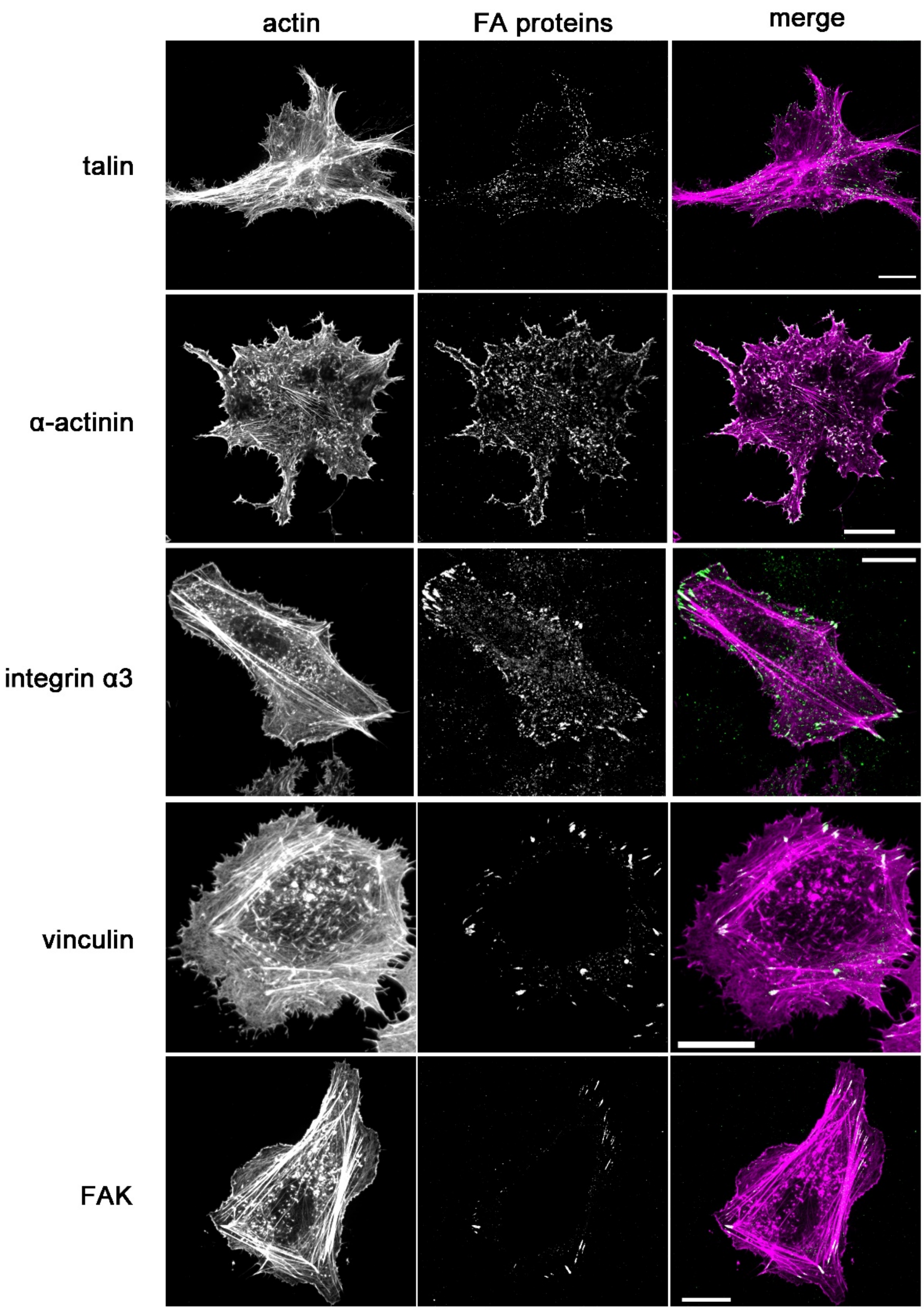


**Fig. S19.**

Immunofluorescence of focal adhesion (FA) proteins co-stain with phalloidin. HuH-7 cells were incubated with FFFIKLLI for 12 hr before fixed. The first column, phalloidin staining; the second column, immunofluorescence of FA proteins; the third row, merge images, green: FA proteins, magenta, phalloidin. Scale bar: 20 μm.

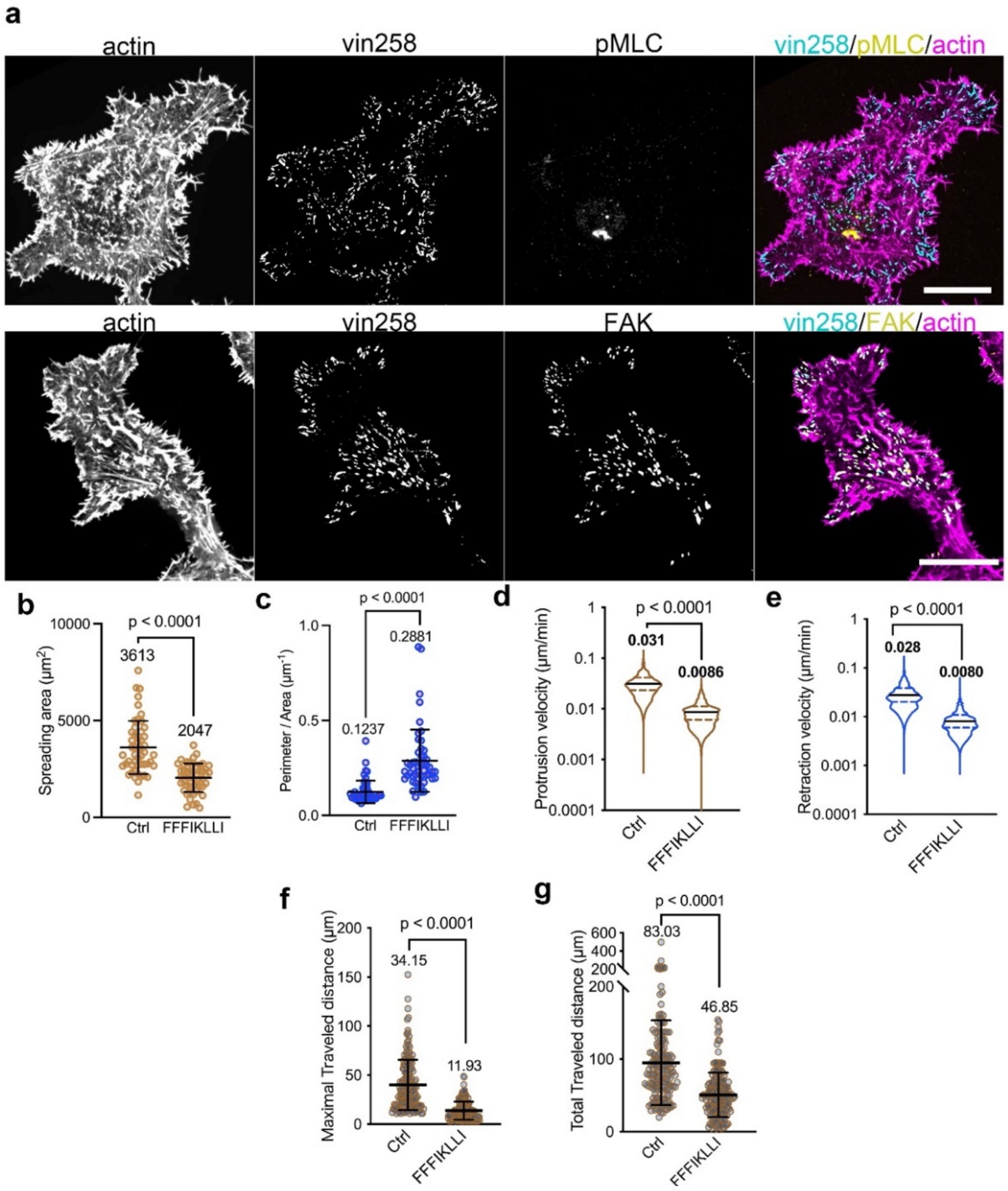


**Fig. S20. Vin258 maintained the FAs on the periphery edge.**

**(a).** Immunofluorescence of pMLC or FAK co-stain with phalloidin. HuH-7 cells were transfected with Pefgpc1/Gg Vcl 1-258 and treated with FFFIKLLI for 12 hr. Scale bar, 20 μm. **(b-c).** Spreading area and P/A ratio of vin258-transfected HuH-7 cells with or without treating with FFFIKLLI for 12 hr. Mann-Whitney test was used for analysis of the data. Error bars represent standard deviation. n = 51 cells. **(d-e).** Violin plot of all protrusion or retraction velocity values collected from each time point for transfected HuH-7 cells pretreated with or without FFFIKLLI for 12 hr. Mann-Whitney test was used for analysis of the data. Median and quartiles were presented in the plot. n = 10 cells for each group. **(f-g).** The maximal traveled distance and total travelled distance of randomly selected migrating transfected HuH-7 cells. Mann-Whitney test was used for analysis of the data. Error bars represent standard deviation. n= 169, 176 cells, respectively.

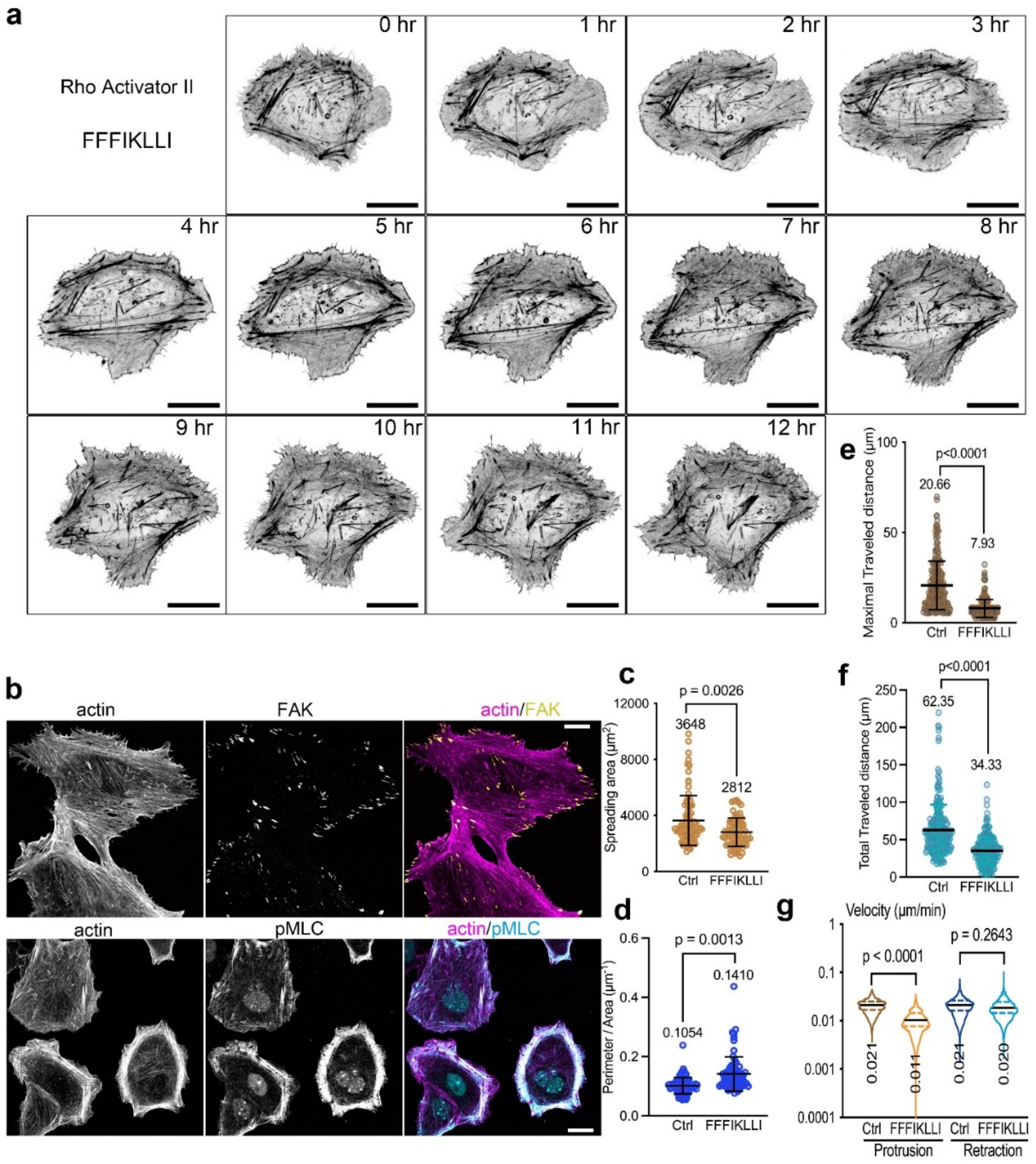


**Fig. S21. Rho activations reboot trailing tail retraction.**

**(a).** Time-lapse imaging of Rho-preactivated HuH-7 cells (mRuby-Lifeact-7 transfected) upon the treatment of FFFIKLLI. Scale bar, 20 μm. **(b).** Immunofluorescence of FAK and pMLC co-stain with phalloidin. Cells were preincubated with Rho Activator II and treated with FFFIKLLI for 12 hr. Scale bar, 20 μm. **(c, d).** Spreading area and P/A ratio of Rho-preactivated cells with or without treating with FFFIKLLI for 12 hr. Mann-Whitney test was used for analysis of the data. Error bars represent standard deviation. n = 85, 76 cells, respectively. **(e, f).** The maximal traveled distance and total travelled distance of randomly selected migrating Rho-preactivated HuH-7 cells. Mann-Whitney test was used for analysis of the data. Error bars represent standard deviation. n= 219, 221 cells, respectively. **(g).** Violin plot of all protrusion or retraction velocity values collected from each time point for Rho-activated HuH-7 cells and treated with or without FFFIKLLI for 12 hr. Mann-Whitney test was used for analysis of the data. Median and quartiles were presented in the plot. n = 10 cells for each group.

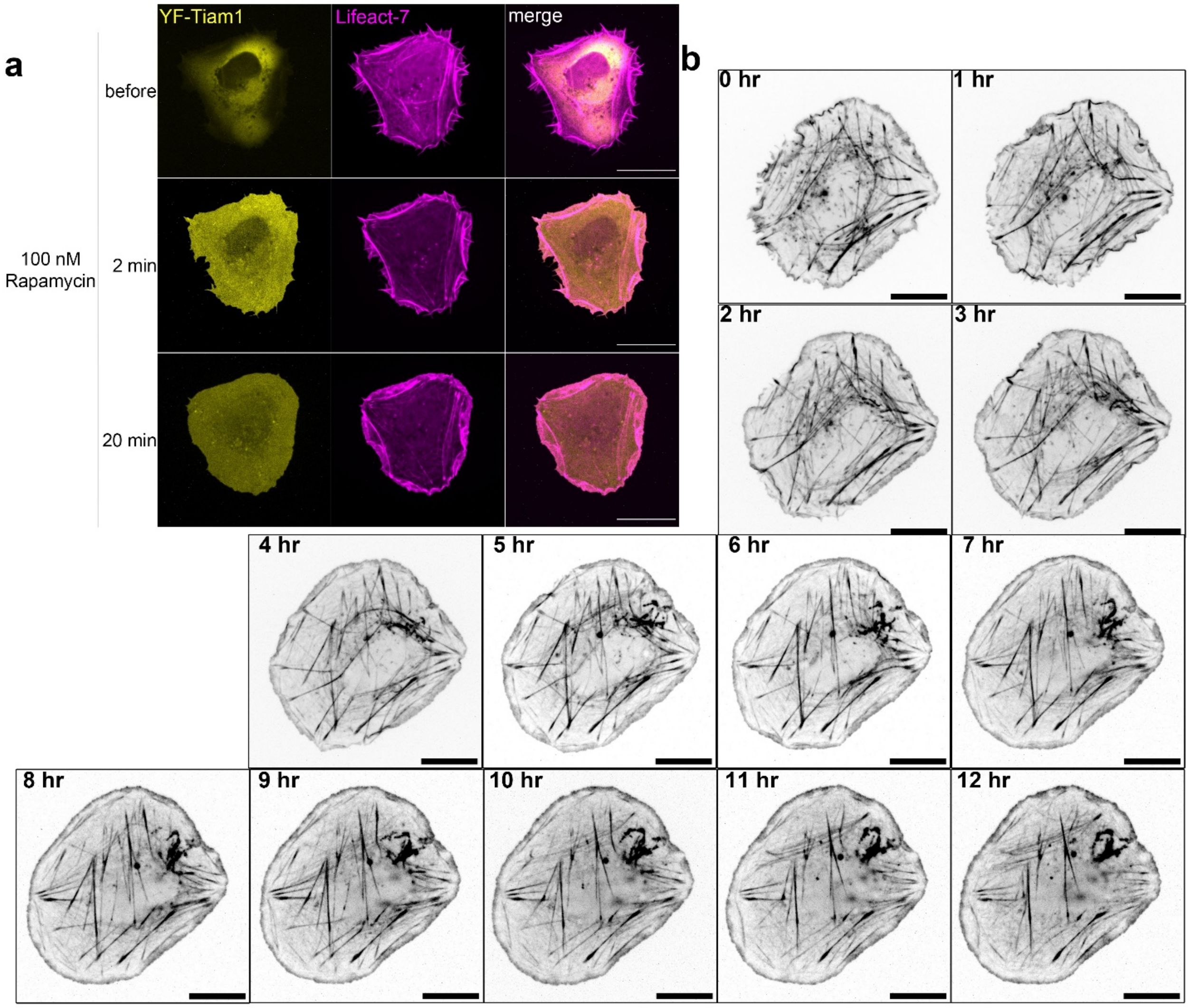


**Fig. S22. Tiam1/Rac1 activation resume the cell protrusion**

**(a).** Lamellipodia stared to form within 2 minutes after Rac1 activation. Scale bar, 20 μm. **(b).** Time-lapse imaging of Rac1-activated HuH-7 cells (co-transfected with mRuby-Lifeact-7). Scale bar, 20 μm.

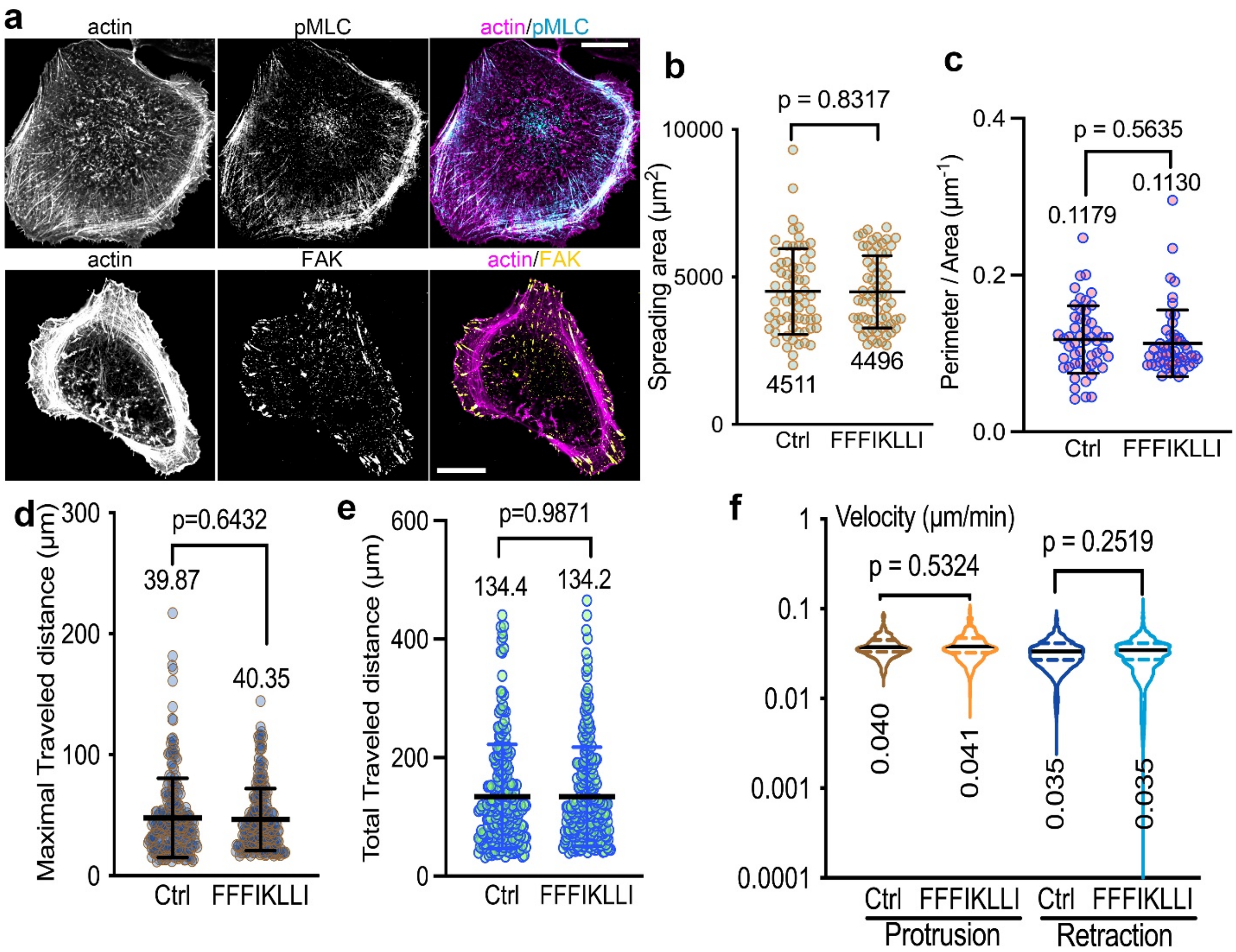


**Fig. S23. Tiam1/Rac1 activation reboots the cell migration.**

**(a).** Immunofluorescence of FAK and pMLC co-stain with phalloidin. Tiam1/Rac1 was preactivated right before starting the 12 hr treatment of FFFIKLLI. Scale bar, 20 μm. **(b, c).** Spreading area and P/A ratio of Rho-preactivated cells with or without treating with FFFIKLLI for 12 hr. Mann-Whitney test was used for analysis of the data. Error bars represent standard deviation. n = 64 cells. **(d, e).** The maximal traveled distance and total travelled distance of randomly selected migrating Tiam1/Rac1-activated HuH-7 cells. Mann-Whitney test was used for analysis of the data. Error bars represent standard deviation. n= 230, 234 cells, respectively. **(f).** Violin plot of all protrusion or retraction velocity values collected from each time point for Tiam1/Rac1-activated HuH-7 cells and treated with or without FFFIKLLI for 12 hr. Mann-Whitney test was used for analysis of the data. Median and quartiles were presented in the plot. n = 10 cells for each group.

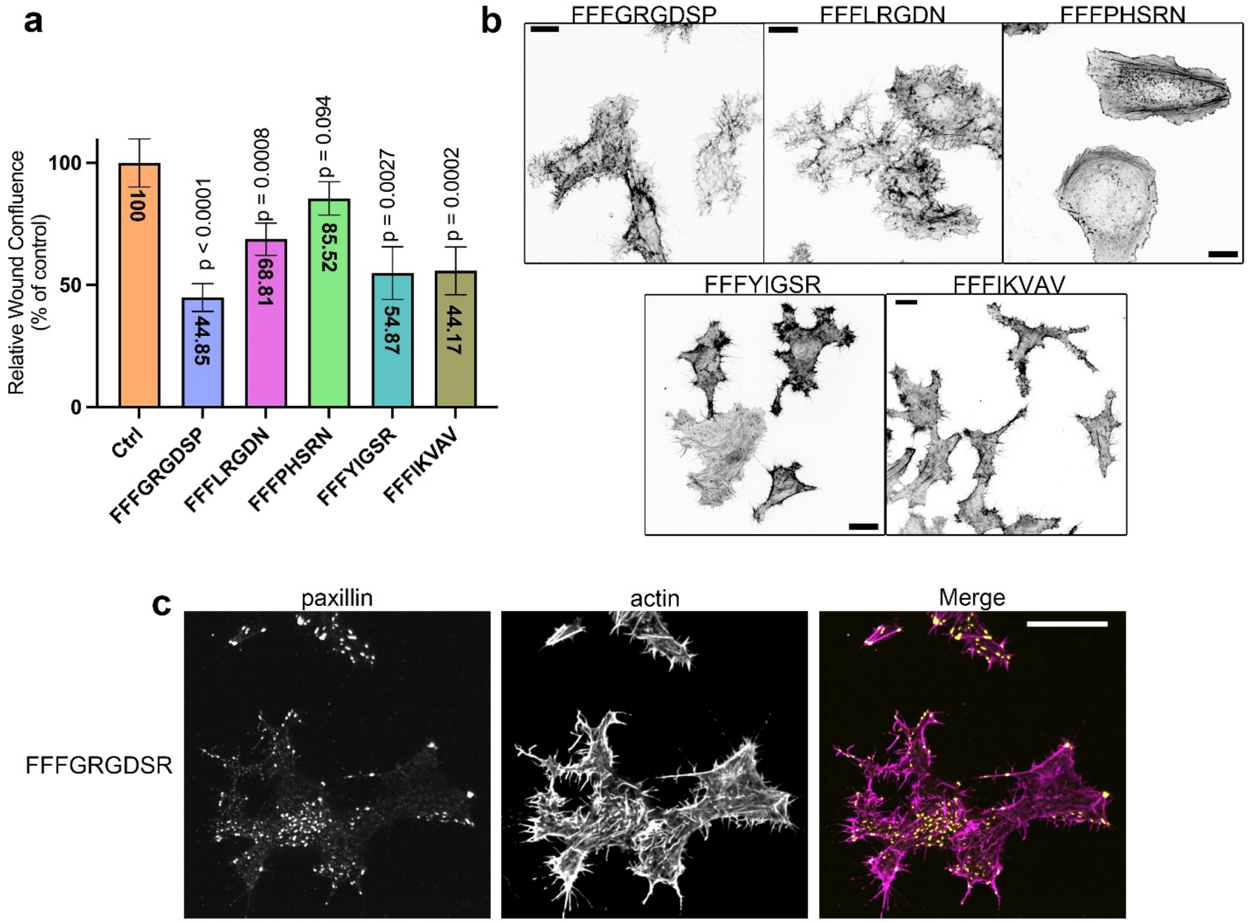


**Fig. S24. Other assembling ligands affect cell migration.**

**(a).** 48hr-Wound healing rate of HuH-7 cells upon the treatment of 200 μM assembling ligands. **(b).** The phalloidin staining of HuH-7 cells incubated with 200 μM assembling ligands for 12 hr. Scale bar, 20 μm. (c). Immunofluorescence of paxillin co-stain with phalloidin. HuH-7 cells were incubated with 200 μM assembling ligands for 12 hr. Scale bar, 20 μm.

| | | | |
|---|---|---|---|
| Lot No. | :P210114-YW125651 | | |
| Column | :Diamonsil C18, 4.6*250mm, 5um | | |
| Solvent A | :0.1%Trifluoroacetic in 100% Acetonitrile | | |
| Solvent B | :0.1%Trifluoroacetic in 100% Water | | |
| Gradient | : | A | B |
| | 0.0min | 32% | 68% |
| | 25.0min | 57% | 43% |
| | 25.1min | 100% | 0% |
| | 30.0min | Stop | |
| Flow rate | :1.0ml/min | | |
| Wavelength | :220nm | | |
| Volume | :20ul | | |

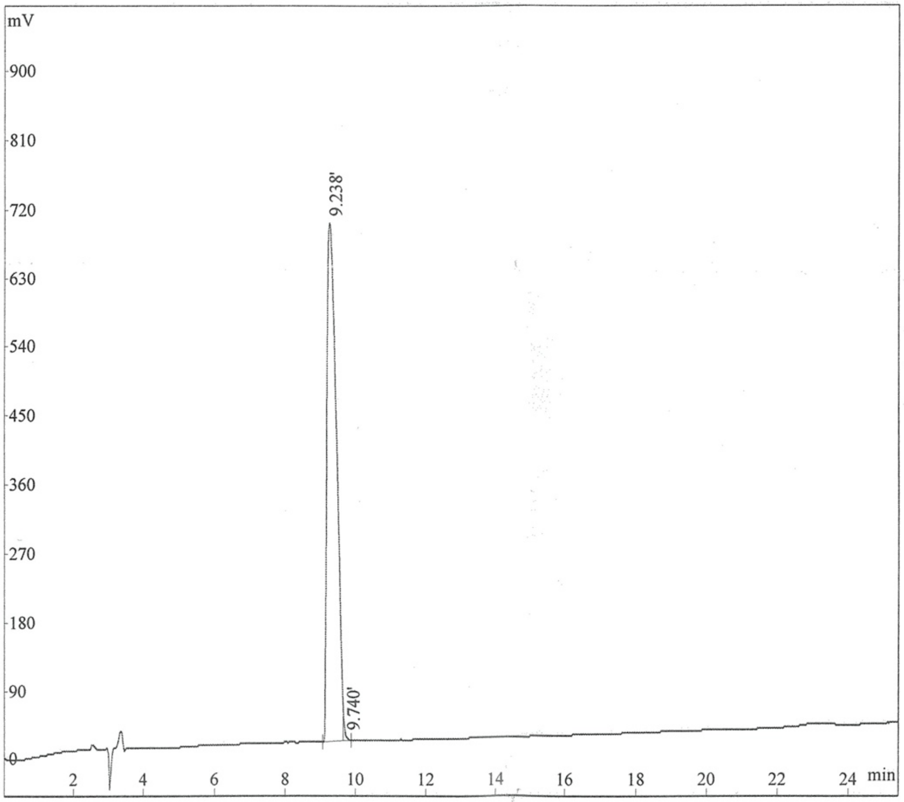


| Rank | Time | Quantity | Area | Height |
|---|---|---|---|---|
| 1 | 9.238 | 99.51 | 11982890 | 679242 |
| 2 | 9.740 | 0.4857 | 58488 | 3542 |
| Total | | 100 | 12041378 | 682784 |

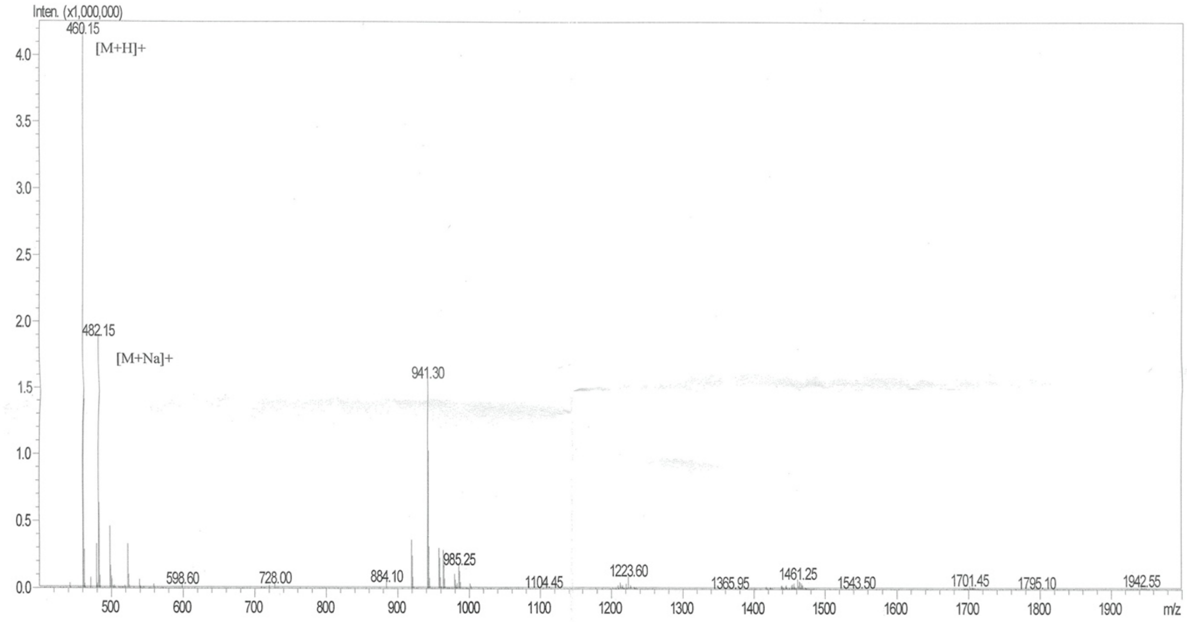


| | | | |
|---|---|---|---|
| Mw | : 459.57 | | |
| Lot No. | : P210114-YW125651 | | |
| Probe | :ESI | Probe bias | :+4.5kv |
| Nebulizer Gas Floe | :1.5L/min | Detector | :1.2kv |
| CDL | :-20.0v | T.Flow | :0.2ml/min |
| CDL Temp | :250℃ | B.conc | :50%H2O/50%ACN |
| Block Temp | :400℃ | | |

**Appendix S1.**

LC-MS spectra of FFF.

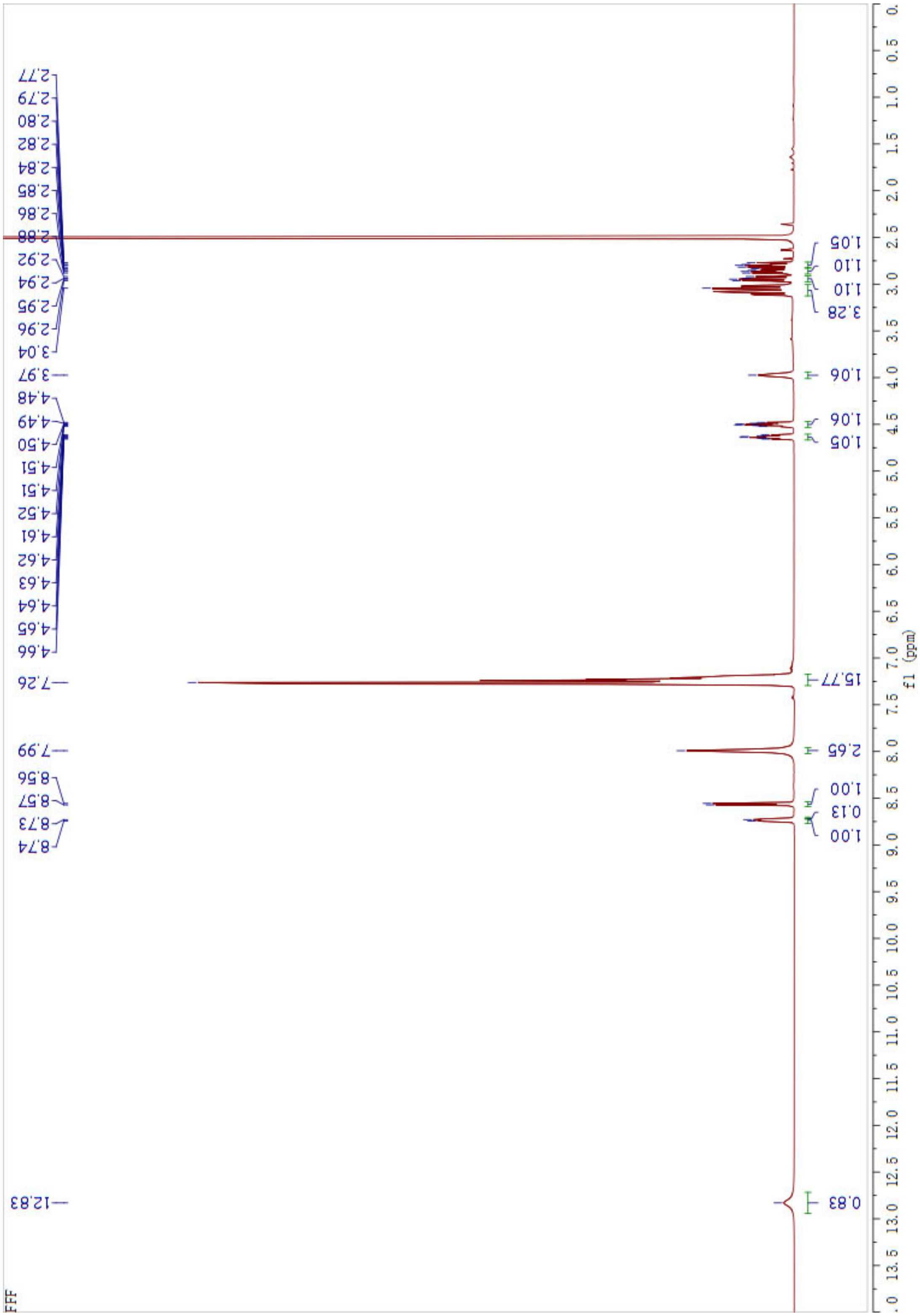


**Appendix S2.**

$^{1}H$ NMR of FFF.

Lot NO :P190510-YS725631
Number :0200049
Column :250*4.6mm,Kromasil-C18-5um
Solvent A:0.1%TFA in 100%water
Solvent B:0.1%TFA in 100%acetonitrile

| Gradient : | A | B |
|---|---|---|
| 0.1min | 68% | 32% |
| 25.0min | 43% | 57% |
| 25.1min | 0% | 100% |
| 30.0min | stop | |

Flow rate:1.0ml/min
Wavelength(nm):220
Volume :10ul

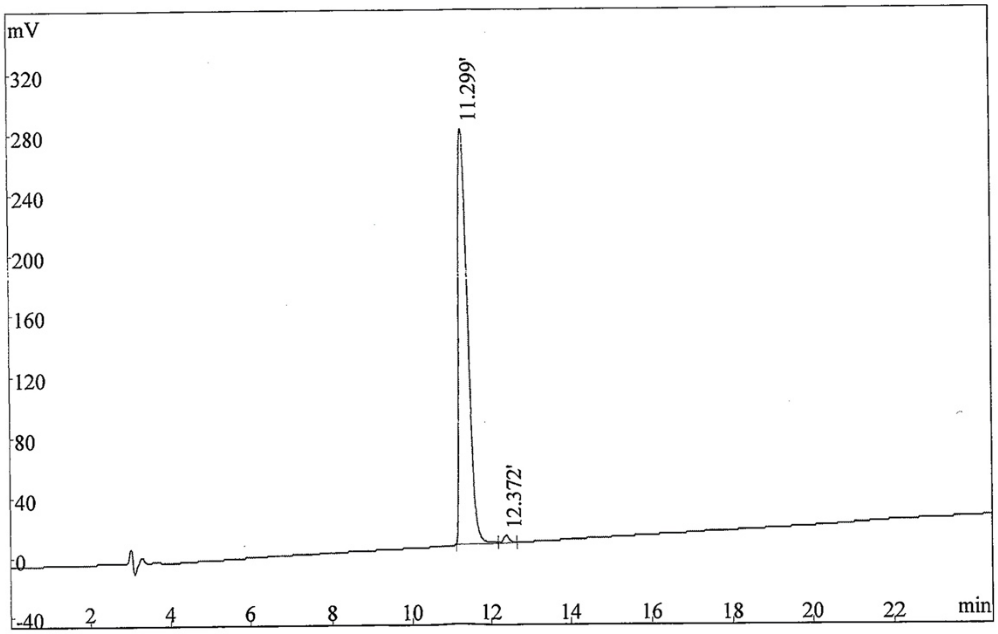


| Rank | Time | Conc. | Area | Height |
|---|---|---|---|---|
| 1 | 11. 299 | 98. 94 | 4131588 | 275311 |
| 2 | 12. 372 | 1. 062 | 44343 | 4677 |
| Total | | 100 | 4175931 | 279988 |

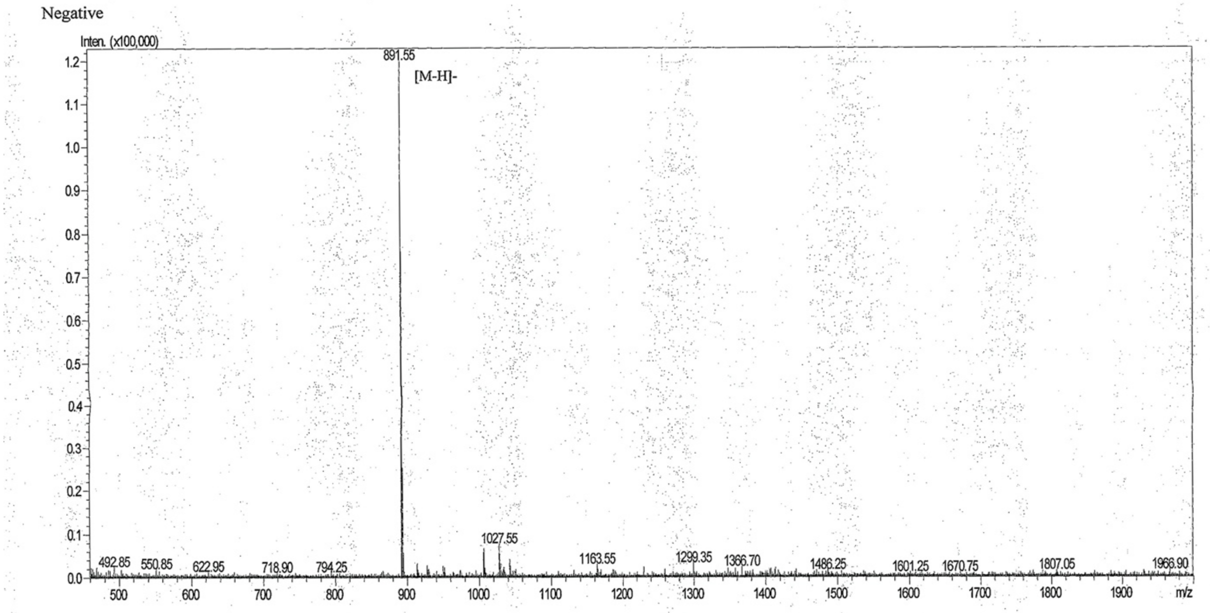


Injection Volume : 1
Sample Name : FI-7
Mw : 893.17
Lot No. : P190510-YS725631

| | | | |
|---|---|---|---|
| Probe | :ESI | Probe bias | :+4.5kv |
| Nebulizer Gas Floe | :1.5L/min | Detector | :1.2kv |
| CDL | :-20.0v | T.Flow | :0.2ml/min |
| CDL Temp | :250℃ | B.conc | :50%H2O/50%ACN |
| Block Temp | :400℃ | | |

**Appendix S3.**

LC-MS spectra of FFIKLLI. The purity of the peptide is 98.94%.

| | |
|---|---|
| Lot No. | :P210114-YW687883 |
| Column | :Diamonsil C18, 4.6*250mm, 5um |
| Solvent A | :0.1%Trifluoroacetic in 100% Acetonitrile |
| Solvent B | :0.1%Trifluoroacetic in 100% Water |

| Gradient | : | A | B |
|---|---|---|---|
| | 0.0min | 40% | 60% |
| | 25.0min | 65% | 35% |
| | 25.1min | 100% | 0% |
| | 30.0min | Stop | |

| | |
|---|---|
| Flow rate | :1.0ml/min |
| Wavelength | :220nm |
| Volume | :20ul |

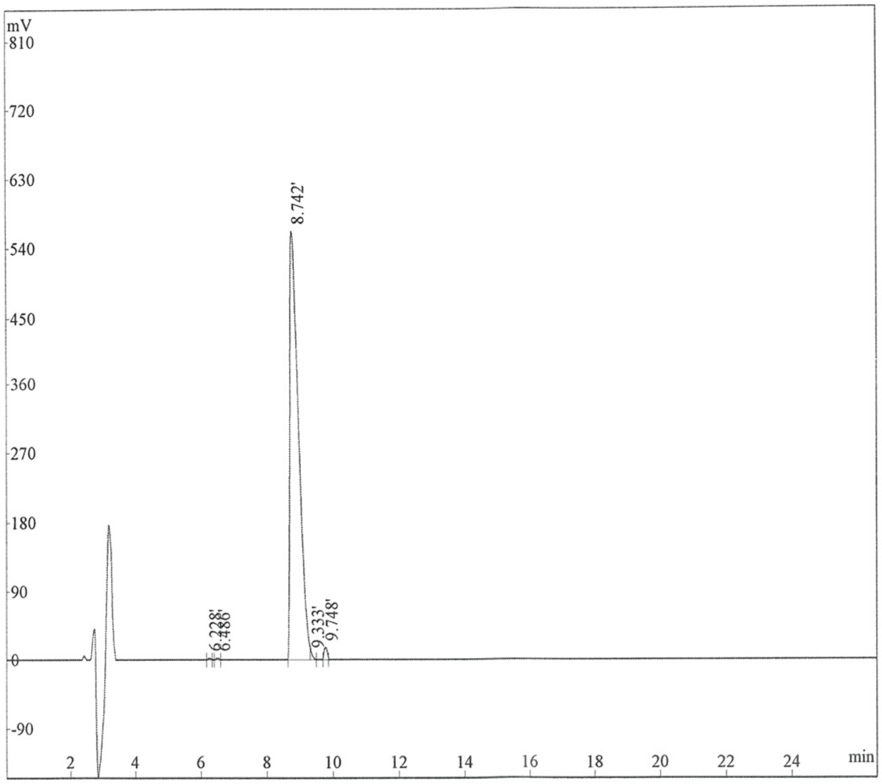


| Rank | Time | Quantity | Area | Height |
|---|---|---|---|---|
| 1 | 6.228 | 0.1375 | 13406 | 2662 |
| 2 | 6.486 | 0.1991 | 19405 | 3339 |
| 3 | 8.742 | 98.32 | 9583033 | 560515 |
| 4 | 9.333 | 0.4022 | 39207 | 6733 |
| 5 | 9.748 | 0.943 | 91913 | 15890 |
| Total | | 100 | 9746964 | 589139 |

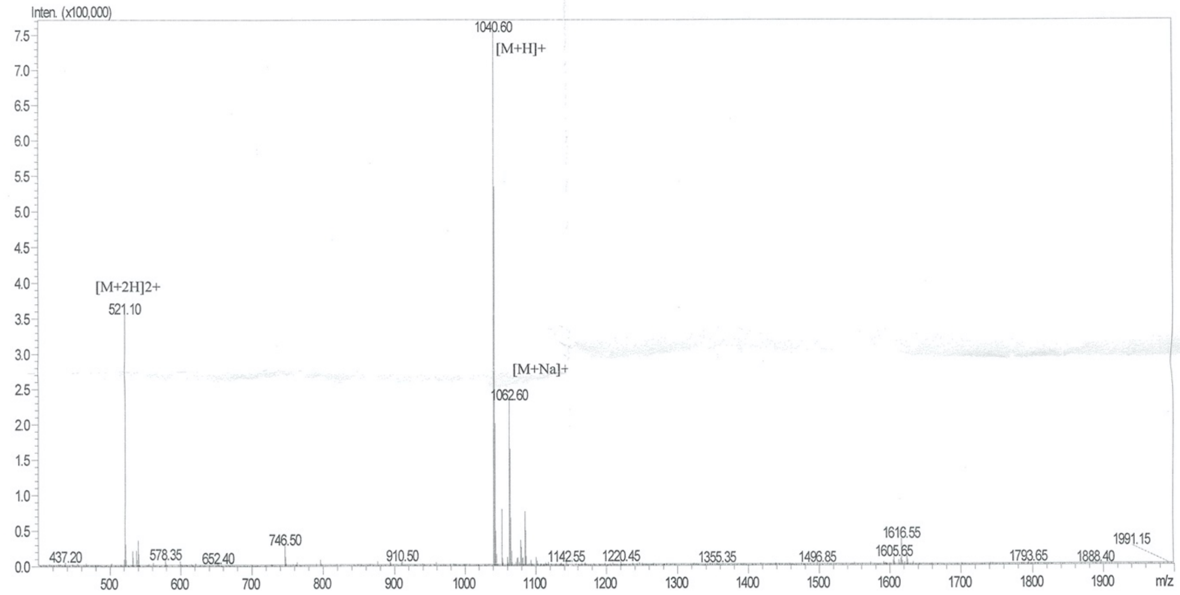


| | | | |
|---|---|---|---|
| Mw | : 1040.44 | | |
| Lot No. | : P210114-YW687883 | | |
| Probe | :ESI | Probe bias | :+4.5kv |
| Nebulizer Gas Floe | :1.5L/min | Detector | :1.2kv |
| CDL | :-20.0v | T.Flow | :0.2ml/min |
| CDL Temp | :250℃ | B.conc | :50%H2O/50%ACN |
| Block Temp | :400℃ | | |

**Appendix S4.**

LC-MS spectra of FFFIKLLI.

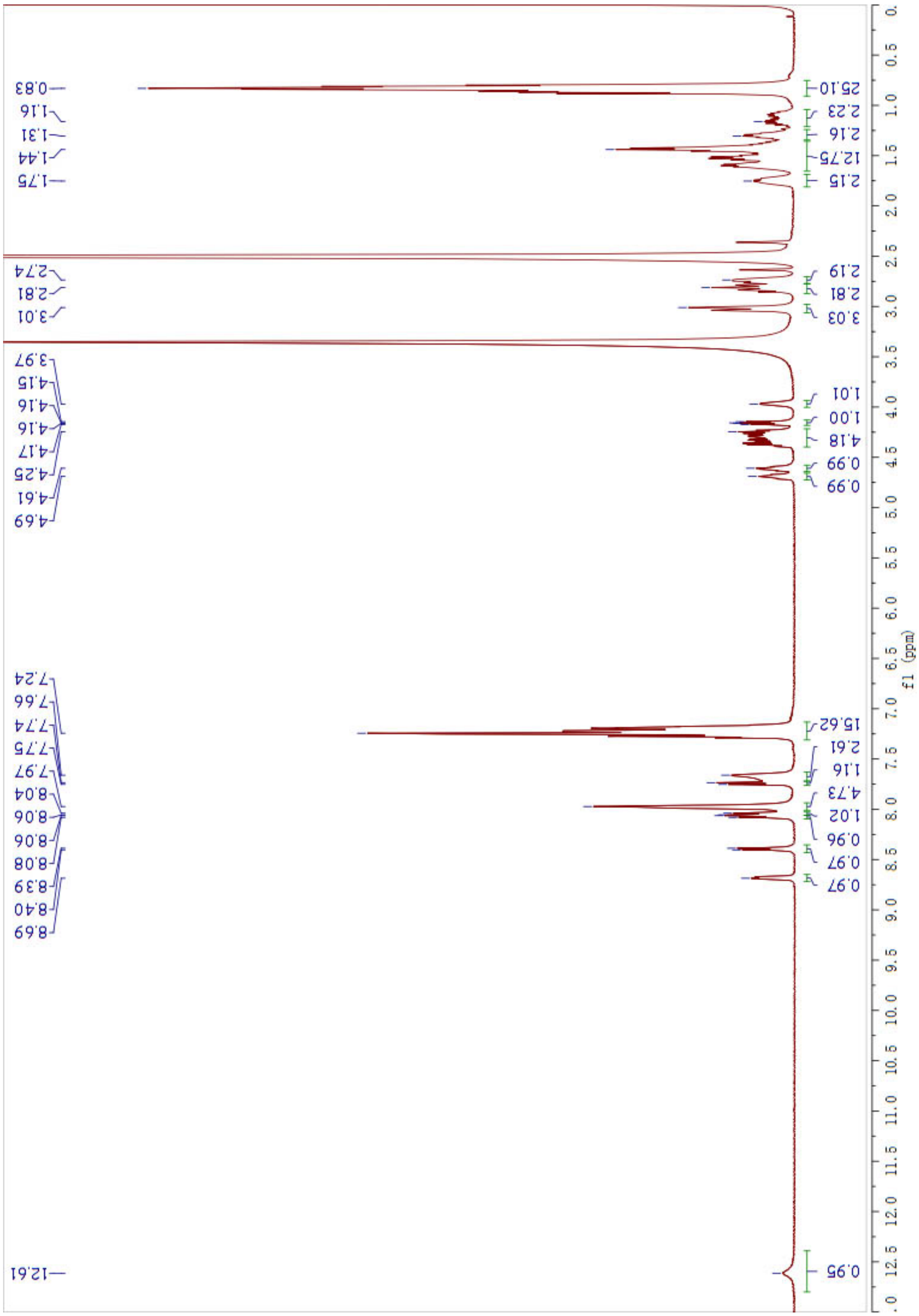


**Appendix S5.**

$^{1}$H NMR of FFFIKLLI.

| | |
|---|---|
| Lot No. | :P191203-YS766468 |
| Column | :Diamonsil C18, 4.6*250mm, 5um |
| Solvent A | :0.1%Trifluoroacetic in 100% Acetonitrile |
| Solvent B | :0.1%Trifluoroacetic in 100% Water |
| Gradient | : A B |
| | 0.0min 35% 65% |
| | 25.0min 60% 40% |
| | 25.1min 100% 0% |
| | 30.0min Stop |
| Flow rate | :1.0ml/min |
| Wavelength | :220nm |
| Volume | :20ul |

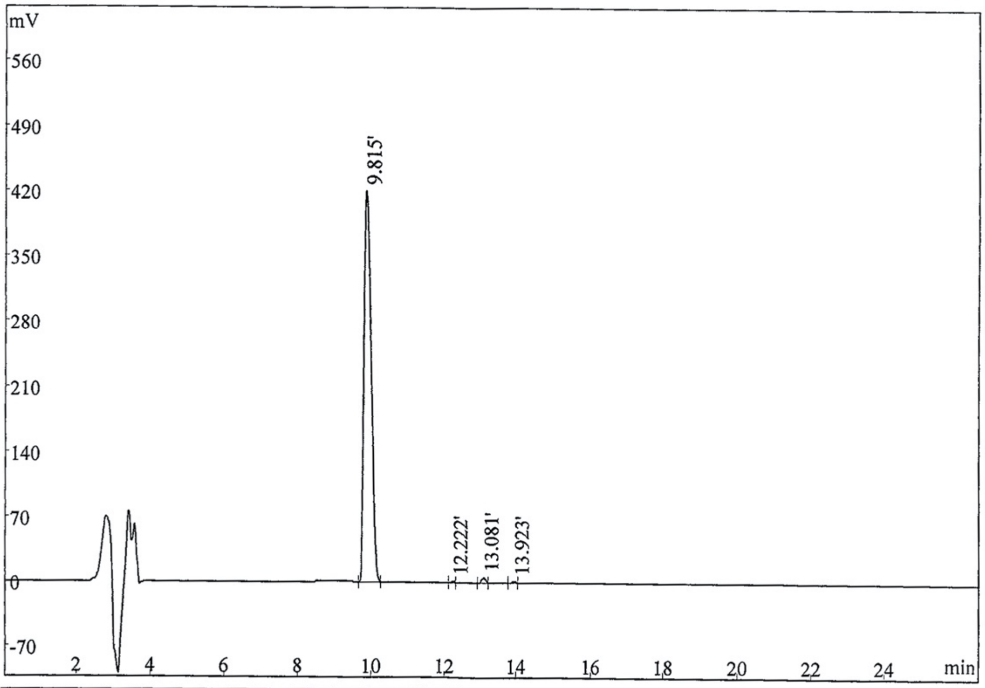


| Rank | Time | Quantity | Area | Height |
|---|---|---|---|---|
| 1 | 9.815 | 98.72 | 5120495 | 418503 |
| 2 | 12.222 | 0.2486 | 12896 | 1939 |
| 3 | 13.081 | 0.7815 | 40537 | 5163 |
| 4 | 13.923 | 0.2506 | 12999 | 1792 |
| Total | | 100 | 5186927 | 427397 |

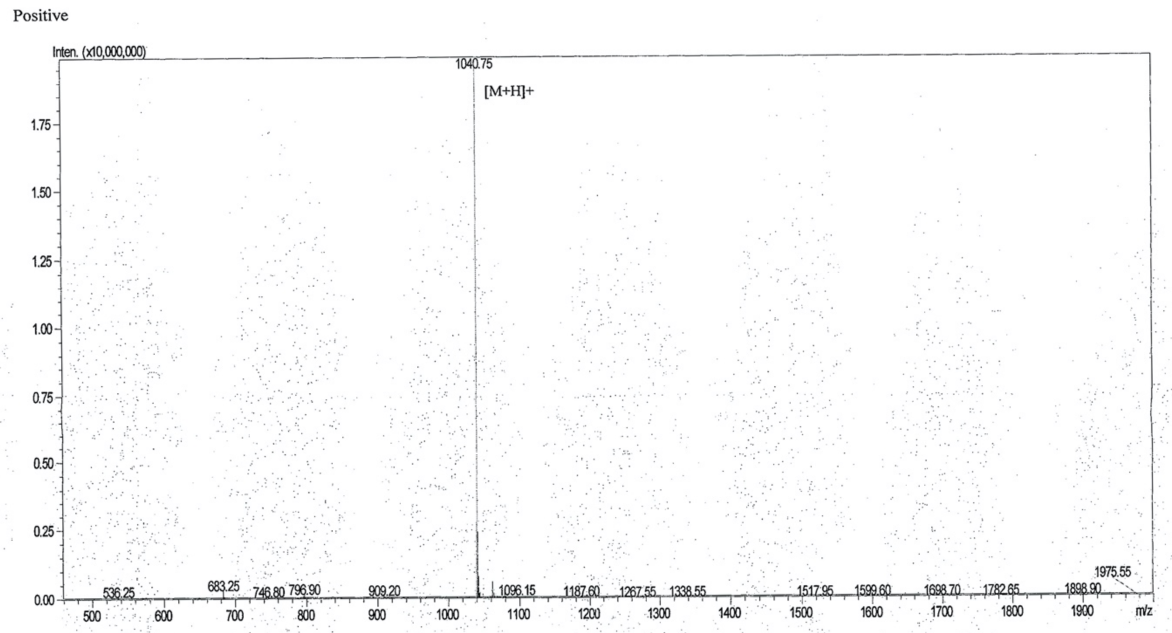


| | | | |
|---|---|---|---|
| Mw | : 1040.34 | | |
| Lot No. | : P191203-YS766468 | | |
| Probe | :ESI | Probe bias | :+4.5kv |
| Nebulizer Gas Floe | :1.5L/min | Detector | :1.2kv |
| CDL | :-20.0v | T.Flow | :0.2ml/min |
| CDL Temp | :250℃ | B.conc | :50%H2O/50%ACN |
| Block Temp | :400℃ | | |

**Appendix S6.**

LC-MS spectra of FFFKLIIL.

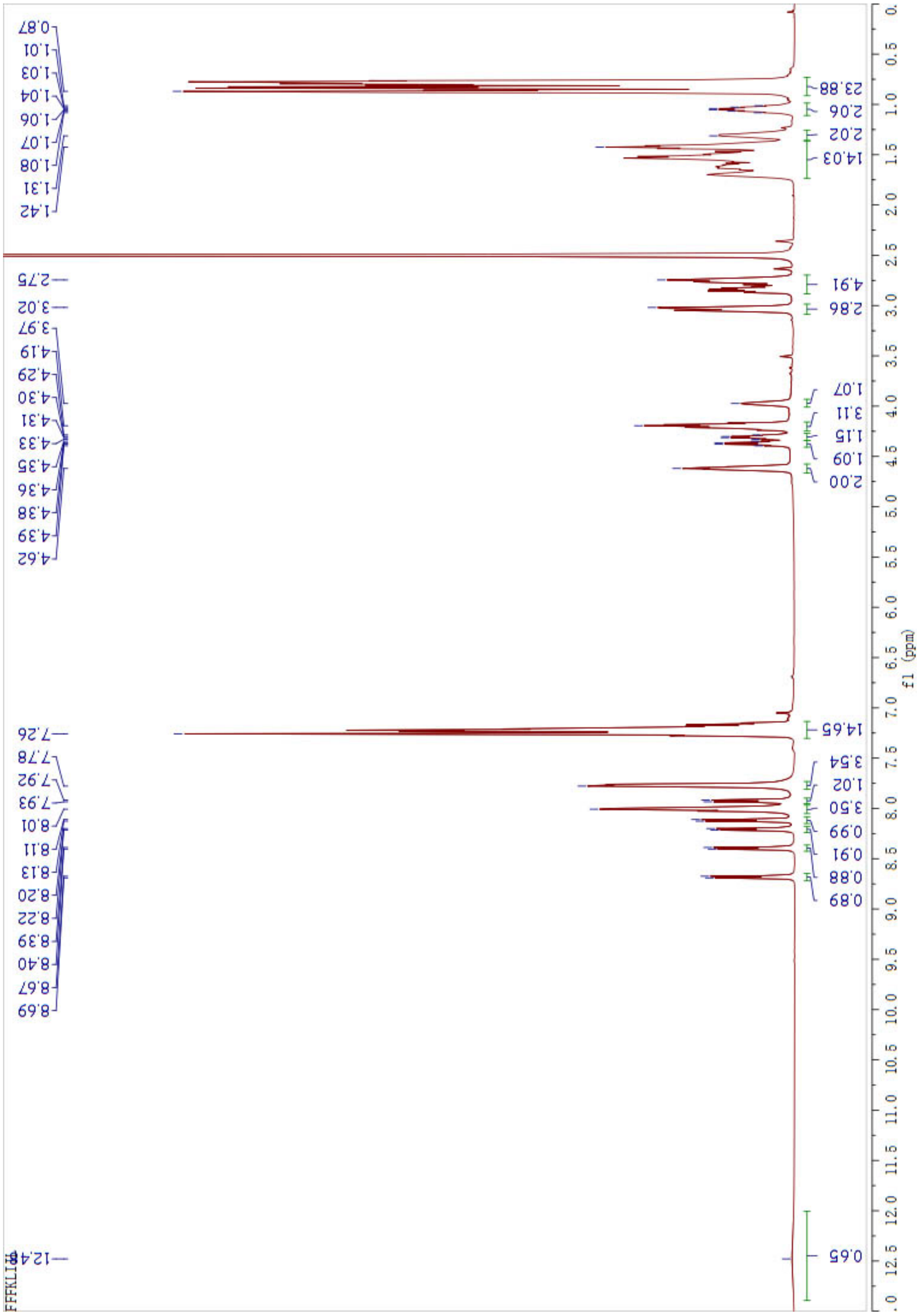


**Appendix S7.**

$^{1}H$ NMR of FFFKLIIL.

Lot No ;P201123-YW848469

Column :4.6*250mm,kromasil C18-5

Solvent A :0.1%Trifluoroacetic in 100% Acetonitrile

Solvent B :0.1%Trifluoroacetic in 100% Water

Gradient :

| | A | B |
|---|---|---|
| 0.01min | 25% | 75% |
| 25min | 50% | 50% |
| 25.1min | 100% | 0% |
| 30min | Stop | |

Flow rate :1.0ml/min

Wavelength :220nm

Volume:5ul

File opened: F:848469-25-50.hw, where

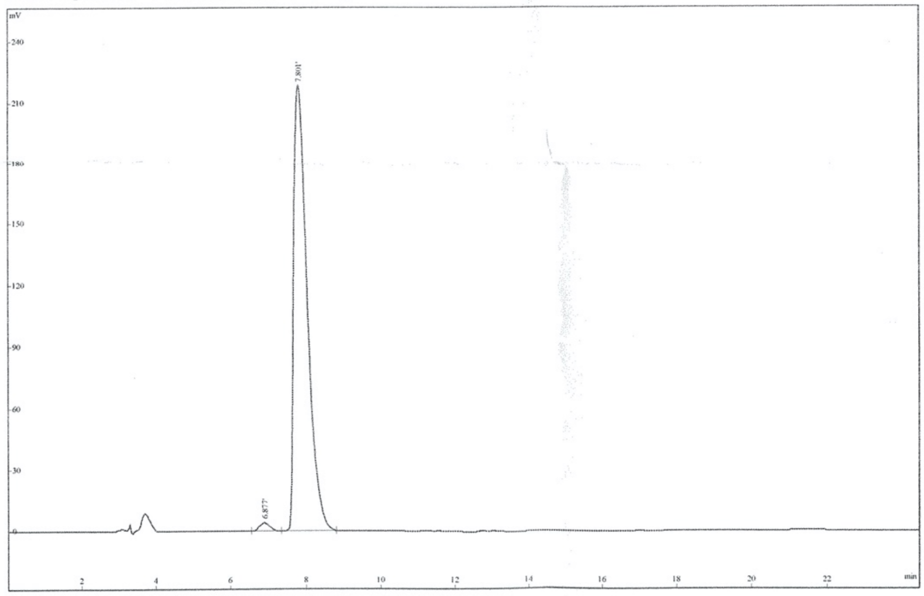

| Rank | Time | Conc. | Area | Height |
|---|---|---|---|---|
| 1 | 6.877 | 1.467 | 81003 | 4093 |
| 2 | 7.801 | 98.54 | 5441318 | 218925 |
| Total | | 100 | 5522321 | 223018 |

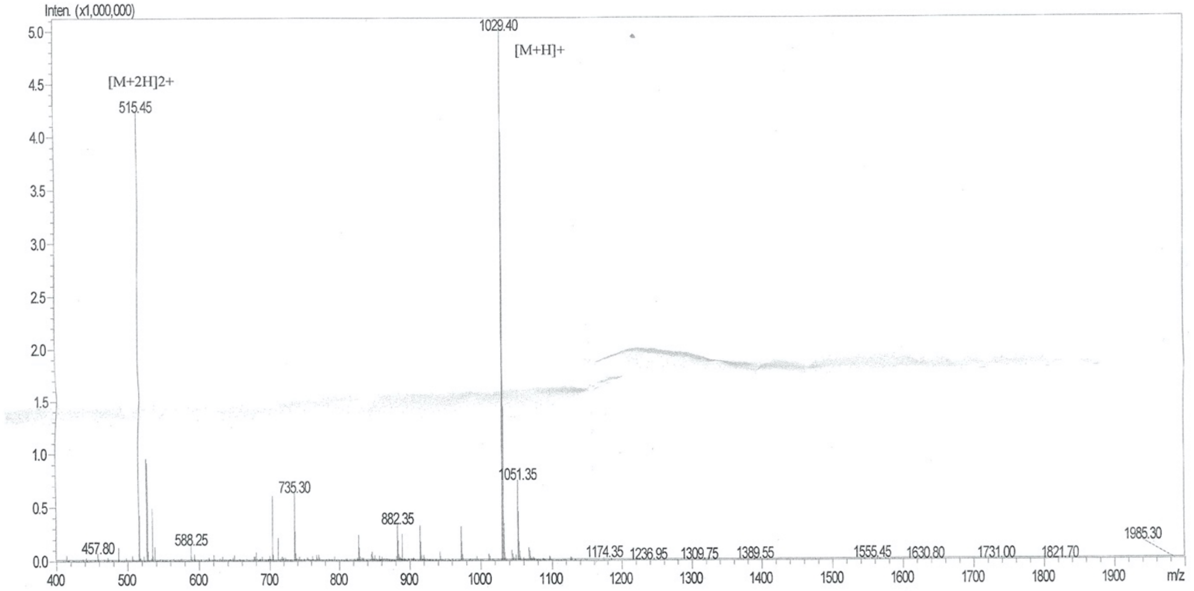


| | | | |
|---|---|---|---|
| Mw | : 1029.20 | | |
| Lot No. | : P201123-YW848469 | | |
| Probe | :ESI | Probe bias | :+4.5kv |
| Nebulizer Gas Floe | :1.5L/min | Detector | :1.2kv |
| CDL | :-20.0v | T.Flow | :0.2ml/min |
| CDL Temp | :250℃ | B.conc | :50%H2O/50%ACN |
| Block Temp | :400℃ | | |

**Appendix S8.**

LC-MS spectra of FFFGRGDSP.

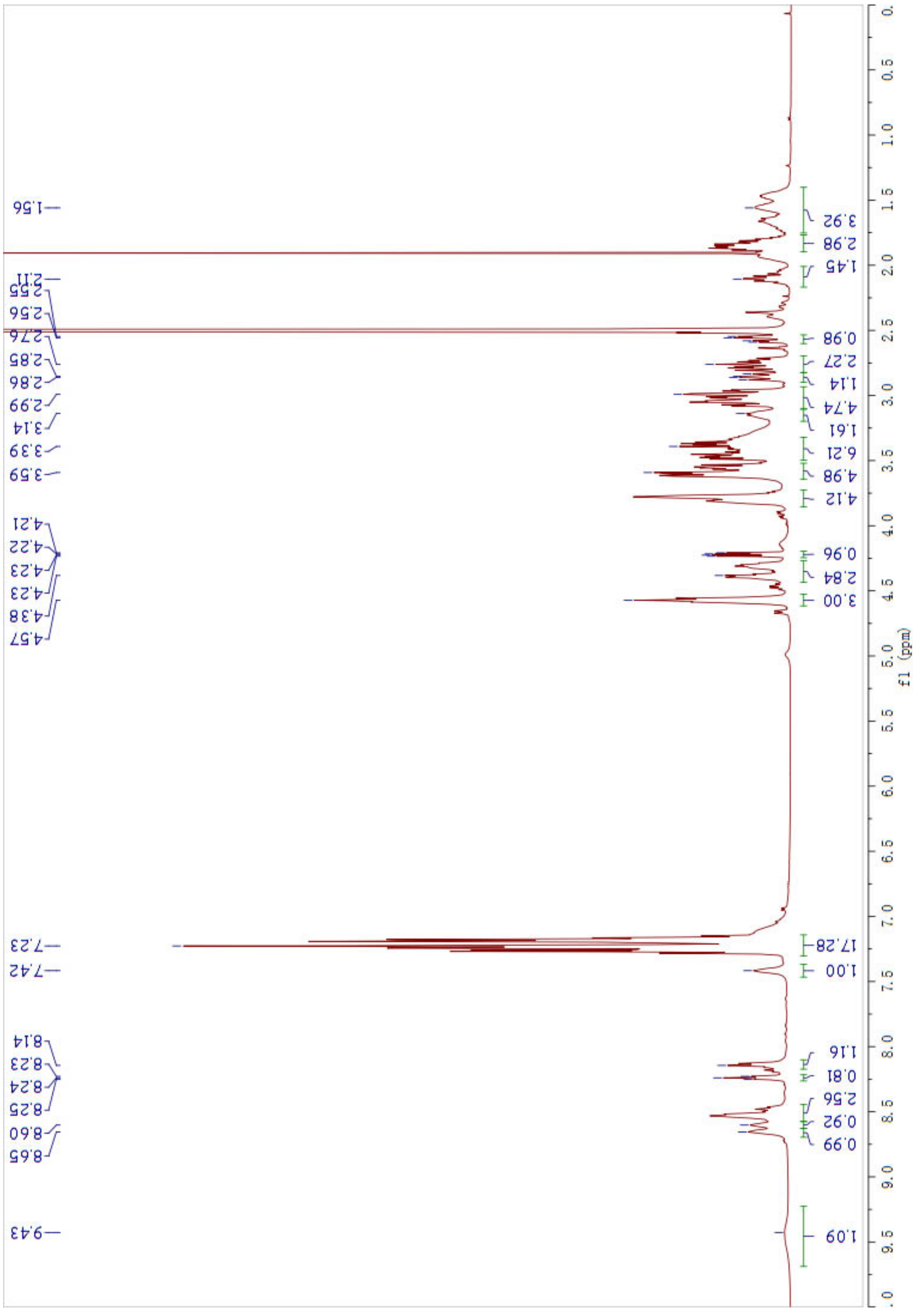

**Appendix S9.**

$^1$H NMR of FFFGRGDSP.

Lot No. : P201013-YW837788

Column : 4.6*250mm, 5um, 100A, Agela

Solvent A : 0.1% trifluoroacetic in 100% acetonitrile

Solvent B : 0.1% trifluoroacetic in 100% water

Gradient

| | A | B |
|---|---|---|
| 0.01min | 26% | 74% |
| 25min | 51% | 49% |
| 25.1min | 100% | 0% |
| 30min | STOP | |

Flow rate : 1.0ml/min

Wavelength : 220nm

Volume :

10ul

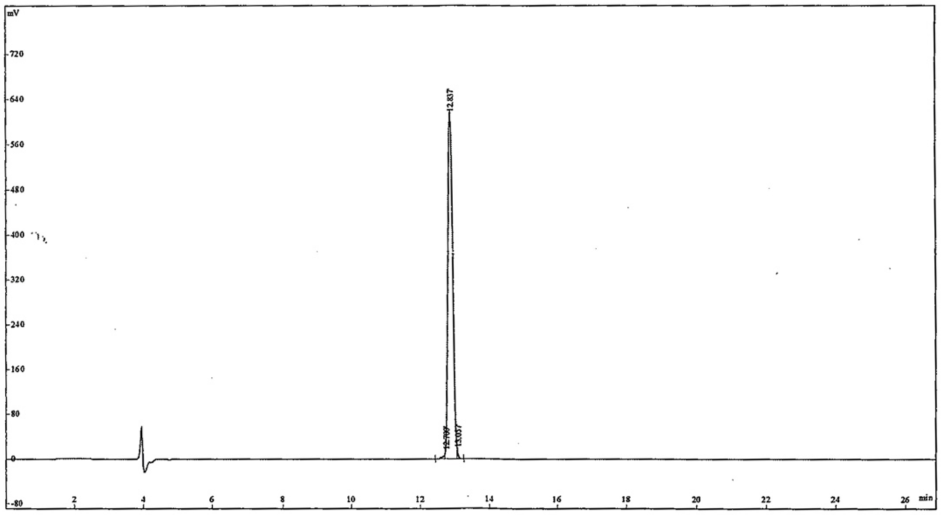


| Peak No. | Ret Time | Conc | Area | Height |
|---|---|---|---|---|
| 1 | 12.700 | 0.9246 | 57443 | 15317 |
| 2 | 12.837 | 98.73 | 6133771 | 623094 |
| 3 | 13.057 | 0.3464 | 21519 | 10059 |
| Total | | 100 | 6212733 | 648470 |

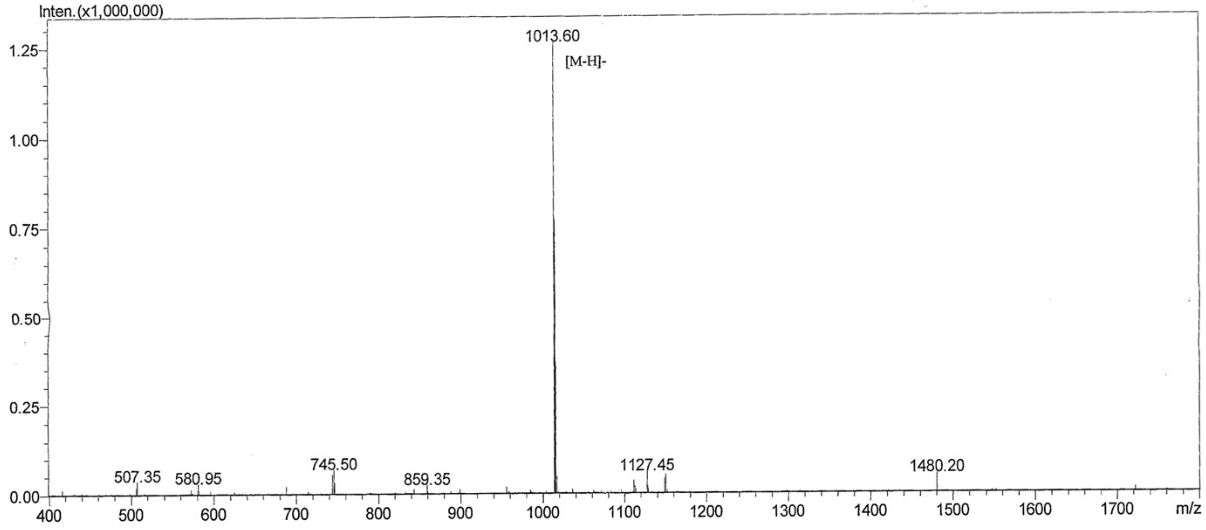


| | | | |
|---|---|---|---|
| M.W.: | 1015.21 | | |
| Lot. No.: | P201013-YW837788 | | |
| Instrument | SHIMADZU LCMS-2010EV | | |
| Probe: | ESI | Probe Bias: | +4.5kv |
| Nebulizer Gas Flow: | 1.5L/min | Detector: | 1.5kv |
| CDL: | -20.0v | T. Flow: | 0.2ml/min |
| CDL Temp.: | 250 °C | B. Conc.: | 50%H2O/50%ACN |
| Block Temp.: | 200 °C | | |

**Appendix S10.**

LC-MS spectra of FFFLRGDN.

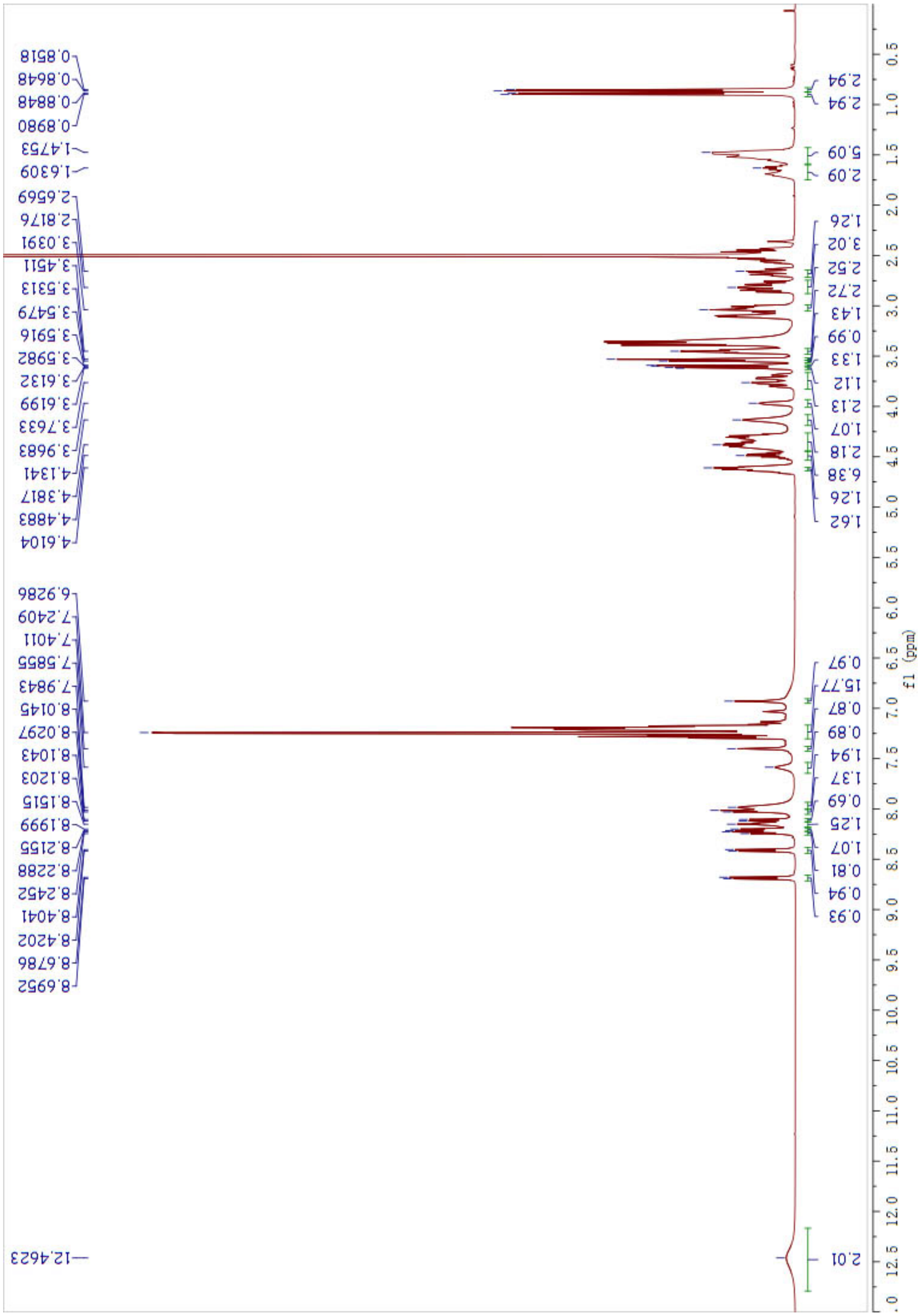

**Appendix S11.**
$^1$H NMR of FFFLRGDN.

Lot No :P190510-YS725632
Number :0200049
Column :250*4.6mm,Kromasil-C18-5um
Solvent A:0.1%TFA in 100%water
Solvent B:0.1%TFA in 100%acetonitrile

| Gradient : | A | B |
|---|---|---|
| 0.1min | 72% | 28% |
| 25.0min | 47% | 53% |
| 25.1min | 0% | 100% |
| 30.0min | stop | |

Flow rate:1.0ml/min
Wavelength(nm):220
Volume:5ul

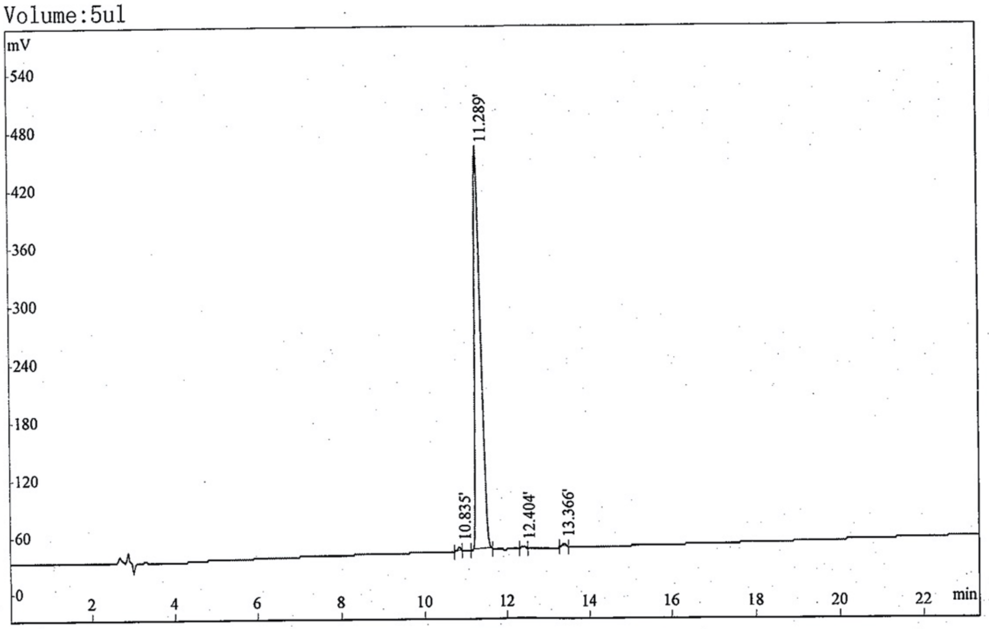


| Rank | Time | Conc. | Area | Height |
|---|---|---|---|---|
| 1 | 10.835 | 0.4214 | 18034 | 3546 |
| 2 | 11.289 | 98.7 | 4224327 | 421554 |
| 3 | 12.404 | 0.3394 | 14527 | 2434 |
| 4 | 13.366 | 0.5327 | 22798 | 3146 |
| Total | | 100 | 4279686 | 430680 |

Positive

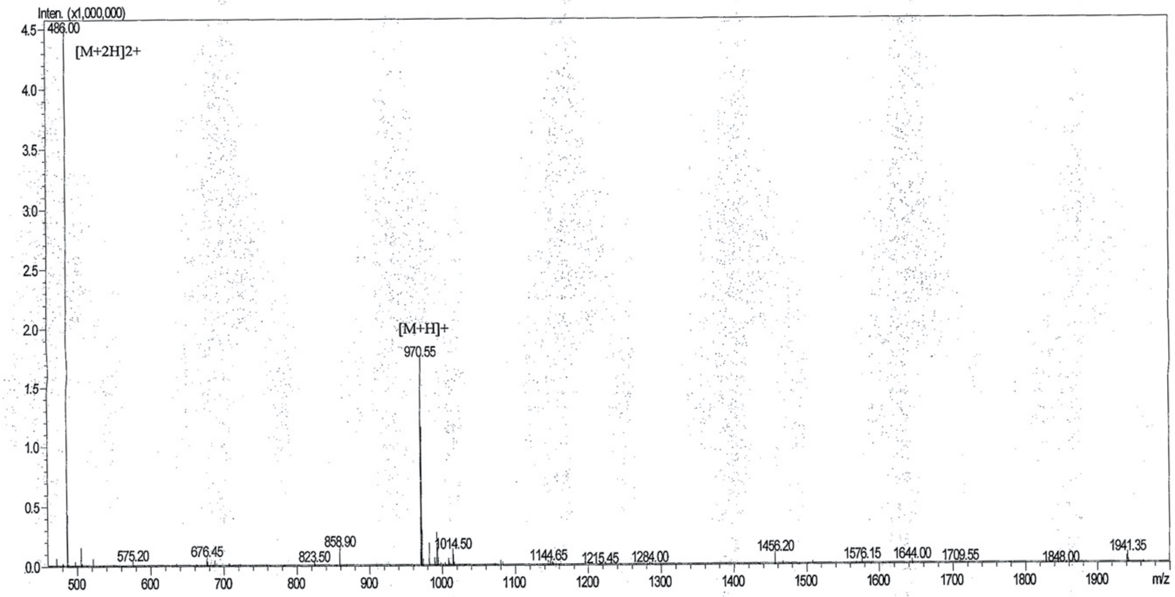


Mw : 970.21
Lot No. : P190510-YS725632

Probe :ESI Probe bias :+4.5kv
Nebulizer Gas Floe :1.5L/min Detector :1.2kv
CDL :-20.0v T.Flow :0.2ml/min
CDL Temp :250℃ B.conc :50%H2O/50%ACN
Block Temp :400℃

**Appendix S12.**

LC-MS spectra of FFFIKVAV.

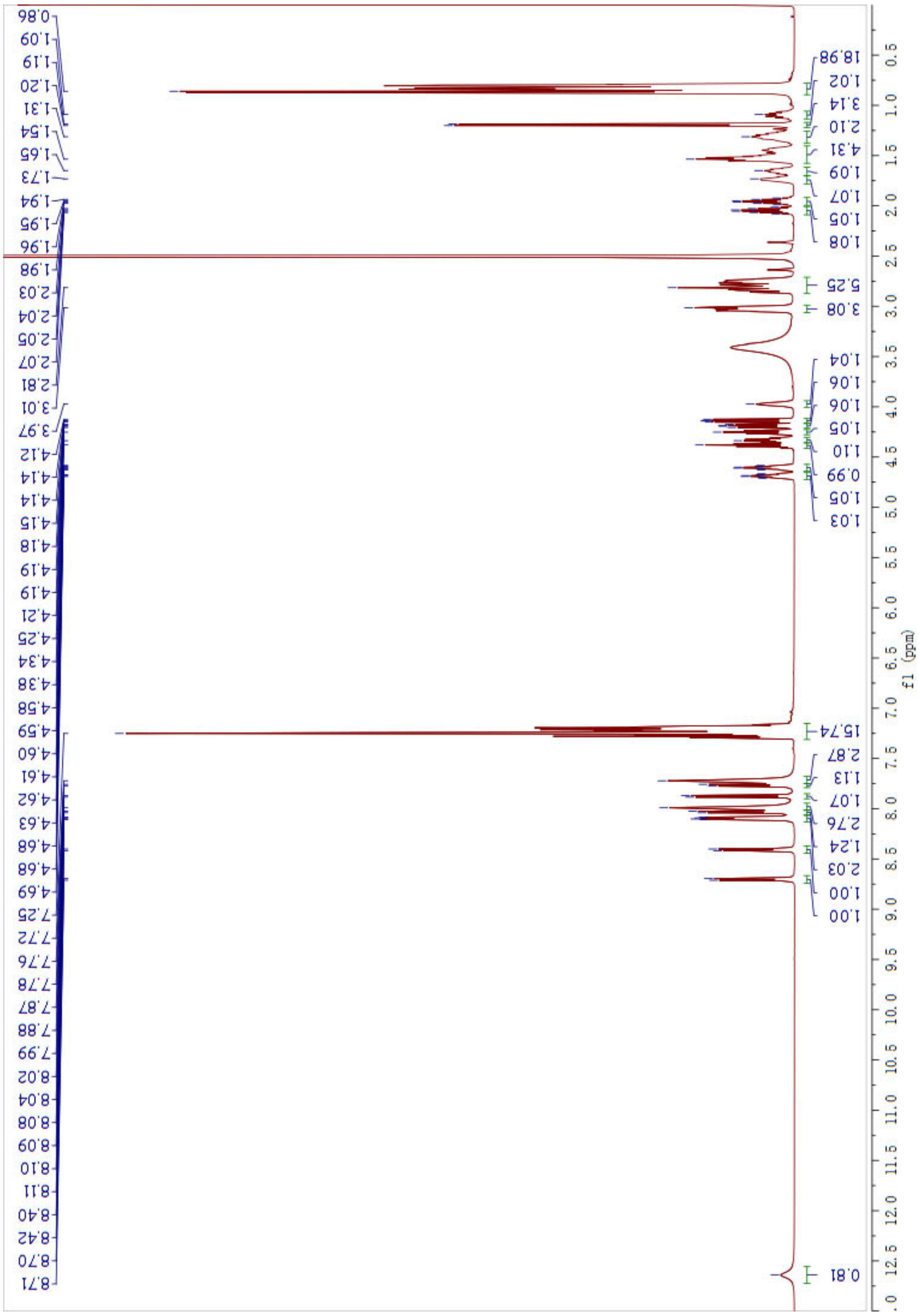

**Appendix S13.**

$^{1}H$ NMR of FFFIKVAV
